# General-purpose LLMs outperform single-cell language models at cell type annotation when expected types are known

**DOI:** 10.64898/2026.11.27.769702

**Authors:** Niklas Brunn, Adrian Fritz, Jonatan Menger, Anup Kumar, Sonia Maria Krißmer, Pavankumar Videm, Ryan Köksal, Laia Canal Guitart, Lara Gießler, Maren Hackenberg, Harald Binder, Rolf Backofen

## Abstract

Cell type annotation is the first interpretive step in almost every single-cell RNA sequencing study. General-purpose large language models (LLMs) can assign cell type labels to single cells in a zero-shot setting, without task-specific fine-tuning. In contrast, domain-adapted single-cell language models (scLMs) are trained on gene expression paired with text. Benchmarks comparing them cover only a subset of scLMs and provide models with different contextual information, such as a list of expected cell types, leaving the effect of context on annotation performance unclear. We built a benchmarking pipeline that covers a variety of scLMs, LLMs, and established marker- and reference-based baselines, and scores predicted labels against the Cell Ontology. We evaluated each method with and without a list of expected cell types. With the list, LLMs were most accurate on average. Without it, their advantage shrank, and neither LLMs nor scLMs consistently outperformed the baselines. LLMs and embedding-based scLMs benefited strongly from the list, whereas generative scLMs were barely affected. Ablations of the provided context further showed that the list mattered most and that omitting types from the list hurt more than adding types absent from the data. Overall, rare cell types were the hardest to annotate. The baselines often lacked labels for them, while LLMs achieved the highest accuracy for rare types, although accuracy remained lower than for common types. Our results suggest that LLMs are the best choice among the evaluated methods for zero-shot annotation when a comprehensive list of expected types is available.

## Introduction

Cell type annotation is a fundamental step in single-cell RNA sequencing (scRNA-seq) studies, and many downstream comparisons, from differential expression to atlas integration, inherit the labels assigned at this stage [1]. It is usually performed by domain experts with established marker- and reference-based methods, which score known marker genes or transfer labels from annotated reference data [2–6] and have been benchmarked against published annotations for years [7, 8]. Each of these methods can only name the cell types in its marker database or reference. Consequently, the cells of a missing cell type often receive another label. Language models offer a complementary route that requires no annotated reference, because biomedical text already records the marker genes of cell types.

General-purpose large language models (LLMs) are trained on large text corpora and can annotate clusters of cells from their marker genes in a zero-shot setting, without task-specific fine-tuning [9, 10]. They have also been applied to single cells, which are given as their most highly expressed genes [11–13]. Domain-adapted single-cell language models (scLMs) instead aim to combine the knowledge recorded in text with patterns learned directly from expression data. We use this term for models that build on a pretrained language model and are trained on gene expression paired with text. Generative scLMs such as C2S-Scale and InstructCell write the label as free text [14, 15]. Embedding-based scLMs such as CellWhisperer, LangCell and mmContext align expression and text in a shared embedding space and return the best-matching label from a set of candidate cell types [16–18]. Thus, these methods differ in how their label space is defined, and constraining it with a list of candidate cell types in a dataset works differently for each. Such a list reflects the cell types that a researcher expects in a sample, but earlier studies mostly provided exactly the cell types annotated in the data [10, 11, 19]. The list can guide annotation methods that write free text, defines the candidates of an embedding-based scLM, and restricts marker- or reference-based methods to the listed cell types.

Earlier benchmarks compared these methods only in part. LLMs were benchmarked mostly per cluster and without scLMs [9, 10]. For example, Xiao et al. and Cheng et al. compared many LLMs with each other and with established methods by giving each LLM the marker genes of a cluster and scoring its answers against the published annotations on the Cell Ontology [20, 21]. Only a few benchmarks included scLMs, and these tested mainly small Cell2Sentence models [14, 22] with up to 1B parameters, each with different context. SOAR used prompts with the top marker genes of each cell type [23], CellVerse offered the cell types of the respective dataset as multiple choice for each cell [11], and SC-Arena asked about single cells without any candidates [12]. In SOAR and CellVerse, the best LLMs outperformed the Cell2Sentence models, whereas in SC-Arena a Cell2Sentence model ranked highest. Giving LLMs the expected cell types improved their answers in one study [19] but not in another [10], and a broad catalog of cell types helped in a third [13]. In practice, however, the expected types are rarely known exactly [19]. None of these benchmarks compared LLMs with embedding-based scLMs or with C2S-Scale models of 2B or more parameters, ran every kind of method both with and without the list of expected types, or tested lists with controlled errors. It therefore remains unclear to what extent the provided context, particularly the list of expected cell types, influences the annotation accuracy of different model types.

To this end, we built a benchmarking pipeline in Nextflow [24] that compares six scLMs with 11 LLMs and with four established marker- and reference-based methods as baselines on the same cells of four datasets (Figure 1A,B). The datasets come from different human and mouse tissues and differ in their number of cell types. We let the LLMs annotate each cell on its own from a ranked list of its genes (Figure 1C). Every method runs unconstrained and constrained by the list of expected cell types, which holds exactly the cell types of the dataset. The baselines use marker databases and references chosen for the tissue of each dataset, and the references come from other studies, apart from a small overlap in the pancreas data. Because the label spaces of the methods differ, we map all labels to the Cell Ontology [25, 26], where different names of the same cell type share one term and answers naming a related type can receive partial credit (Figure 1D,E).

**Figure 1.**
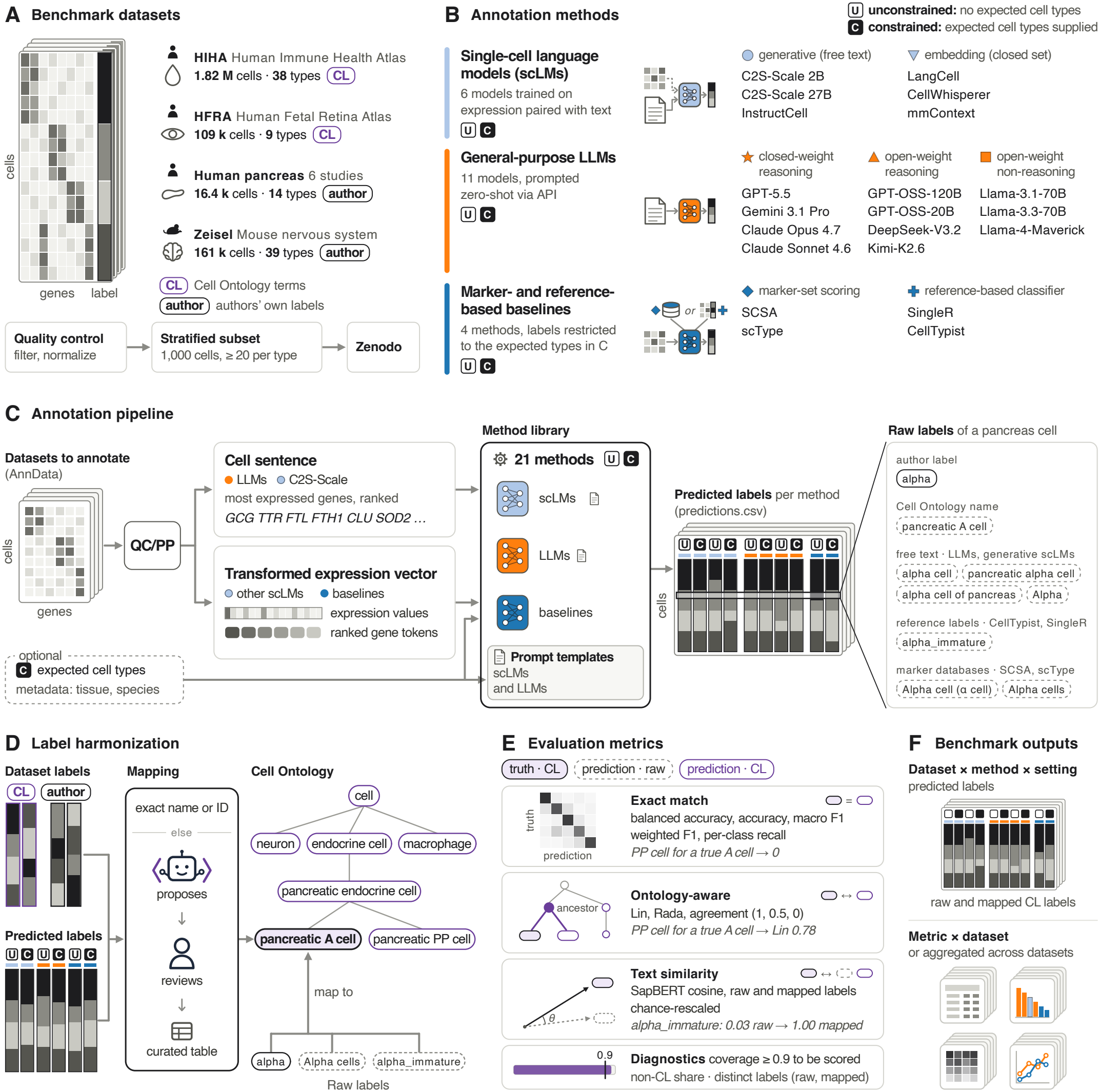
Benchmark overview. (**A**) The four benchmark scRNA-seq datasets, with their number of cells and cell types as downloaded. CL marks Cell Ontology terms and author the authors’ own labels. After quality control and preprocessing (QC/PP), a test set of 1,000 cells with at least 20 cells per cell type is drawn and stored on Zenodo for the benchmark. (**B**) The 21 annotation methods. scLMs are trained on gene expression paired with text. They write the label as free text (generative) or pick the best-matching candidate (embedding-based). General-purpose large language models (LLMs) are queried through an application programming interface (API) without examples. Marker- and reference-based baselines use known marker genes or annotated reference data. Every method runs unconstrained (U) and constrained (C) by the list of expected cell types. The list enters the prompt of the generative scLMs and LLMs, replaces all Cell Ontology names as the candidates of the embedding-based scLMs, and restricts the baselines to labels that are a listed type or a subtype of one. (**C**) The LLMs and C2S-Scale read each cell as the names of its most highly expressed genes in descending order. The other methods read its transformed expression values. Optional inputs are the list of expected cell types and metadata such as tissue and species. In the end, each method returns one label per cell. The output labels of methods differ, as shown for pancreatic alpha cells. (**D**) The labels of the datasets and all predictions are mapped to the Cell Ontology, a hierarchy of cell types. A label that matches the name or identifier of a term maps directly. Other labels are mapped by a researcher, helped by automatic suggestions, in a curated table. (**E**) In the evaluation, exact-match metrics count only correct labels. Ontology-aware metrics give partial credit for a related cell type. Text similarity compares the meaning of a raw or mapped prediction with the true label using a biomedical language model (SapBERT). A run is scored only if at least 90% of its cells have a mapped label. (**F**) The pipeline returns the raw and mapped labels of every dataset, method and setting, and tables and plots of every metric per dataset and across datasets.

The results of our benchmark indicate that the advantage of LLMs depends on the list of expected cell types. Constrained by the list, LLMs were the most accurate on average. Without the list, the closed-weight LLMs remained the most accurate on average but did not outperform the baselines on every dataset, and the small C2S-Scale 2B matched the best open-weight LLMs. Embedding-based scLMs strongly depended on the list of expected types, whereas generative scLMs were barely affected by it. The curated mapping to the Cell Ontology mainly affected the baselines without the list and had little effect on most language models. A prompt-context ablation showed that providing the list helped LLMs more than tissue and species information, and that annotating single cells was on average more accurate than annotating clusters but came at a higher cost. We also tested lists that contained errors, which is a more realistic scenario. LLMs, embedding-based scLMs and the baselines then lost most or all of the benefit of the list, whereas C2S-Scale 2B remained stable. Cell types missing from the list reduced accuracy more than additional types that were not in the data. Finally, cell types that are rare in the datasets were often missing from the baselines’ label spaces, and LLMs annotated them best but still less accurately than common types. The benchmarking pipeline, the code for all analyses and the test sets used in the benchmark are openly available.

## Materials and methods

### Problem formulation

Our aim is to compare annotation methods that differ in the labels they can return and to measure how constraining them by a list of expected cell types changes their annotation performance. A test set consists of *n* cells with expression matrix **X** and true labels *y*_1_, …, *y*_*n*_ taken from the published annotation of the dataset. An annotation method *f* returns one label per cell,

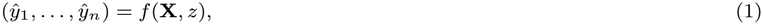

where the context *z* is any information the method receives besides the expression data. The set of labels a method can return, its label space, depends on the kind of method,

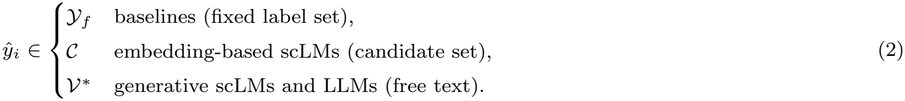

Here *Y*_*f*_ contains the labels of a baseline’s marker database, reference data or trained model, *C* is a set of candidate names supplied at inference, and *V*^*^ is the set of all finite strings over a model’s vocabulary.

To compare labels from different label spaces, we map every label to a term of the Cell Ontology with a map *φ* (Ontology mapping) and count an answer as correct if *φ*(*ŷ*_*i*_) = *φ*(*y*_*i*_). A method can thus label cell *i* correctly only if its label space contains a label that maps to *φ*(*y*_*i*_), which is always true for free text but not for a fixed label set or a candidate set.

The context *z* may contain the list of expected cell types, which we define as the set of true labels of the test set,

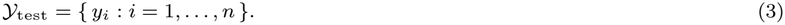

We run every method in two settings, constrained, in which it receives *Y*_test_, and unconstrained, in which no list is provided. Constrained by the list, a baseline keeps only those labels in *Y*_*f*_ that map to a listed type or to one of its subtypes and returns the listed type. For an embedding-based scLM the list becomes the candidate set, *C* = *Y*_test_, which otherwise holds the names of 3,169 Cell Ontology terms. A generative model is asked to choose from the list but can still write any string. Within each of the two settings, all methods annotate the same cells, so we can compare their performance directly.

### Datasets and test sets

We used four public scRNA-seq datasets of human and mouse tissues (Figure 1A and Supplementary Table S1). The Human Fetal Retina Atlas (HFRA) [27], which combines two studies [28, 29], and the Human Immune Health Atlas (HIHA) of peripheral blood [30] were downloaded from release 2025-11-08 of the CELLxGENE Census [31] and are labeled with Cell Ontology terms. The Zeisel data of the mouse nervous system [32] carry the authors’ labels, and the human pancreas data combine six studies [33–38] with labels harmonized for a benchmark of data integration [39]. We semi-automatically mapped the labels of these two datasets to Cell Ontology terms, which merged some of them (Ontology mapping).

Each dataset was preprocessed from its raw counts with Scanpy [40] (Supplementary Note 1). We first removed low-quality cells based on median absolute deviations (MADs) [1]. A cell was removed if its total counts, its number of detected genes or the share of its counts in the 20 most expressed genes lay more than five MADs from the median of the dataset. A cell was also removed if its share of mitochondrial counts lay more than three MADs from the median or exceeded 8%. We then discarded cells with fewer than 200 detected genes and genes detected in fewer than 10 cells. Finally, the counts of each cell were divided by the total counts of that cell, multiplied by 10,000 and log-transformed with a pseudocount of one.

For our benchmark, we drew a test set of 1,000 cells (999 in three datasets after rounding) from each preprocessed dataset. Every cell type with at least 20 cells is represented in proportion to its share of the dataset, but by at least 20 cells drawn at random. This resulted in 9 cell types in HFRA, 38 in HIHA, 29 in Zeisel and 11 in the pancreas data. A second test set of 10,000 cells per dataset, drawn in the same way, was annotated by all methods except the LLMs and unconstrained LangCell, due to limited resources and long runtime, respectively. All test sets are available on Zenodo (Data availability).

### Annotation methods and their inputs

We compared 21 methods from three families (Figure 1B and Supplementary Table S2). The four baselines are the marker-based scType [4] and SCSA [3] and the reference-based CellTypist [6] and SingleR [5]. The six scLMs are the generative C2S-Scale 2B and 27B [14] and InstructCell [15] and the embedding-based CellWhisperer [16], LangCell [17] and mmContext [18]. We included scLMs with public weights [41–46] whose authors describe their use for cell type annotation without fine-tuning. The 11 LLMs are the closed-weight reasoning models Claude Opus 4.7 [47], Claude Sonnet 4.6 [48], Gemini 3.1 Pro Preview [49], referred to as Gemini 3.1 Pro, and GPT-5.5 [50], the open-weight reasoning models DeepSeek-V3.2 [51], GPT-OSS-120B, GPT-OSS-20B [52] and Kimi-K2.6 [53], and the open-weight models without reasoning Llama-3.1-70B [54], Llama-3.3-70B [55] and Llama-4-Maverick [56].

The annotation methods receive different representations of a cell as input (Figure 1C). The LLMs and C2S-Scale receive a cell sentence [22], the names of up to 200 of the most highly expressed genes of the cell in decreasing order of expression, with ties broken at random [14]. CellWhisperer and LangCell receive the genes of a cell ranked as in Geneformer [57], mmContext the log-normalized expression of 10,000 preselected genes, and InstructCell the raw counts of the genes in its vocabulary. Because these four models expect human genes, we replaced the genes of the mouse Zeisel data by their one-to-one human orthologs from Ensembl release 114 [58] where available.

The baselines use resources chosen to match the tissue of each dataset (Supplementary Table S3). CellTypist applies a pretrained model [6, 59–61] and SingleR an annotated reference [62–65]. scType and SCSA score marker genes, from scType’s own database and from CellMarker 2.0 [66], on Louvain clusters [67], and each test cell takes the label of its cluster. Both cluster a sample of up to about 100,000 cells of the preprocessed dataset, which contains the test sets and is also available on Zenodo (Supplementary Note 1). The SingleR references come from studies other than those of the test data, except that the pancreas reference includes cells of one of the pancreas studies [34]. This concerns 11 of the 999 pancreas test cells.

### Prompts and model access

The prompt templates for the LLMs are given in Supplementary Note 2 (see also Figure 1C). The unconstrained template asks for the canonical Cell Ontology name of the cell type, and the constrained template provides the list of expected cell types and asks the model to choose from it. The list is reshuffled for every cell to avoid position bias. The prompts of C2S-Scale and InstructCell follow their model cards [41–43] and additionally name the species, and for InstructCell also the tissue.

We queried the LLMs through the OpenRouter API (model versions and dates in Supplementary Note 1). The reasoning models ran with medium reasoning effort or, for the Anthropic and Google models, a budget of 10,000 reasoning tokens, and GPT-5.5 ran with the effort xhigh (Supplementary Table S2). We left the temperature and the output limit at the providers’ defaults, sent a seed of 1234 with every request and let OpenRouter route each request to the cheapest available provider [68]. The whole response counts as the answer. Failed requests were retried, with at most 15 requests per cell, and a cell that still had no answer counts as wrong. Because hosted models can change behind the same name, a rerun may not reproduce these answers exactly. We measured the cost of a run as its spend in OpenRouter’s activity log per 1,000 annotated cells. This cost reflects the providers that served the requests and their prices at the time of the runs.

### Ontology mapping

Predicted and true labels can name the same cell type differently, for example through synonyms or abbreviations. In addition, a prediction of a closely related cell type is less wrong than one of an unrelated type. Earlier benchmarks addressed this by string overlap [23], by the similarity of text embeddings [69] or by mapping labels to the Cell Ontology [9]. Similar to Xiao et al. [20], we map the true labels and the answers to Cell Ontology terms semi-automatically (Figure 1D).

Each answer is first stripped of surrounding whitespace, quotes and trailing periods, converted to lower case and joined into one line. It maps directly if it equals the name or identifier of a current term in the Cell Ontology release of 8 June 2026 [25, 26, 70]. Synonyms listed in the ontology are not matched automatically, because they can point to an unintended term. For example, *alpha cell* is a synonym of a retinal ganglion cell type of the mouse. Every other answer enters a curated table with a term suggested by CellOntologyMapper [71], which we reviewed manually to define the final mapping. When a model explained or corrected itself in its output, we used its final answer. An answer that clearly meant a certain cell type was mapped to that type. A general name was not made more specific than written. For instance, *acinar cells* maps to *acinar cell* and not to *pancreatic acinar cell*, even for a cell from the pancreas. Answers that name no cell type map to the root term *cell* and count as wrong.

### Evaluation metrics

We compare the answers with the true labels after mapping both to the Cell Ontology (Figure 1E). We score them with exact-match, ontology-aware and text-similarity metrics, all defined in Supplementary Table S4. We use balanced accuracy as the main metric of our benchmark,

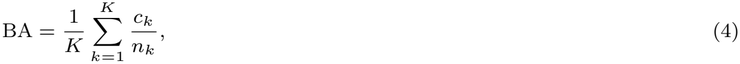

where *K* is the number of cell types in the test set, *n*_*k*_ is the number of cells of type *k* and *c*_*k*_ is the number of these cells that were labeled correctly. Rare and common cell types thus count equally. In addition, we report two ontology-aware metrics that give partial credit to related answers. For the agreement score, which we adapted from Hou and Ji [9], we assign 1 to an answer that names the true type and 0.5 to an answer that names a finer type, a subtype of the true type, or a coarser type, a type above it in the hierarchy. A coarser type earns no credit if it is broad, which we define as having more than 250 terms at or below it, such as *neuron* or *leukocyte*. Any other answer receives 0. We average these values over the cells of each type and then over the types. As the second ontology-aware metric, we use the Lin similarity [72],

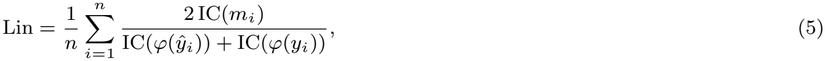

where *m*_*i*_ is the most informative common ancestor of *φ*(*ŷ*_*i*_) and *φ*(*y*_*i*_), the term with the highest information content among all terms at or above both. The information content of a term *t* is IC(*t*) = 1 − log *d*(*t*)*/* log *N*, where *d*(*t*) is the number of terms at or below *t* and *N* is the number of terms in the ontology [73]. It is 0 for the root term *cell* and 1 for a term without subtypes. With both metrics, the answer *T cell* for a *CD4-positive, alpha-beta T cell* scores higher than a cell type from another branch of the ontology.

Further, we use a text similarity metric to score the raw answers of the models, without the mapping. We embed the answer and the Cell Ontology name of the true type with SapBERT [74] and take the cosine between the two embeddings. We average it like the agreement score and rescale it as in BERTScore [75], so that two random term names score 0 on average and identical names score 1, with negative values set to 0. We report the text similarity for the raw answers and, in addition, after the mapping. We additionally report accuracy, F1 scores and a path-based similarity [76] in the result tables (Supplementary Tables S5–S8).

To characterize the answers of a method beyond right and wrong, we sort each answer into the classes of the agreement score, as the exact type, a coarser or a finer type, another cell type or no usable answer. To mark the chance level of each metric, we include a random model that assigns cell types in proportion to their frequency in the test set. The pipeline returns the raw and mapped labels of every run, together with tables and plots of every metric per dataset and across datasets (Figure 1F).

### Prompt-context ablation and imperfect-list analysis

In further experiments, we varied the context provided to selected methods (Supplementary Note 1). Every condition was run with three seeds, and we report the mean over the seeds.

In the prompt-context ablation, GPT-OSS-120B, C2S-Scale 2B and its base model Gemma-2-2B [77] annotated single cells from cell sentences, and GPT-OSS-120B and Gemini 3.1 Pro annotated clusters from their 25 top-ranked marker genes. Each of these five arms received the genes (i) alone, (ii) with the tissue and species, (iii) with the list of expected cell types or (iv) with both (Supplementary Note 2). Because the prompt of C2S-Scale always names the species, the tissue and species add only the tissue for it. Gemma-2-2B received the prompts of the LLMs, and like C2S-Scale 2B it decoded greedily [14] with at most 20 new tokens [41]. We obtained the clusters with the Leiden method [78] at resolution 1.0 from the same sample of cells as for scType and SCSA, and every test cell took the answer of its cluster.

In the imperfect-list analysis, GPT-OSS-120B per cell, Gemini 3.1 Pro per cluster, C2S-Scale 2B, mmContext, LangCell and the four baselines received a list with errors in place of the exact list. We added a parent or a child in the Cell Ontology for half of the listed types (A1), added five types absent from the data (A2), removed a quarter of the types (A3) or combined all three errors. The types to add and to remove were drawn once per dataset and seed. We took all added types from the Cell Ontology terms used in CELLxGENE. An absent type shares a parent with a listed type and is neither coarser nor finer than any cell type of the dataset. In A3 and in the combination of all three errors, the prompts of the generative models included the instruction to answer *uncertain* if none of the listed cell types fits. We scored the answer *uncertain* as wrong. To test the effect of the instruction alone, we also ran the generative models with the exact list and this instruction.

In contrast to the benchmark, we took only the first non-empty line of a response as the answer and mapped new answers to the Cell Ontology by a fixed rule instead of manual review.

### Rare cell types

We call a cell type rare if its share of the cells in the full preprocessed dataset is below 1%, which holds for 44 of the 87 cell types. For each run, we computed balanced accuracy separately over the rare and over the common types of a dataset and averaged it over the four datasets. In each run, a rare type has one of four outcomes. The run (i) labeled at least half of its cells correctly, (ii) labeled some but fewer than half correctly, (iii) labeled none correctly but gave the name of the type to other cells, or (iv) never used the name. We counted these outcomes over all 44 rare types. For the constrained baselines, we also checked whether a method could name the type at all, that is, whether one of its labels maps to the type or to one of its subtypes. We report means over the methods of a family, separately for the unconstrained and the constrained setting.

## Results

### Benchmarking annotation performance with and without the list of expected cell types

We ran each of the 21 methods (Figure 1B) unconstrained and constrained on the same 1,000 cells of every dataset (Supplementary Tables S1 and S2). Their balanced accuracy (Supplementary Table S4) differed strongly between the datasets (Figure 2C–F and Supplementary Tables S5– S8). Averaged over all runs and methods, it was 0.60 on human pancreas (range 0.06 to 0.94), 0.32 on HFRA (0.00 to 0.59), 0.25 on Zeisel (0.01 to 0.60) and 0.19 on HIHA (0.02 to 0.40). Despite these differences, an LLM scored highest with the list of expected cell types on every dataset, whereas without the list the highest-scoring method was a baseline on HFRA and an scLM on HIHA (Figure 2C–F).

**Figure 2.**
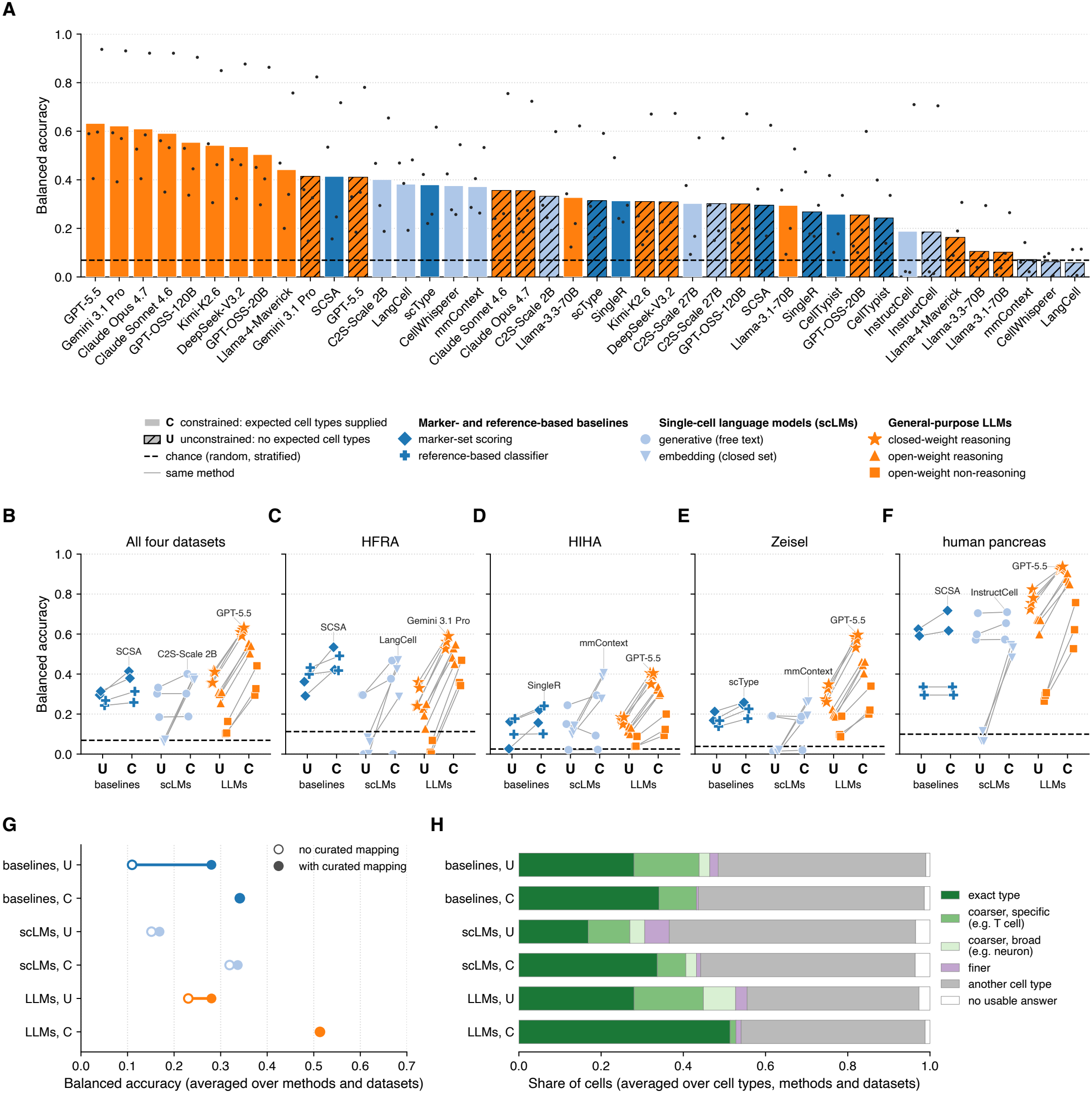
Benchmark results. (**A**) Balanced accuracy of every method in both settings, averaged over the four datasets at 1,000 cells each, from best to worst. Balanced accuracy is the share of correctly labeled cells, averaged over cell types. The dots on each bar are the four datasets, from left to right HFRA, HIHA, Zeisel and human pancreas. Solid bars are runs constrained by the list of expected cell types (C), and hatched bars are unconstrained runs (U). (**B**) Balanced accuracy of each method, grouped by family and setting, averaged over the four datasets, and (**C**–**F**) on each dataset. Grey lines join the unconstrained and constrained runs of the same method. Symbols give the subcategory, as in Figure 1B. The best constrained run of each family is named. (**G**) Balanced accuracy when only the answers that need no curated mapping count as correct, meaning Cell Ontology terms or the dataset’s own labels as written (open circles), and with the curated mapping of the other answers (filled circles). Each point is the mean over the methods of one family and setting and over the four datasets. Where only one circle is visible, the two overlap. Supplementary Figure S1B gives the values per method. (**H**) Where each family’s answers fall in the Cell Ontology relative to the true cell type, in the rows of G. An answer is the exact type, a coarser or a finer type, another cell type, or no usable answer. Coarser types are split into specific ones and broad ones, which cover more than 250 cell types in the ontology. Shares are averaged over cell types, as in balanced accuracy, and then over methods and datasets as in G. Supplementary Figure S2 shows them per dataset and per subcategory. In A to F, the dashed line is a random model that assigns cell types in proportion to their frequency. HFRA, Human Fetal Retina Atlas; HIHA, Human Immune Health Atlas; LLMs, general-purpose large language models; scLMs, single-cell language models. Averages over datasets use *n* = 4 datasets.

Averaged over the datasets, the LLMs scored 0.51 with the list, against 0.34 for both the scLMs and the baselines (Figure 2B). The best LLM, GPT-5.5, reached 0.63, closely followed by Gemini 3.1 Pro at 0.62 and Claude Opus 4.7 at 0.61 (Figure 2A). Nine of the 11 LLMs ranked above every scLM and baseline. Only Llama-3.3-70B and Llama-3.1-70B fell below the best baseline, SCSA, and the best scLM, C2S-Scale 2B, which scored 0.41 and 0.40. On every dataset, the best LLM scored higher than the best baseline, and on three datasets also higher than the best scLM (Figure 2C–F). On HIHA, mmContext matched GPT-5.5, both at 0.40.

Without the list, the advantage of the LLMs in balanced accuracy shrank. The four closed-weight LLMs remained the most accurate on average, but the 11 LLMs and the four baselines both averaged 0.28 (Figure 2A,B). Neither the best LLM nor the best scLM scored higher than the best baseline on every dataset, and both scored lower than SingleR on HFRA (Figure 2C–F). The small C2S-Scale 2B ranked fifth, behind only the four closed-weight LLMs and just above every open-weight LLM and every baseline (Figure 2A). On HIHA it scored higher than every LLM, 0.24 against at most 0.19.

The effect of adding the list differed between the kinds of methods (Figure 2B). On every dataset, the list raised the balanced accuracy of every LLM, by 0.06 to 0.45, and of every embedding-based scLM, by 0.18 to 0.48 (Supplementary Tables S5–S8). In the unconstrained setting, the embedding-based scLMs chose among all 3,169 Cell Ontology names and scored about as well as the random model while the generative scLMs were barely affected by the list. Only C2S-Scale 2B improved, while C2S-Scale 27B and InstructCell did not change on average. On HIHA, the list even lowered the balanced accuracy of C2S-Scale 27B, from 0.15 to 0.09. All three generative scLMs often did not follow the instruction to pick from the list, naming a cell type not on it for 27 to 41% of the cells, whereas the LLMs almost never did (Supplementary Table S9). Finally, the baselines improved only modestly when restricted to the listed types.

For a fair comparison, we mapped all labels to the Cell Ontology, as earlier benchmarks did [20, 21]. Overall, the methods depended unequally on the curation (Figure 2G, Supplementary Figure S1 and Methods). Baselines kept less than half of their accuracy when only answers that needed no curated mapping counted. For example, no label of scType matched a Cell Ontology term or a label of the dataset as written, and without the mapping it would have scored zero. The language models depended much less on the mapping.

The list changed the kind of answers the LLMs gave (Figure 2H and Supplementary Figure S2). Without the list, a quarter of their answers named a coarser type than the true one. With the list, coarser answers almost disappeared, and the share of exact answers rose from about a quarter to about half. Answers that named another cell type stayed about as frequent and remained the most common error in every model family. The ontology-aware metrics (Figure 1E), which give credit to related types, ranked the methods similar to balanced accuracy, as do metrics based on text similarity that needs no curated mapping (Supplementary Figure S3). We additionally evaluated a subset of methods on 10,000 cells per dataset. Aggregate performance was lower in these experiments because particularly the highest-performing LLMs were evaluated only on the 1,000-cell subsets. When the comparison was restricted to models run at both scales, performance did not change much with the larger sample, with mean weighted F1 increasing from 0.34 on 1,000 cells to 0.39 on 10,000 cells but an equal balanced accuracy of 0.29.

### Effects of the prompt context and of imperfect lists of expected cell types

Providing LLMs with the list of expected cell types raised their balanced accuracy on every dataset in our benchmark. To assess the benefit of the list compared to more general metadata, namely the tissue and species, we ran a prompt-context ablation on the 1,000-cell test sets (Methods and Supplementary Table S10). We compared GPT-OSS-120B and C2S-Scale 2B, the open-weight LLM and the scLM that scored highest with the list, and included Gemma-2-2B, the base model of C2S-Scale 2B [14, 77]. GPT-OSS-120B and Gemini 3.1 Pro also annotated clusters, which is how LLMs are commonly used [9]. For the LLMs, adding the list had a stronger effect than adding the tissue and species (Figure 3A and Supplementary Figure S4). The list raised their balanced accuracy on all four datasets, for GPT-OSS-120B per cell from 0.31 to 0.54 (Supplementary Table S11). The tissue and species raised it about half as much, mainly on HFRA and the pancreas data, and added almost nothing on top of the list. For C2S-Scale 2B, both effects on balanced accuracy were much smaller and of similar size. Gemma-2-2B stayed near zero in every condition, and most of its answers matched no Cell Ontology term (Supplementary Figure S5). Further, with the list, annotating single cells was more accurate on average than annotating clusters (Supplementary Table S11), but it needed one model call per cell instead of one per cluster and cost more than 30 times as much. For example, Gemini 3.1 Pro reached a balanced accuracy of 0.62 per cell at $8.57 per 1,000 cells, and of 0.56 per cluster at $0.25 (Supplementary Figure S6).

**Figure 3.**
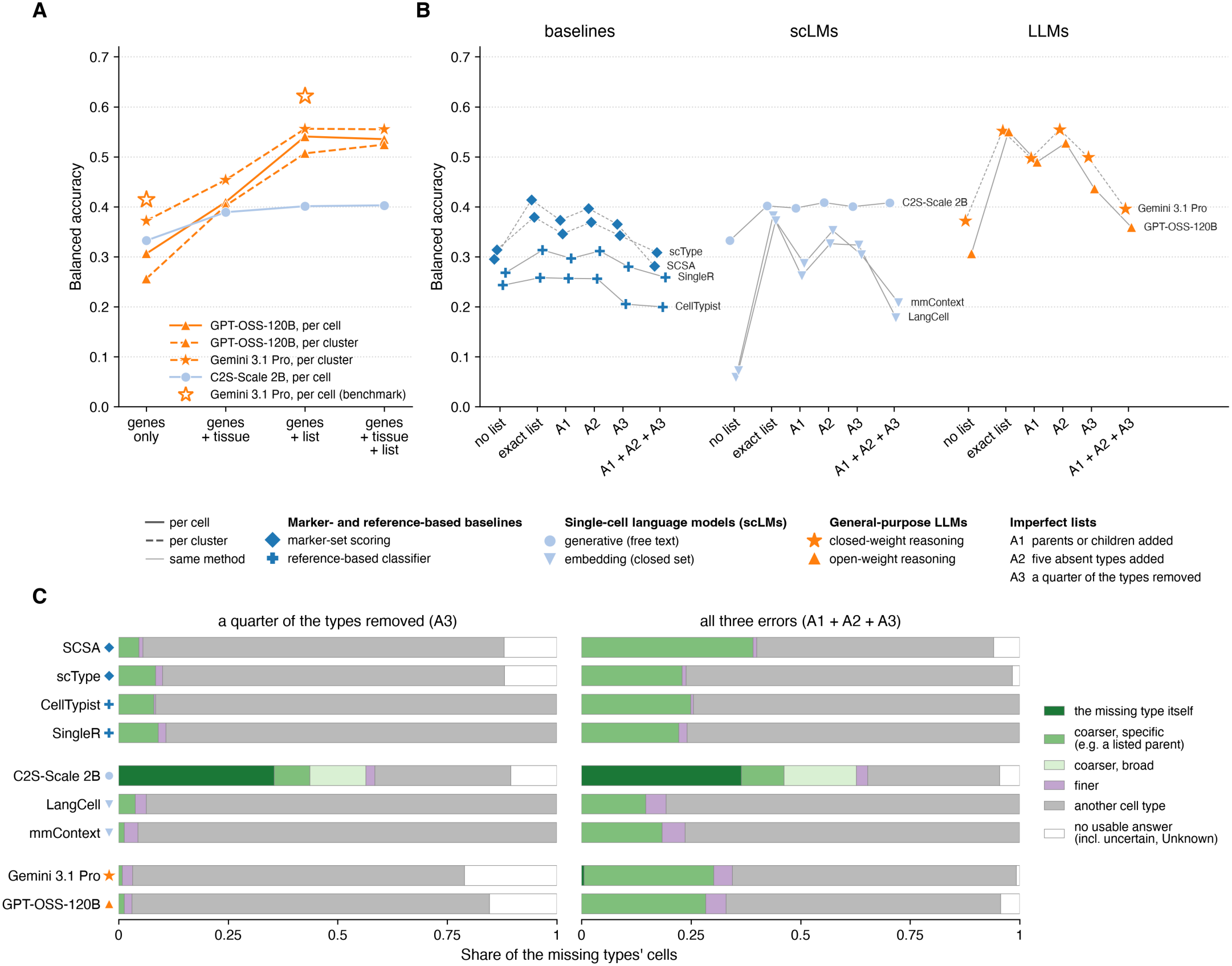
Effects of the prompt context and of imperfect lists of expected cell types. (**A**) Balanced accuracy of four arms of the prompt-context ablation, given genes only, genes with the tissue and species (genes + tissue), genes with the list of expected cell types (genes + list), or genes with both. Lines join the conditions of one arm, solid for the arms per cell and dashed for those per cluster. Open stars are the benchmark’s runs of Gemini 3.1 Pro per cell, unconstrained at genes only and constrained at genes + list. (**B**) Balanced accuracy in the imperfect-list analysis for each method without a list, with the exact list, with a parent or a child added for half of the types (A1), with five types absent from the data added (A2), with a quarter of the types removed (A3) and with all three errors (A1 + A2 + A3). Balanced accuracy is computed over all true cell types, including those removed from the list. Methods are grouped by family, and thin lines join the points of one method, dashed for the methods that annotate clusters. For comparison, the points without a list show the ablation’s genes-only runs for the generative models, GPT-OSS-120B, Gemini 3.1 Pro and C2S-Scale 2B, and the benchmark’s unconstrained runs on the same cells for the other methods. In A3 and A1 + A2 + A3, the generative models could also answer “uncertain”. (**C**) Share of answers for the cells of the types missing from the list, with A3 (left) and with all three errors (right), averaged over the missing types. An answer names the missing type itself, a coarser type, a finer type or another cell type in the Cell Ontology, or gives no usable answer. A coarser type is broad if it has more than 250 terms at or below it and specific otherwise. Colors give the method family and marker shapes the subcategory. Each marker and bar is the mean over the four datasets, at 1,000 cells each, of the mean over three seeds per dataset, except for the benchmark’s runs, which ran once per dataset.

Next, we evaluated lists that contain errors, a more realistic scenario, in an imperfect-list analysis (Methods). Combined, the three kinds of errors removed most of the benefits of the list of expected cell types (Figure 3B, Supplementary Figure S7 and Supplementary Table S12). GPT-OSS-120B dropped from 0.55 to 0.36, close to its 0.31 without a list. A similar pattern was observed for Gemini 3.1 Pro per cluster. The embedding-based scLMs lost about as much balanced accuracy, but from a lower level. The baselines lost the entire benefit of the list and ended at or below the balanced accuracy of their unconstrained runs (Figure 3B). Of all methods, only C2S-Scale 2B remained stable across the errors introduced into the list. Among the single errors, removing types (A3) lowered the balanced accuracy of every other method, whereas adding absent types (A2) had little effect on most methods. Adding parents or children (A1) reduced the balanced accuracy of the embedding-based scLMs even more than removing types, mostly because more cells were assigned a coarser type, which ontology-aware metrics such as the agreement score count as partly correct (Supplementary Figure S8 and Supplementary Table S12).

When a quarter of the types was removed from the list, every method except C2S-Scale 2B assigned most cells of the missing types to another listed type, 76 to 96% of them (Figure 3C). The generative models could then also answer “uncertain”, which changed their balanced accuracy by at most 0.01 when offered with the exact list (Supplementary Table S12). Gemini 3.1 Pro and GPT-OSS-120B gave this answer for only 21 and 15% of the missing types’ cells, and SCSA and scType answered “Unknown” for 12%. With all three errors, more of these cells went to a coarser listed type, but rarely to a finer one. C2S-Scale 2B, in contrast, did not follow the instruction to pick from the list. It never answered “uncertain” and named the missing type itself for about a third of these cells (Figure 3C and Supplementary Table S12). With the exact list, the balanced accuracy of GPT-OSS-120B was higher than that of C2S-Scale 2B on all four datasets, but with all three errors it was lower on three of them (Supplementary Figure S7).

### Rare cell type analysis

The baselines can name only the cell types that their marker database or reference covers, whereas the language models can in principle name any cell type. This difference could matter most for rare cell types, which count as much as common ones in the balanced accuracy score. We call a cell type rare when it makes up less than 1% of its full dataset, which holds for 44 of the 87 cell types, most of them in HIHA and Zeisel (Methods and Supplementary Figure S9). We compared the balanced accuracy over the rare types with that over the common types (Figure 4A). Averaged over the datasets, every family scored lower on rare than on common types, with and without the list, and so did the closed-weight LLMs. The baselines dropped most, from 0.36 on common types to 0.12 on rare types without the list and from 0.43 to 0.15 with it. The language models dropped much less, and the list raised their balanced accuracy on rare types nearly as much as on common types. For the baselines, the list added only 0.03 on rare types, against 0.07 on common types. With the list, the LLMs reached 0.47 on rare types, ahead of the scLMs and the baselines.

**Figure 4.**
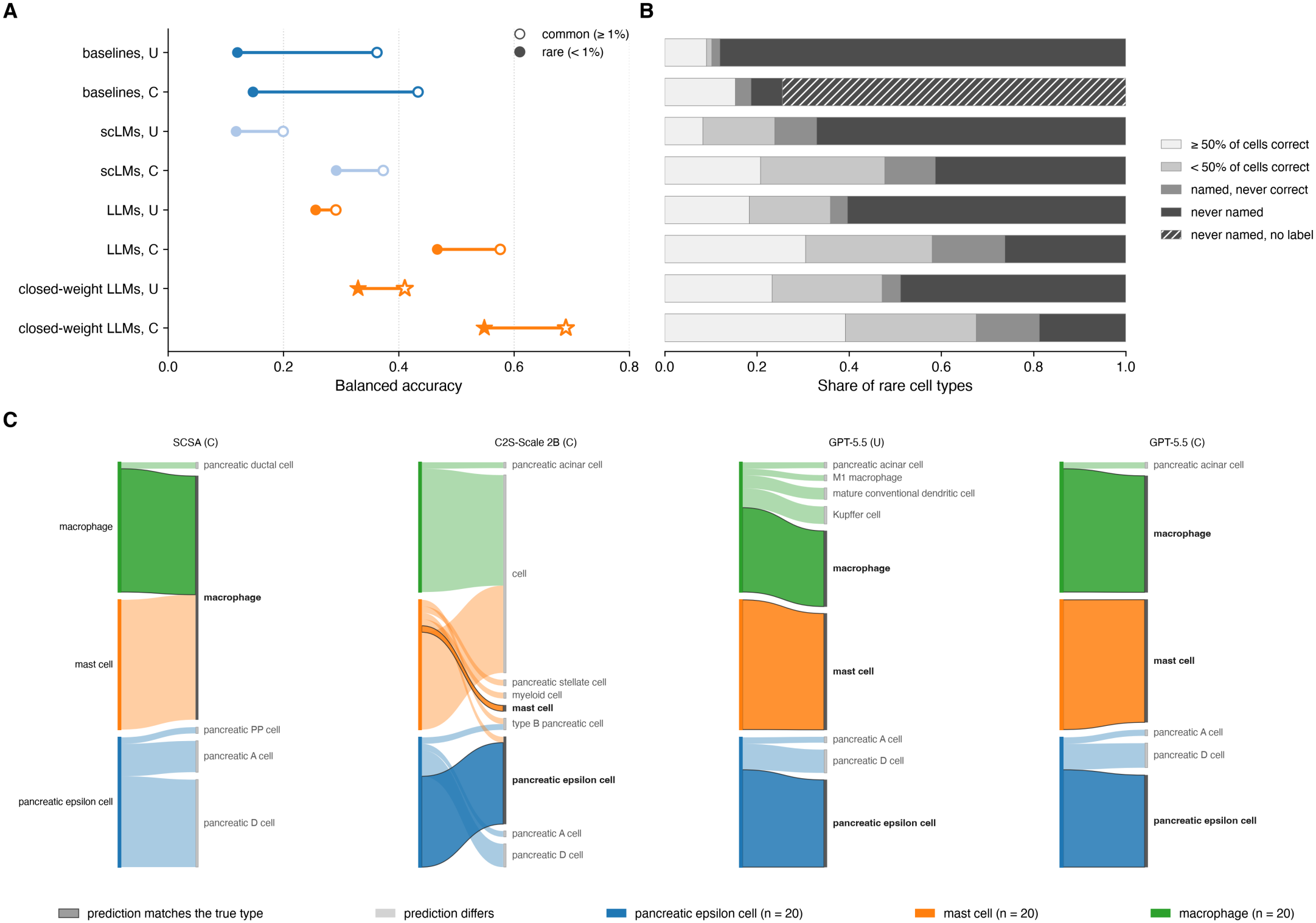
Rare cell type analysis. (**A**) Balanced accuracy, the share of correctly labeled cells averaged over cell types, computed over the common cell types (open markers) and over the rare ones (filled markers) of each dataset and averaged over the four datasets. A cell type is rare when it makes up less than 1% of its full dataset. Stars mark the closed-weight LLMs. (**B**) Outcome of each of the 44 rare cell types, as the share of types with at least half of their cells labeled correctly, with fewer than half, with none although the method used the type’s name for other cells (named, never correct), and with none and the name never used (never named). Each type counts once per method. For the constrained baselines, hatching marks the never-named types for which our mapping assigns none of the method’s labels to the type or to a more specific one (never named, no label). (**C**) Predicted labels for the cells of the three rare cell types in the human pancreas data, from left to right from SCSA and C2S-Scale 2B with the list and from GPT-5.5 without and with it. These are the constrained runs of each family with the highest balanced accuracy averaged over the four datasets. In each alluvial diagram, true types are on the left and predicted labels on the right, and ribbon width is proportional to the number of cells. A ribbon that ends at its own true type is opaque with a dark edge, and that label is set in bold. A and B include all 21 methods, and every panel uses the 1,000-cell test sets. Each row is the mean over its methods, without (U) or with (C) the list of expected cell types.

The baselines dropped on rare types mostly because they never named them, and with the list most of these types were missing from their label space (Figure 4B). Without the list, they never named 88% of the rare types, and with the list they still never named 81%. With the list, a baseline can use only those of its labels that our mapping assigns to a listed cell type or to a more specific one. On average, the baselines had no such label for 74% of the rare types, so they could not name these types at all. Where the baselines had a label for a rare type, they labeled at least half of its cells correctly in 60% of cases. The LLMs never named 60% of the rare types without the list and 26% with it, and the closed-weight LLMs 49% and 19%. With the list, the LLMs labeled at least half of the cells correctly for 31% of the rare types and the closed-weight LLMs for 39%. For another 16% and 14%, they used the name of the rare type only for cells of other types. None of the 11 LLMs labeled a single cell of five rare types correctly with the list, four of them in HIHA (two thymocyte stages, a regulatory T cell subtype and erythrocytes) and cranial visceromotor neurons in Zeisel (Supplementary Figure S9).

Cells of rare types that were labeled wrongly mostly received another cell type rather than a coarser or finer one (Supplementary Figure S10). Depending on the group, 42 to 77% of the cells of rare types were labeled as another cell type, more than for the common types in every group (30 to 55%). Figure 4C follows the cells of the three rare types in the human pancreas data. SCSA was the most accurate baseline on this dataset (Figure 2F). With the list, it labeled all 20 pancreatic epsilon cells as pancreatic D, A or PP cells, other islet endocrine types, since its marker database has no label for epsilon cells. It had a label for mast cells but labeled all 20 of them as macrophages, while it labeled 19 of the 20 macrophages correctly. With the list, C2S-Scale 2B labeled 14 of the 20 epsilon cells correctly but answered *native cell*, an obsolete Cell Ontology term replaced by the root term *cell*, for 19 of the 20 macrophages and 15 of the 20 mast cells. Without the list, GPT-5.5 already labeled all 20 mast cells and 15 of the 20 epsilon cells correctly, but with no tissue named in its prompt, it labeled 3 of the 20 macrophages as *Kupffer cells*, the macrophages of the liver. With the list, it labeled 19 of the 20 macrophages correctly. Supplementary Figures S11–S14 follow the cells of up to five of the rarest types in each dataset.

## Discussion

In this study, we benchmarked 21 methods across three different method categories on the task of cell type annotation. We created a comprehensive benchmarking pipeline implemented in nextflow which automatically runs general-purpose LLMs, domain-adapted scLMs and baseline methods on four different data sets and evaluates them against the ground truth with a broad range of evaluation metrics, both ontology-aware and ontology-independent. Our benchmark shows that the performance of language models in zero-shot cell type annotation depends strongly on the information available at inference. When provided with the complete list of expected cell types, the best general-purpose LLMs consistently outperformed scLMs and baseline annotation methods. However, their advantage decreased substantially when this information was withheld, with neither LLMs nor scLMs consistently outperforming established baselines. General-purpose LLMs appear capable of exploiting biological knowledge acquired during pretraining when the annotation problem is sufficiently constrained. Nevertheless, the strong dependence on the candidate list limits the generalizability of results obtained under ideal conditions, as the exact composition of a dataset is rarely known beforehand.

Our analysis of imperfect candidate lists further highlights the importance of label-space specification for practical applications. Removing cell types that were actually present substantially reduced annotation accuracy, whereas including additional, absent cell types had comparatively little effect. The model families differed considerably in their responses to imperfect lists. General-purpose LLMs and embedding-based scLMs benefited strongly from accurate candidates but frequently assigned cells to incorrect available labels when the true type was missing. In contrast, generative scLMs benefited less from candidate lists but were also more robust to their inaccuracies, with C2S-Scale 2B frequently predicting cell types omitted from the list. This reveals a trade-off between exploiting prior knowledge and retaining the flexibility to identify unexpected populations, suggesting that future annotation approaches could benefit from combining constrained classification with mechanisms for detecting cell types outside the candidate set. One such approach might be the usage of agentic frameworks: Extending the single-prompt, zero-shot LLM predictions into a multi-step framework, in which the LLMs are navigated by the harness.

Rare cell types remain a major challenge across all evaluated methods. Conventional marker- and reference-based approaches were particularly limited by incomplete label coverage, whereas general-purpose LLMs achieved the highest accuracy on rare populations when supplied with the expected cell types. Nevertheless, even under these favorable conditions, many rare cell types were not reliably identified, indicating that access to the correct label vocabulary does not necessarily imply an ability to distinguish closely related or transcriptionally similar populations. We observed substantial differences in annotation performance across the included datasets. Future benchmarks could benefit from including a broader range of datasets, including more tissues and different species. While we chose datasets which should not have been available for the general-purpose LLMs during training time we cannot rule out that some of the, particularly closed-weight, LLMs have previously seen some of the data. Additionally, when interpreting comparisons between model categories it should be taken into account that the concrete method of constraining the models differs for the LLMs, scLMs and baselines, based on their implementation. Finally, cell type annotations themselves represent expert-defined interpretations of biological identity and may vary in granularity or contain inaccuracies.

Domain adaptation helped, as C2S-Scale 2B was far more accurate than its base model Gemma-2-2B, which stayed near zero. Yet the general-purpose reasoning LLMs were more accurate than every scLM when given the list. For this task, general capability currently outweighs domain adaptation, and it remains open whether a frontier model adapted to single-cell data would outperform the next generation of general-purpose models.

Overall, our findings highlight both the potential and the current limitations of language models for automated single-cell annotation. Future benchmarks should incorporate larger and more diverse datasets, uncertain or incomplete candidate label spaces, and genuinely open-ended annotation scenarios. Furthermore, evaluating domain-adapted models on additional tasks, such as perturbation prediction, gene expression generation and cross-modal integration, will be essential to determine whether specialized training provides capabilities beyond those of general-purpose LLMs. Our reproducible benchmarking framework provides a basis for systematically assessing these developments.

## Key Points

- Our open Nextflow pipeline runs general-purpose LLMs, scLMs and marker- and reference-based baselines on the same cells, with and without a list of expected cell types, and scores them against the Cell Ontology.
- Constrained by the list of expected types, LLMs annotated cells most accurately. Unconstrained, no group consistently outperformed the baselines, and the small C2S-Scale 2B matched the best open-weight LLMs.
- Adding the list helped LLMs more than adding tissue and species information. Embedding-based scLMs depended on it, whereas generative scLMs barely used it.
- Introducing errors in the list of expected types harms annotation performance. Omitting present types hurt more than adding absent ones, and the cells of omitted types mostly received another listed name instead of being flagged as uncertain.
- Rare cell types remained the hardest to annotate. Baselines often had no label for them, and constrained LLMs labeled most cells correctly for only about a third of the rare types.

## Author contributions

N.B. and A.F. devised the study, implemented the software, performed analyses and wrote the manuscript. J.M., A.K., S.M.K., P.V. and L.C.G. implemented the software and performed analyses. R.K. performed analyses. L.G. implemented the software and contributed to the ontology mapping. M.H. devised the study and supervised the project. H.B. contributed to the design of the study, supervised the project and acquired funding. R.B. devised the study, supervised the project and acquired funding. All authors revised the manuscript and approved its final version.

## Data availability

The test sets and clustering samples of all four datasets are deposited on Zenodo [79], together with the answers of all runs of the prompt-context ablation and the imperfect-list analysis. HFRA [80] and HIHA [81] were downloaded from CZ CELLxGENE Discover, release 2025-11-08 of its Census, and the human pancreas data from figshare [82]. The Zeisel data come from the authors’ file l5_all_rev1.loom [83], which we downloaded from mousebrain.org in December 2025 and converted to the h5ad format. The authors now provide this file through their code repository, and the pipeline downloads our converted copy from Zenodo [79]. The SingleR references for HFRA, Zeisel and human pancreas are CELLxGENE datasets [84–86], and the reference for HIHA is MonacoImmuneData of the celldex package (GEO GSE107011).

The code of the pipeline is available at *https://github.com/BackofenLab/scLM_annotation_benchmark* . The pipeline runs with Nextflow 25.04.6 and provides a conda environment for every step, with container images as an alternative.

## Use of AI

We used Anthropic’s Claude Opus models in Claude Code and OpenAI’s ChatGPT models to help write code and optimize text formulation, and to assist in designing the model overview Figure 1. The authors directed all work, revised all text, verified every output and take full responsibility for the content.

