## Supplemental Material for "General-purpose LLMs outperform single-cell language models at cell type annotation when expected types are known"

### Contents


Supplementary Notes 3
Supplementary Note 1: Supplementary Methods 3
Preprocessing 3
Test sets 3
Clustering samples 3
Inputs of the single-cell language models 3
Inputs of the baselines 3
General-purpose LLMs 3
Ontology mapping and scoring 4
Prompt-context ablation and imperfect-list analysis 4
Supplementary Note 2: Prompt templates 4
Per-cell template, unconstrained 4
Per-cell template, constrained 4
C2S-Scale's template, unconstrained 5
C2S-Scale's template, constrained 5
InstructCell's template, unconstrained 5
InstructCell's template, constrained 5
Candidate texts of the embedding-based scLMs 5
Per-cluster template, unconstrained 5
Per-cluster template, constrained 5
Imperfect-list analysis 6
Supplementary Figures 7
Figure S1 Dependence of each method on the curated mapping 7
Figure S2 Answer granularity in the Cell Ontology, per dataset and per method subcategory 8
Figure S3 Correlation between the evaluation metrics 9
Figure S4 Prompt-context ablation per dataset 10
Figure S5 Gemma-2-2B answers in the prompt-context ablation 11
Figure S6 Balanced accuracy against cost and model calls 11
Figure S7 Imperfect-list analysis per dataset 12
Figure S8 Agreement score in the imperfect-list analysis 13
Figure S9 Recall and Lin similarity for the 44 rare cell types by method group 14
Figure S10 Answer types for the common and the rare cell types by method group 15
Figure S11 Predicted labels for the cells of the two rare cell types in HFRA 15
Figure S12 Predicted labels for the cells of the five rarest cell types in HIHA 16
Figure S13 Predicted labels for the cells of the five rarest cell types in Zeisel 17
Figure S14 Predicted labels for the cells of the three rare cell types in human pancreas 18
Supplementary Tables 19
Table S1 Datasets of the benchmark 19
Table S2 Annotation methods 20
Table S3 Inputs of the baselines 21
Table S4 Evaluation metrics 21
Table S5 Benchmark results for HFRA 22
Table S6 Benchmark results for HIHA 24
Table S7 Benchmark results for Zeisel 26
Table S8 Benchmark results for human pancreas 28
Table S9 Answers outside the list of expected cell types 30
Table S10 Arms of the prompt-context ablation 30
Table S11 Results of the prompt-context ablation 31
Table S12 Results of the imperfect-list analysis 32

### Supplementary Notes

#### Supplementary Note 1: Supplementary Methods

##### Preprocessing

Quality control runs on the whole dataset, not per batch. The outlier rule for the total counts and the number of detected genes takes their log-transformed values when the downloaded file already holds them, which was the case only for Zeisel, and the untransformed values otherwise. The rule on mitochondrial counts finds the mitochondrial genes by the prefix MT- of their names and therefore acted only on Zeisel, since HFRA and HIHA are indexed by Ensembl identifiers at this step and the pancreas data hold no mitochondrial genes. In the pancreas data, quality control removed all cells of the two Fluidigm C1 studies, labeled SMARTer and Fluidigm C1 in the compilation, and 2,239 of the 2,394 Smart-seq2 cells, whose read counts lie far above the molecule counts of the other technologies. HFRA, HIHA and Zeisel carry Ensembl identifiers, and the gene symbols of the pancreas data were converted to Ensembl identifiers with BioMart after normalization, which dropped the genes without a unique identifier.

##### Test sets

Cell types with fewer than 20 cells after quality control are left out, which concerns only the Schwann cells (17) and T cells (7) of the pancreas data. The other types receive cells in proportion to their share of the dataset, at least 20, with the excess taken proportionally from the larger types, and the cells are drawn at random within each type (seed 42). The labels of the test sets, and thus the lists of expected cell types, are names of Cell Ontology terms except for six original labels, Enteric glia, Microglia and Satellite glia in Zeisel and acinar, delta and epsilon in the pancreas data.

##### Clustering samples

scType, SCSA and the per-cluster arms of the ablation cluster a larger sample of each dataset, which also provides the highly variable genes of the ablation. The sample starts from the union of both test sets, adds cells of every type until the type has 150 cells or all of its cells, and is then filled in proportion to the full dataset to about 100,000 cells (seed 42). For the pancreas data, it holds all 11,704 cells, including the types left out of the test sets.

##### Inputs of the single-cell language models

C2S-Scale 2B and 27B (`vandijklab/C2S-Scale-Gemma-2-2B` and `-27B`) read the cell sentence and the organism and answer greedily with at most 20 new tokens. InstructCell (`zjunlp/InstructCell-instruct`) reads the raw counts of the 18,961 genes of its vocabulary, absent genes set to zero, and scales and log-transforms them itself. As on its model card, it samples its answer with top-p 0.95, top-k 50 and at most 256 new tokens (seed 1234). LangCell and CellWhisperer read the raw counts as a ranking of at most 2,048 genes, as Geneformer does, and LangCell scores a candidate as 0.1 times the softmax of its embedding similarity, at temperature 0.05, plus 0.9 times the softmax of its matching score between cell and text. mmContext (`jo-mengr/mmcontext-pubmedbert-gs10k`) reads the log-normalized expression of its 10,000 preselected genes, absent genes as zero. On the mouse Zeisel data, the 5,069 genes without a one-to-one human ortholog keep their mouse identifiers, and the expression values stay unchanged. Without the list of expected cell types, the embedding-based scLMs choose among 3,169 names of current terms of the Cell Ontology release of 16 October 2025, and with it among the test set's own labels. C2S-Scale and the Gemma-2-2B of the ablation are subject to the Gemma Terms of Use, and the pipeline downloads their weights from Hugging Face instead of redistributing them.

##### Inputs of the baselines

The baselines used the models, references and tissues of Supplementary Table S3. CellTypist annotates each cell from its log-normalized counts without majority voting. SingleR (version 2.8.0, celldex 1.16.0) uses its default settings, the Cell Ontology labels of MonacoImmuneData and the CELLxGENE references reduced to at most 500 cells per label (seed 0). scType clusters the clustering sample with 2,000 variable genes, 30 principal components, 20 neighbors and the Louvain method at resolution 0.8, and labels a cluster Unknown when its best score is below a quarter of its number of cells. SCSA clusters it with 2,000 variable genes, 50 principal components, 15 neighbors and the Louvain method at resolution 1.0, scores CellMarker genes with a fold change of at least 1.5 and a p-value below 0.01, and labels a cluster Unknown when no cell type scores for it. In their constrained runs, the baselines keep the labels that our mapping resolves to a listed type or to a type below it, and a label below several listed types answers with the nearest one, the fewest is_a steps above, with ties broken by the term identifier. A coarser label never counts, and a label that the mapping does not cover is dropped. CellTypist then takes the best kept label, SingleR trains on the kept reference labels, scType keeps the marker sets of the kept labels and SCSA the scores of the kept labels.

##### General-purpose LLMs

OpenRouter sorted the providers of each model by price and allowed fallbacks. It recorded the model versions `anthropic/claude-4.7-opus-20260416`, `anthropic/claude-4.6-sonnet-20260217`, `google/gemini-3.1-pro-preview-20260219`, `openai/gpt-5.5-20260423`, `deepseek/deepseek-v3.2-20251201`, `openai/gpt-oss-120b`, `openai/gpt-oss-20b`, `moonshotai/kimi-k2.6-20260420`, `meta-llama/llama-3.1-70b-instruct`, `meta-llama/llama-3.3-70b-instruct` and `meta-llama/llama-4-maverick-17b-128e-instruct`. The constrained runs were sent between 6 and 9 May 2026 and the unconstrained runs between 10 and 26 May 2026. Requests timed out after 90 seconds. A request that timed out or met a rate limit or a server error was sent again after a growing pause, up to five times in all, and a call that still failed was repeated up to two more times, so a cell received at most 15 requests. Kimi-K2.6 returned no answer for 83 to 398 of about 1,000 cells per dataset unconstrained and for 5 to 211 constrained, mostly after long reasoning that reached a provider's output limit or stopped without a final answer. Claude Opus 4.7 used almost no reasoning tokens despite the requested budget, about 5,400 to 5,700 in total over about 4,000 calls per setting.

#### Ontology mapping and scoring

The direct match ignores case and runs of spaces and matches neither synonyms nor retired identifiers. A name ending in (Mmus), (Homo sapiens), ", human" or ", mouse" counts as its direct parent if that parent has the same name without the tag, which concerns 51 terms of the release of 8 June 2026. True labels go through the same lookup, after only surrounding whitespace is removed and the case is lowered. The text similarity uses the [CLS] embedding of SapBERT (`cambridgeltl/SapBERT-from-PubMedBERT-fulltext`) for at most 32 tokens, and its baseline, the mean cosine over all pairs of distinct term names, is 0.434.

#### Prompt-context ablation and imperfect-list analysis

Supplementary Table S10 lists the arms of the ablation. The seeds 1234, 5678 and 9012 set the request seed, the order of the listed types and the shuffled gene order. The arms queried through OpenRouter used its default routing. Every arm took the first non-empty line of a response as its answer, skipping header lines such as `Cell 1:`. The 5,000 variable genes were selected on the clustering sample with the Seurat method, accounting for batches, after removing the genes whose symbols start with RPS, RPL, MT- or HB. The per-cluster arms clustered the same sample with the Leiden method (random state 0) and took the 25 genes that best distinguish each cluster by the Wilcoxon test after the same removal. In the imperfect-list analysis, each condition took the first types of the order drawn for its dataset and seed. A1 adds a parent or a child for half of the true types, A2 adds five absent types, A3 removes a quarter of the types, at least one, and A1 + A2 + A3 does all three. Half and a quarter of the types are rounded to whole types. A parent is a direct parent with at most 250 terms at or below it, added types come from the CELLxGENE subset of the Cell Ontology without species-qualified names, and an absent type is a child of such a parent that is unrelated to every true type. Both experiments mapped new answers into a table of their own by a fixed rule. It spells out Greek letters, removes diacritics and punctuation, reads + as positive and drops Gemma's lead-in phrase. It then accepts, in this order, one of 15 entries checked by hand, 12 equivalents and 3 generalizations to a parent, the name or an exact or related synonym of a term, the same after removing a leading identifier, article or number or with cell or neuron appended, and a unique prefix of at least six characters that covers at least half of the matching name. Every other answer maps to cell.

#### Supplementary Note 2: Prompt templates

The prompt templates of the benchmark, the prompt-context ablation and the imperfect-list analysis are reproduced below, with the substitution slots in braces. **`num_genes`** caps the gene list at 200 per cell and 25 per cluster. **`cell_sentence`** and **`marker_genes`** carry the gene names, and **`organism`** names the dataset's organism, Homo sapiens or Mus musculus. **`unique_cell_types`** is the list of expected cell types. The per-cell templates receive it as a Python list, permuted independently for each cell, the per-cluster templates separated by commas and permuted for each cluster, and the templates of C2S-Scale and InstructCell separated by commas and sorted. **`context`** is empty in the benchmark. In the conditions with the tissue and species, the prompt-context ablation fills it with them, **`human retina`** for HFRA, **`human blood`** for HIHA, **`mouse nervous system`** for Zeisel and **`human pancreas`** for human pancreas, and with the tissue alone in C2S-Scale's template, which already names the organism (Supplementary Table S10). The benchmark sent each prompt to the general-purpose LLMs with an empty system message, and the ablation's arms through an API sent it as the only message.

##### Per-cell template, unconstrained

Used in the benchmark by the general-purpose LLMs and in the ablation by GPT-OSS-120B and Gemma-2-2B per cell, in the conditions without the list.

```
You are an expert at classifying cell types based on gene expression.
Given below is a list of up to {num_genes} gene names ordered by descending expression level in a single cell.
Your task is to identify the exact cell type this gene profile{context} represents.
Cell sentence: {cell_sentence}
Important rules:
1. Respond with only the cell type label of the predicted cell type.
2. Use the canonical cell type label from the OBO Foundry Cell Ontology (CL) — the primary label string as it appears in CL, not a synonym,
abbreviation, or free-form description.
3. Do not include explanations, reasoning, or extra words in the output.
4. Do not repeat the question or input in the output.
Output:
```

##### Per-cell template, constrained

Used in the benchmark by the general-purpose LLMs and in the ablation by GPT-OSS-120B and Gemma-2-2B per cell, in the conditions with the list.

```
You are an expert at classifying cell types based on gene expression.
Given below is a list of up to {num_genes} gene names ordered by descending expression level in a single cell.
Your task is to identify the exact cell type this gene profile{context} represents from this list of cell types: {unique_cell_types}.
Cell sentence: {cell_sentence}
Important rules:
1. Respond with only the cell type label of the predicted cell type.
2. Do only choose a cell type from the list of cell types provided.
```

```
3. Do not include explanations, reasoning, or extra words in the output.
4. Do not repeat the question or input in the output.
Output:
```

##### C2S-Scale's template, unconstrained

Used in the benchmark by C2S-Scale 2B and 27B and in the ablation by C2S-Scale 2B, in the conditions without the list.

```
The following is a list of {num_genes} gene names ordered by descending expression level in a {organism}{context} cell. Your task is to give the cell
type which this cell belongs to based on its gene expression.
Cell sentence: {cell_sentence}.
The cell type corresponding to these genes is:
```

##### C2S-Scale's template, constrained

Used in the benchmark by C2S-Scale 2B and 27B and in the ablation by C2S-Scale 2B, in the conditions with the list.

```
The following is a list of {num_genes} gene names ordered by descending expression level in a {organism}{context} cell. Your task is to give the cell
type which this cell belongs to based on its gene expression. Pick from this list: {unique_cell_types}.
Cell sentence: {cell_sentence}.
The cell type corresponding to these genes is:
```

##### InstructCell's template, unconstrained

Used in the benchmark. `species` is human or mouse, `tissue` is `retina (fetal)` for HFRA, `blood (peripheral blood immune cells)` for HIHA, `nervous system (whole mouse CNS and PNS: cortex, hippocampus, cerebellum, midbrain, brainstem, spinal cord, and peripheral sensory/sympathetic/enteric ganglia)` for Zeisel and `pancreas` for human pancreas, and InstructCell replaces `input` with embeddings of the cell that its expression encoder computes from the raw counts. The template is the example prompt of InstructCell's model card without the sequencing method.

```
Can you help me annotate this single cell from a {species}? It is derived from {tissue}. The gene expression profile is {input}. Thanks!
```

##### InstructCell's template, constrained

Used in the benchmark. It adds the expected cell types in the benchmark's own wording, not in one of the option phrasings InstructCell was trained with.

```
Can you help me annotate this single cell from a {species}? It is derived from {tissue}. Pick from this list: {unique_cell_types}. The gene expression
profile is {input}. Thanks!
```

##### Candidate texts of the embedding-based scLMs

Used in the benchmark and the imperfect-list analysis. These models compare each cell with a text for every candidate cell type, all 3,169 Cell Ontology names when unconstrained, the expected cell types when constrained and the drawn list in the imperfect-list analysis. CellWhisperer and mmContext take the candidate names as written. LangCell takes the text below, with the name and definition from the Cell Ontology file its authors provide, and `cell type: {candidate}` for a candidate that file lacks.

```
cell type: {name}. {definition};
```

##### Per-cluster template, unconstrained

Used in the ablation by GPT-OSS-120B and Gemini 3.1 Pro per cluster, in the conditions without the list.

```
You are an expert at annotating cell clusters from single-cell RNA sequencing data.
Given below are the top {num_genes} marker genes of one cell cluster, ordered by how strongly they distinguish it from all other clusters.
Your task is to identify the cell type of this{context} cluster.
Marker genes: {marker_genes}
Important rules:
1. Respond with only the cell type label of the predicted cell type.
2. Use the canonical cell type label from the OBO Foundry Cell Ontology (CL) — the primary label string as it appears in CL, not a synonym,
abbreviation, or free-form description.
3. Do not include explanations, reasoning, or extra words in the output.
4. Do not repeat the question or input in the output.
Output:
```

##### Per-cluster template, constrained

Used in the ablation by GPT-OSS-120B and Gemini 3.1 Pro per cluster, in the conditions with the list.

```
You are an expert at annotating cell clusters from single-cell RNA sequencing data.
Given below are the top {num_genes} marker genes of one cell cluster, ordered by how strongly they distinguish it from all other clusters.
Your task is to identify the cell type of this{context} cluster from this list of cell types: {unique_cell_types}.
Marker genes: {marker_genes}
Important rules:
1. Respond with only the cell type label of the predicted cell type.
2. Do only choose a cell type from the list of cell types provided.
```

```
3. Do not include explanations, reasoning, or extra words in the output.
4. Do not repeat the question or input in the output.
Output:
```

##### Imperfect-list analysis

The analysis used the constrained templates, the per-cell template for GPT-OSS-120B, C2S-Scale's own for C2S-Scale 2B and the per-cluster template for Gemini 3.1 Pro, with the drawn list in place of the dataset's own. With the exact list + uncertain, A3 and A1 + A2 + A3, the templates add the rule below, as rule 5 of the per-cell and per-cluster templates and directly after the list in C2S-Scale's template.

```
If none of the listed cell types fits, answer uncertain.
```

#### Supplementary Figures

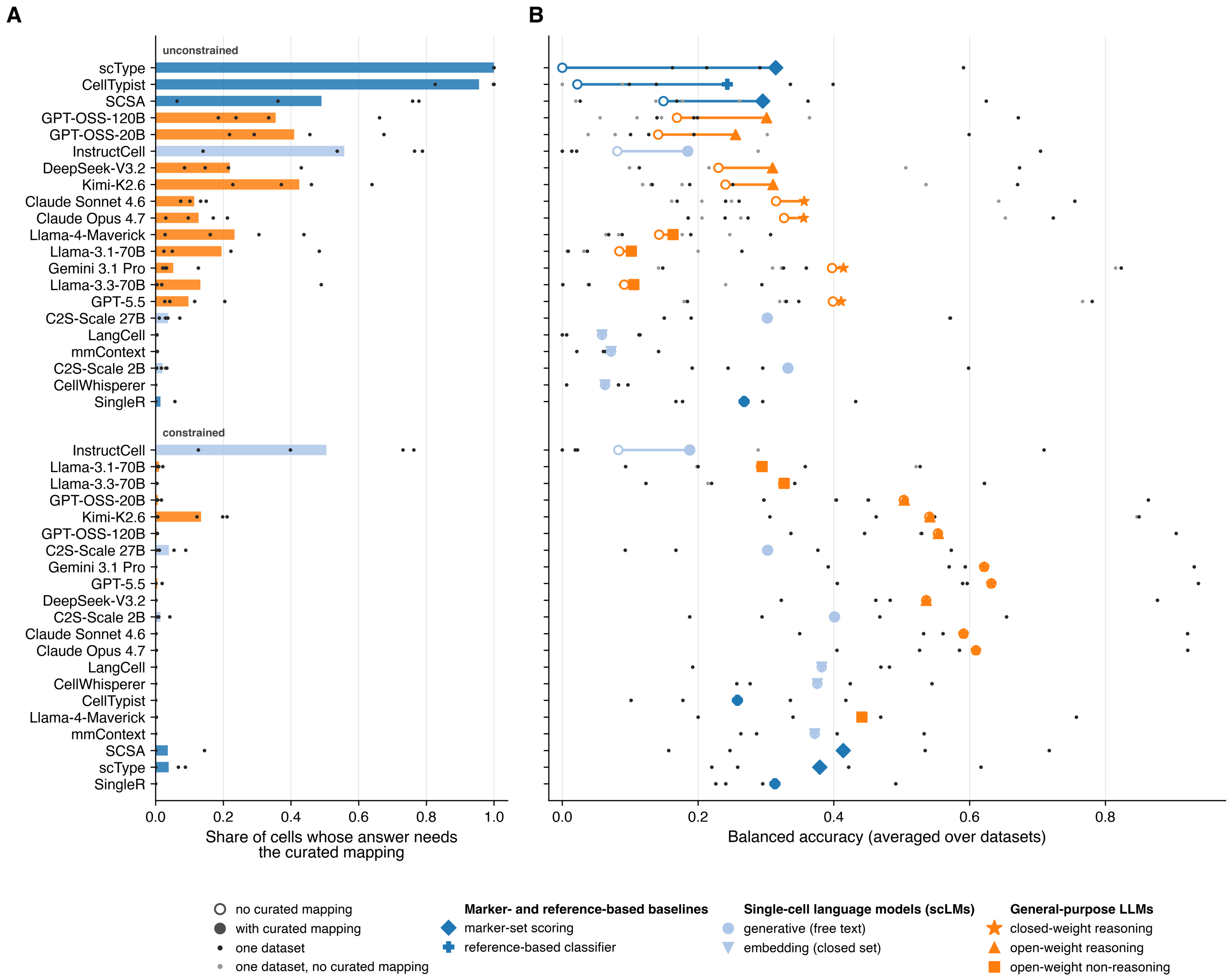


**Figure S1 Dependence of each method on the curated mapping.** Each panel shows the 21 methods twice, in an unconstrained and a constrained block. Within each block, A and B share one row order, sorted by the gap between the two symbols in B, largest first. (**A**) Share of cells whose answer needs the curated mapping, because it is neither a Cell Ontology term nor one of the dataset's own labels as written. Bars are means over the four datasets, dots give the per-dataset values, and bar color gives the method family. (**B**) Balanced accuracy when only the answers that need no curated mapping count as correct (open circles), and with the curated mapping of the other answers (filled symbols). These are the per-method values behind Figure 2G. Symbols give the subcategory, as in Figure 1B, and where only one is visible, the two values overlap. Large symbols are means over the four datasets, and dots give the per-dataset values, dark with the curated mapping and light without it.

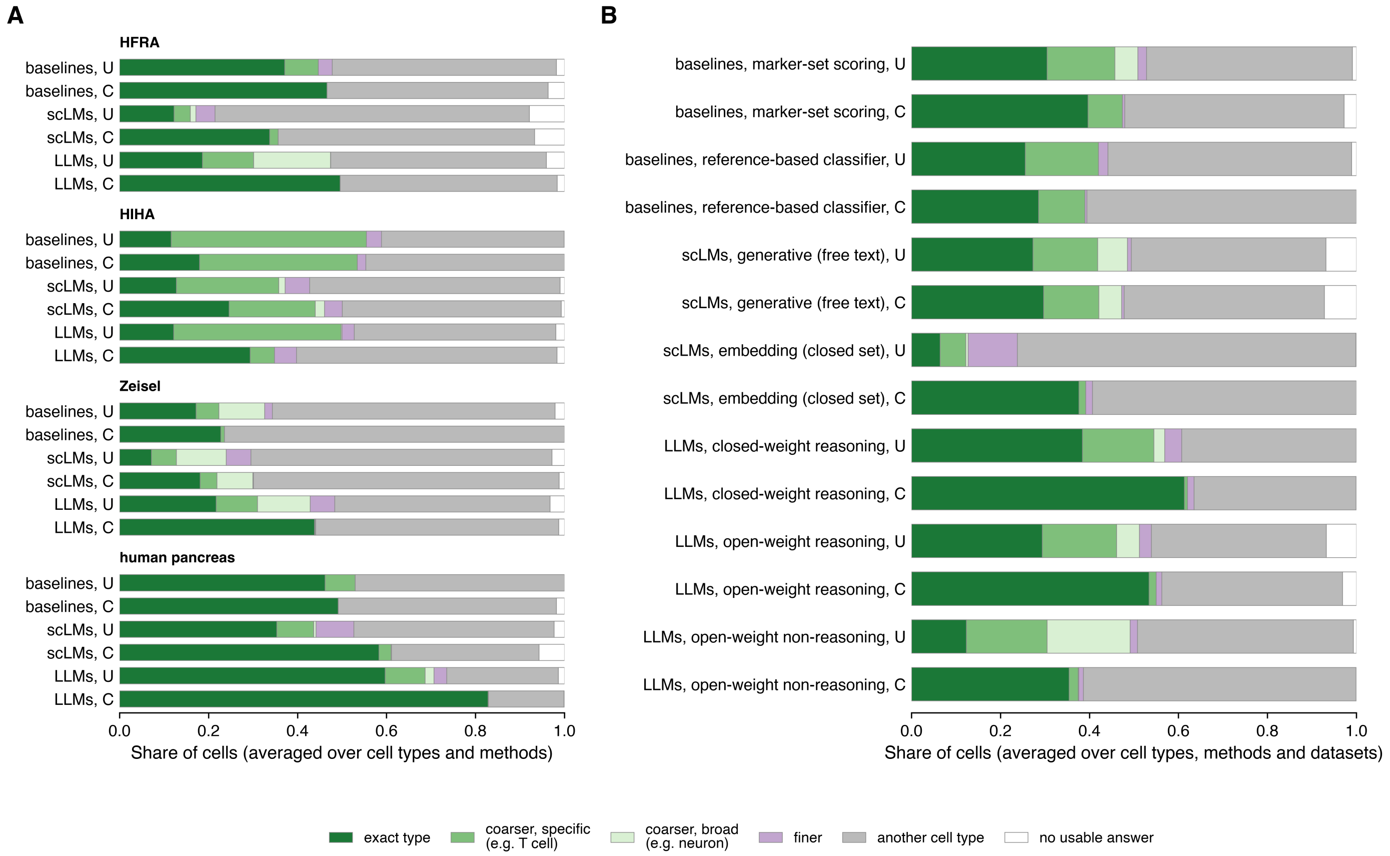


**Figure S2 Answer granularity in the Cell Ontology, per dataset and per method subcategory.** Each bar splits the cells of one group by where its answers fall relative to the true cell type. An answer is the exact type, a coarser or a finer type, another cell type, or no usable answer. Coarser types are split into specific ones and broad ones, which cover more than 250 cell types in the ontology. Shares are averaged over cell types, as in balanced accuracy, and over the methods of the group. (**A**) Each dataset, by method family and setting. (**B**) Each method subcategory, as in Figure 1B, in both settings, averaged over the four datasets. Figure 2H shows the family means over the four datasets. U marks unconstrained runs, without the list of expected cell types, and C constrained runs, with it. HFRA is the Human Fetal Retina Atlas and HIHA the Human Immune Health Atlas. LLMs are general-purpose large language models and scLMs single-cell language models.

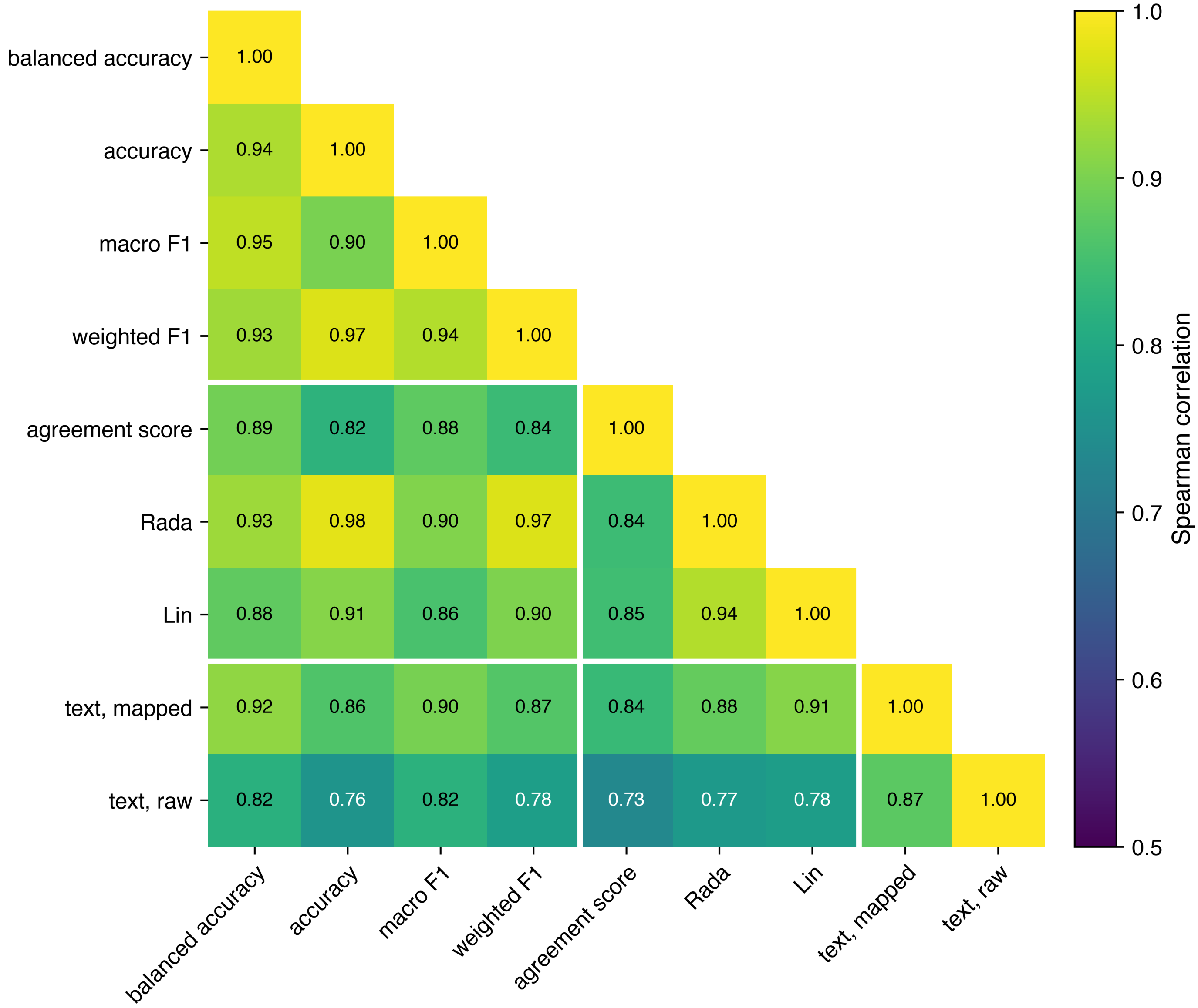


**Figure S3 Correlation between the evaluation metrics.** Spearman correlation between each pair of metrics over the 42 runs, the 21 methods in both settings. Correlations are computed within each dataset and averaged over the four datasets. The metrics are grouped into exact-match metrics, ontology-aware metrics and text similarity. Text similarity compares the meaning of the answer with the Cell Ontology name of the true cell type, after the mapping to the Cell Ontology (mapped) or as the model wrote it (raw), which needs no mapping of the answers.

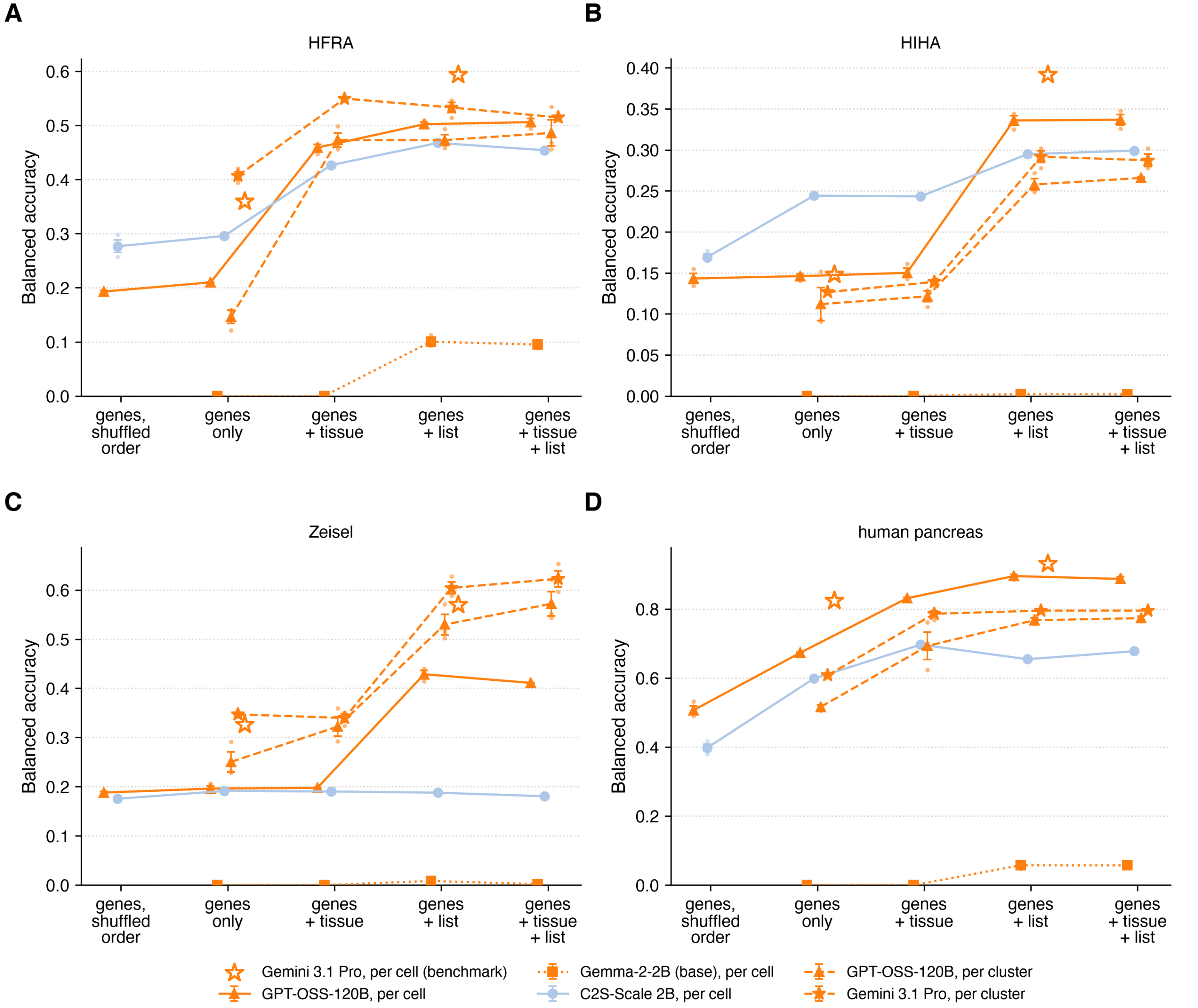


**Figure S4 Prompt-context ablation per dataset.** Balanced accuracy in the prompt-context ablation on HFRA (A), HIHA (B), Zeisel (C) and human pancreas (D), given the genes in shuffled order (genes, shuffled order), genes only, genes with the tissue and species (genes + tissue), genes with the list of expected cell types (genes + list), or genes with both. Only GPT-OSS-120B and C2S-Scale 2B per cell received the genes in shuffled order, the genes of genes only in random order. Lines join the conditions of one arm. Large markers are means over three seeds, with the standard error of the mean as error bars, and small points are the single seeds. Arms are offset horizontally within each condition. Open stars are the benchmark's single runs of Gemini 3.1 Pro per cell, unconstrained at genes only and constrained at genes + list.

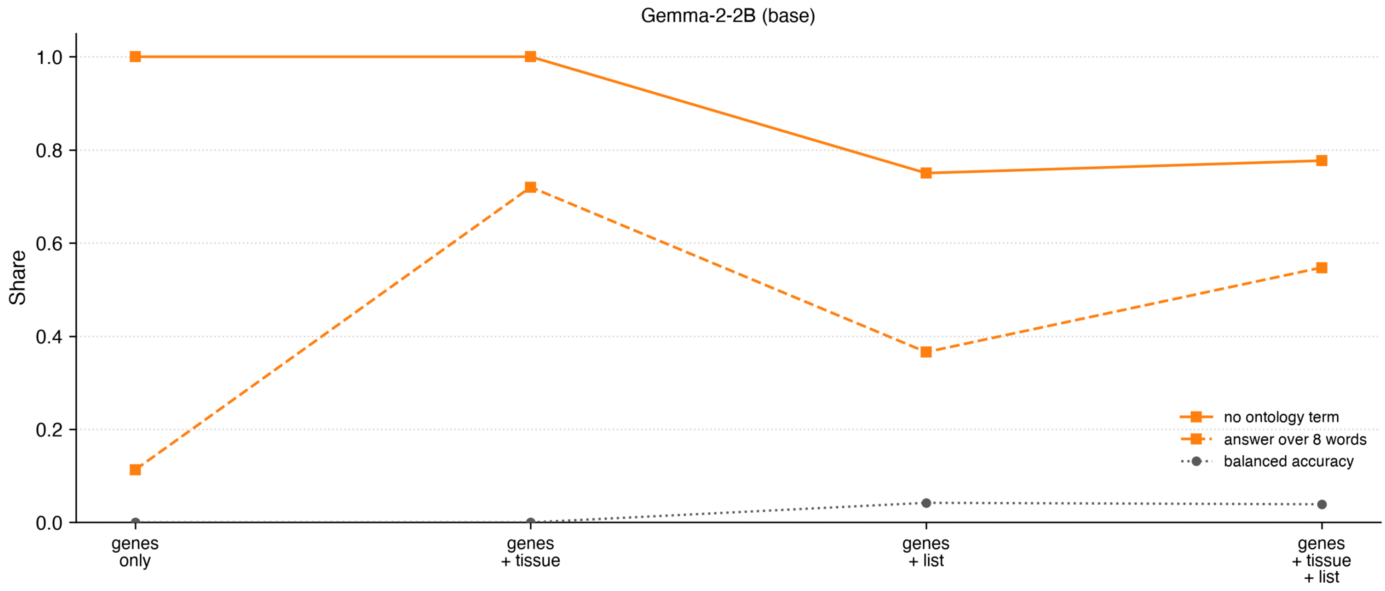


**Figure S5 Gemma-2-2B answers in the prompt-context ablation.** Answers of Gemma-2-2B per cell, the base model of C2S-Scale 2B, given genes only, genes with the tissue and species (genes + tissue), genes with the list of expected cell types (genes + list), or genes with both. Lines show the share of answers that are not the name or identifier of a Cell Ontology term (no ontology term), the share that runs past eight words (answer over 8 words) and the balanced accuracy. Values are means over the four datasets, at 1,000 cells each, and three seeds.

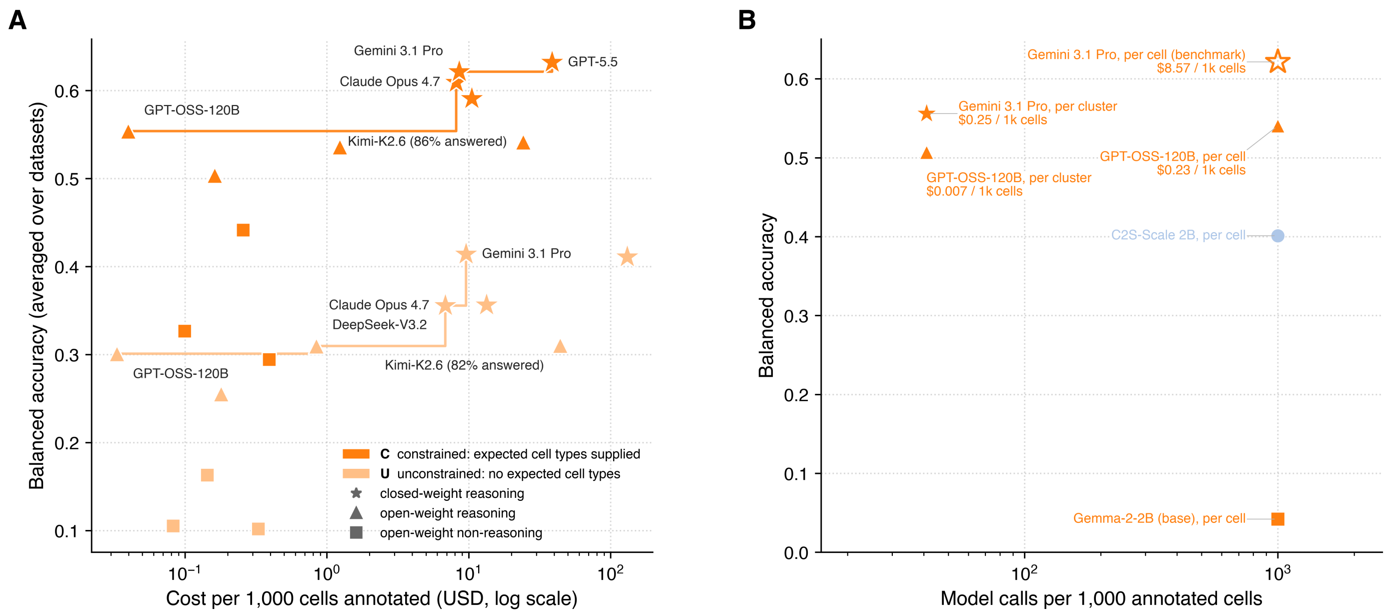


**Figure S6 Balanced accuracy against cost and model calls.** (**A**) Balanced accuracy of each LLM of the benchmark against its cost per 1,000 annotated cells, on a logarithmic scale. Colors give the setting and marker shapes the subcategory. The step line of each setting joins the cost-efficient models, those that no cheaper model matches or exceeds. Models on these lines are named, as are models that answered fewer than 98% of their calls, with the share of answered calls beside the name. (**B**) Balanced accuracy of the arms of the prompt-context ablation given genes with the list of expected cell types (genes + list), against the number of model calls per 1,000 annotated cells, one per cell or one per cluster that the cells fall into, on a logarithmic scale. Labels give the cost per 1,000 cells, and the two arms without a cost ran on local hardware. The open star is the benchmark's constrained run of Gemini 3.1 Pro per cell, with its cost from A. Balanced accuracy is averaged over the four datasets, at 1,000 cells each, and for the arms of the ablation also over three seeds.

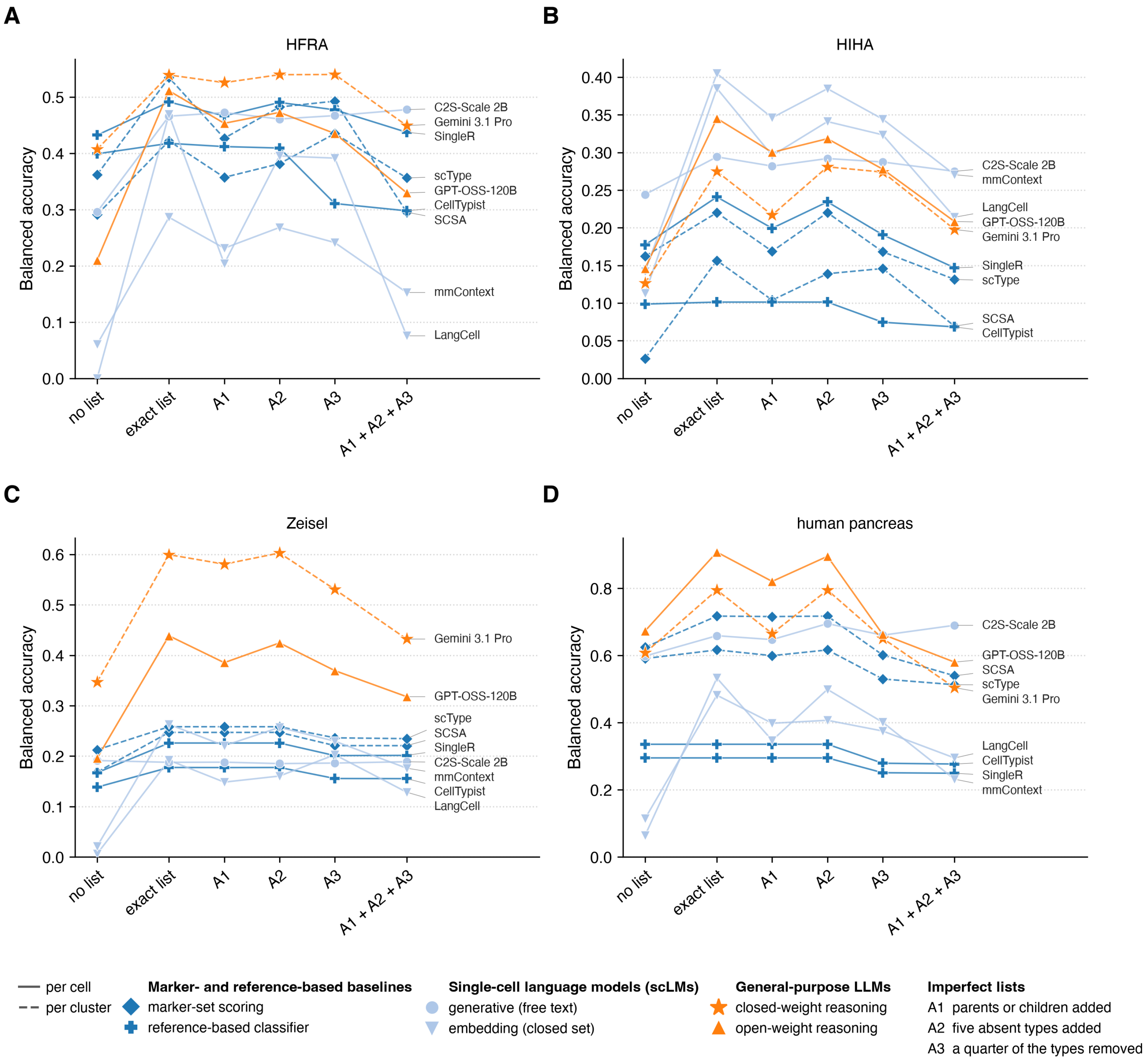


**Figure S7 Imperfect-list analysis per dataset.** Balanced accuracy in the imperfect-list analysis on HFRA (A), HIHA (B), Zeisel (C) and human pancreas (D) for each method without a list, with the exact list of expected cell types, with a parent or a child added for half of the types (A1), with five types absent from the data added (A2), with a quarter of the types removed (A3) and with all three errors (A1 + A2 + A3). Balanced accuracy is computed over all true cell types, including those removed from the list. Each line is one method, named at its last point. Colors give the method family and marker shapes the subcategory. For comparison, the points without a list show the prompt-context ablation's genes-only runs for the generative models, GPT-OSS-120B, Gemini 3.1 Pro and C2S-Scale 2B, and the benchmark's unconstrained runs on the same cells for the other methods. In A3 and A1 + A2 + A3, the generative models could also answer "uncertain". Each marker is the mean of one method over its three seeds on that dataset, except for the benchmark's single runs.

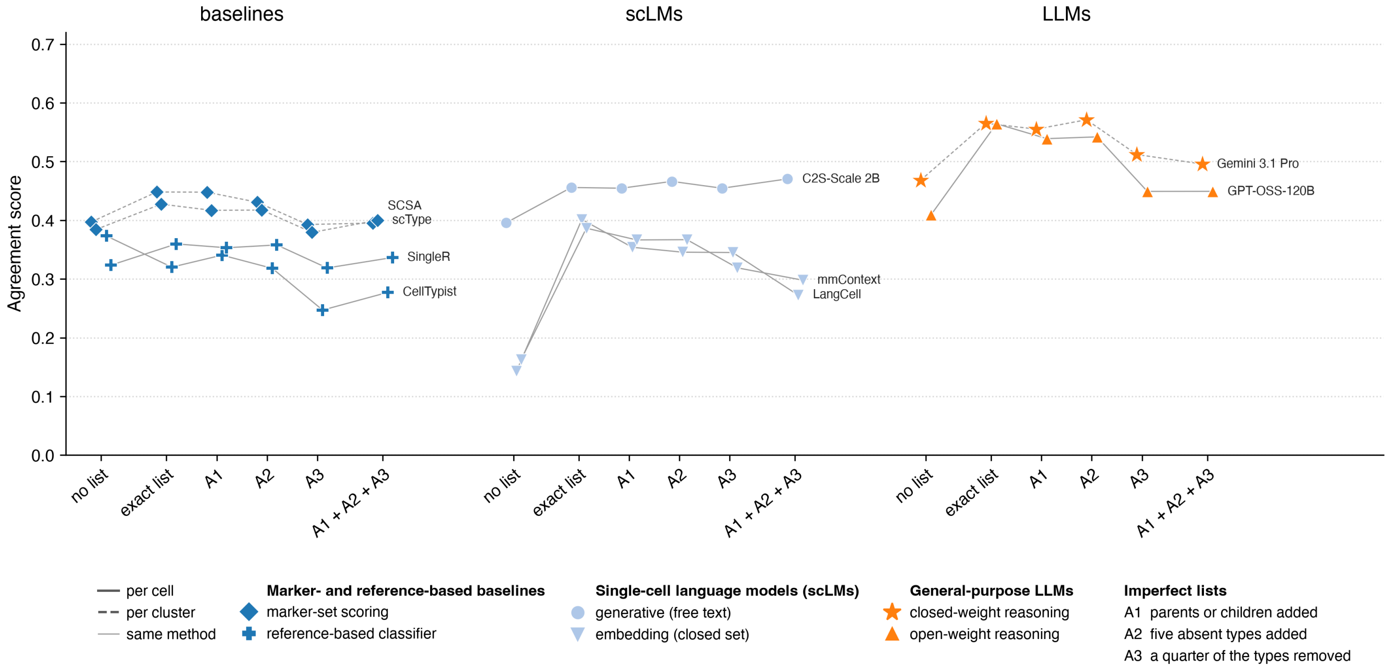


**Figure S8 Agreement score in the imperfect-list analysis.** Agreement score of each method without a list, with the exact list of expected cell types, with a parent or a child added for half of the types (A1), with five types absent from the data added (A2), with a quarter of the types removed (A3) and with all three errors (A1 + A2 + A3). The agreement score gives a cell 1 for its true type, 0.5 for a finer type or for a coarser type with at most 250 terms at or below it in the Cell Ontology, and 0 otherwise, averaged over the cells of each type and then over all true cell types, including those removed from the list. Methods are grouped by family, and thin lines join the points of one method, dashed for the methods that annotate clusters. For comparison, the points without a list show the prompt-context ablation's genes-only runs for the generative models, GPT-OSS-120B, Gemini 3.1 Pro and C2S-Scale 2B, and the benchmark's unconstrained runs on the same cells for the other methods. In A3 and A1 + A2 + A3, the generative models could also answer "uncertain". Colors give the method family and marker shapes the subcategory. Each marker is the mean over the four datasets, at 1,000 cells each, of the mean over three seeds per dataset, except for the benchmark's runs, which ran once per dataset.

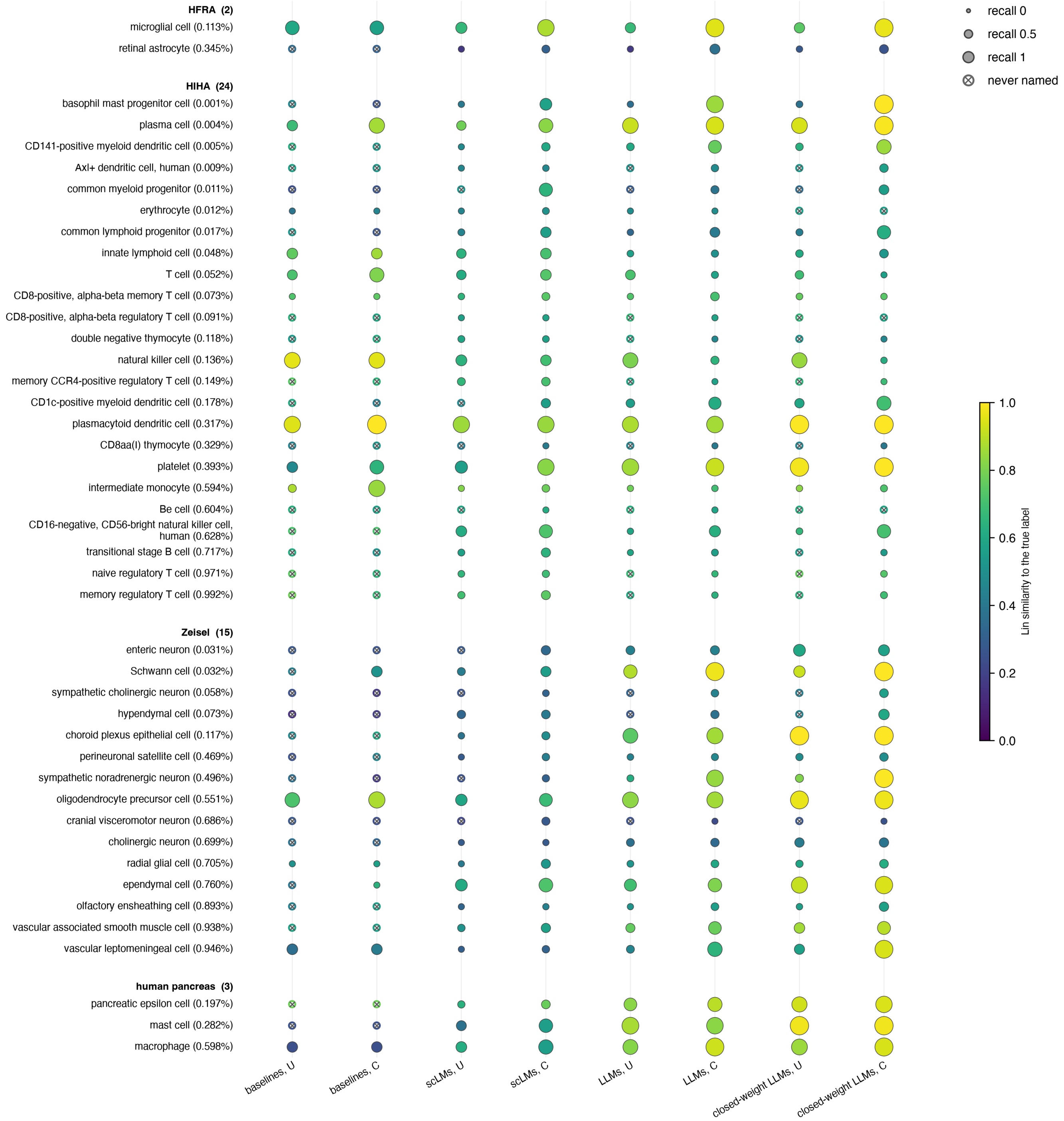


**Figure S9 Recall and Lin similarity for the 44 rare cell types by method group.** The 44 cell types that make up less than 1% of their full dataset, grouped by dataset with the number of such types beside each dataset name and ordered by their share of the full dataset. Columns are the baselines, the scLMs, the LLMs and the four closed-weight LLMs, which also count among the LLMs, each without (U) and with (C) the list of expected cell types. Marker area gives the recall of a cell type, the share of its cells labeled correctly, and marker color the Lin similarity between the answer and the true type in the Cell Ontology, averaged over the type's cells. Both are means over the methods of a column. A ring with a cross marks a cell type that no method of the column ever named. All methods ran on the 1,000-cell test sets.

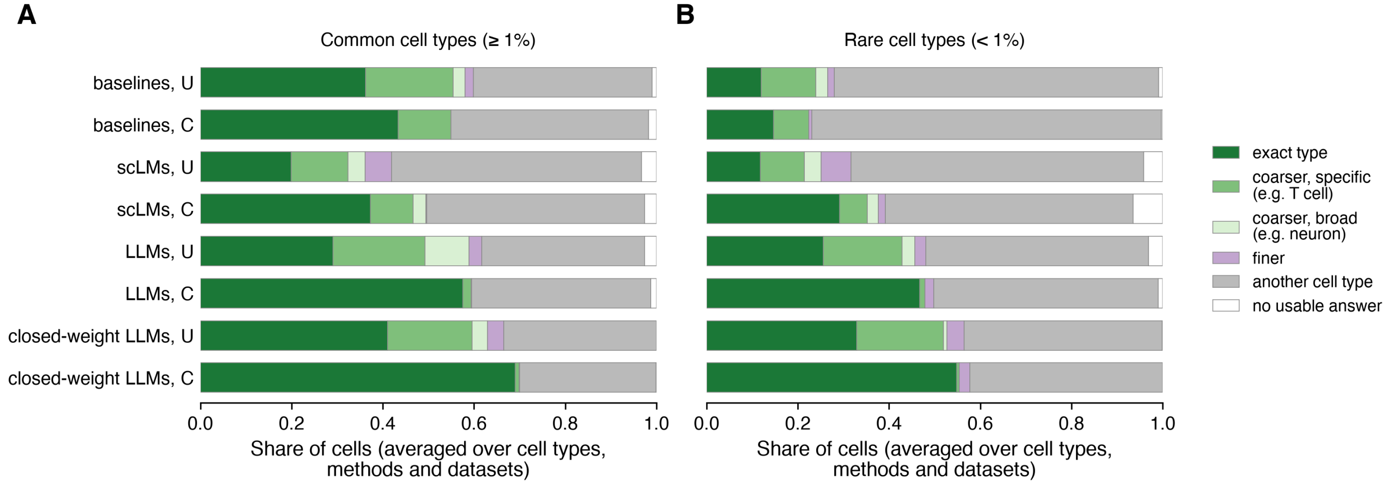


**Figure S10 Answer types for the common and the rare cell types by method group.** Share of cells whose answer names the true type, a coarser type, a finer type or another cell type in the Cell Ontology, or gives no usable answer, for the common cell types (**A**) and the rare ones (**B**). A coarser type is broad if it has more than 250 terms at or below it and specific otherwise. A cell type is rare when it makes up less than 1% of its full dataset. Rows are the baselines, the scLMs, the LLMs and the four closed-weight LLMs, which also count among the LLMs, each without (U) and with (C) the list of expected cell types. Shares are averaged over the cell types of a dataset, then over the four datasets and the methods of a row. All methods ran on the 1,000-cell test sets.

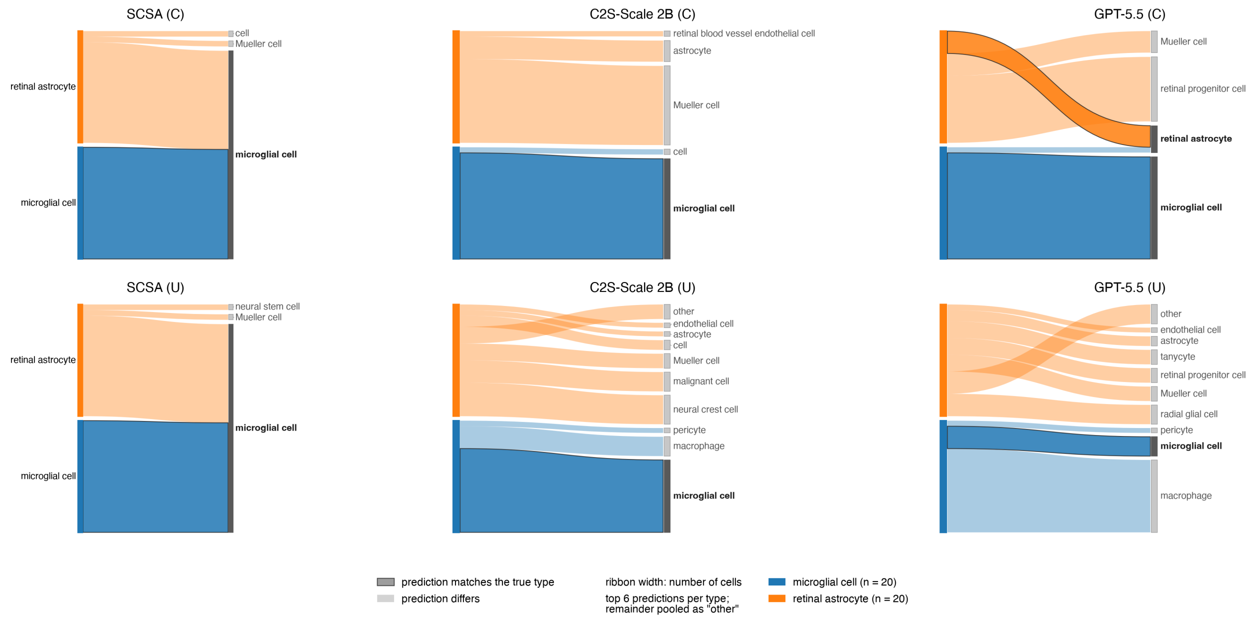


**Figure S11 Predicted labels for the cells of the two rare cell types in HFRA.** Each alluvial diagram follows the cells of the two cell types in HFRA below 1% of the full dataset, from their true type on the left to the labels one method assigned on the right. Ribbon width is proportional to the number of cells, and each true type has its own color. A right-hand label is dark gray and set in bold where it names one of the true types, and a ribbon that ends at its own true type is opaque with a dark edge. For each true type, the six most frequent labels are drawn separately and the rest pooled as "other". The top row shows runs with the list of expected cell types (C) and the bottom row runs without it (U). The columns are SCSA, C2S-Scale 2B and GPT-5.5, for each family the method whose constrained run had the highest balanced accuracy averaged over the four datasets. True and predicted labels are Cell Ontology terms after the mapping used for scoring, so a label on both sides names the same term. All methods ran on the 1,000-cell test set.

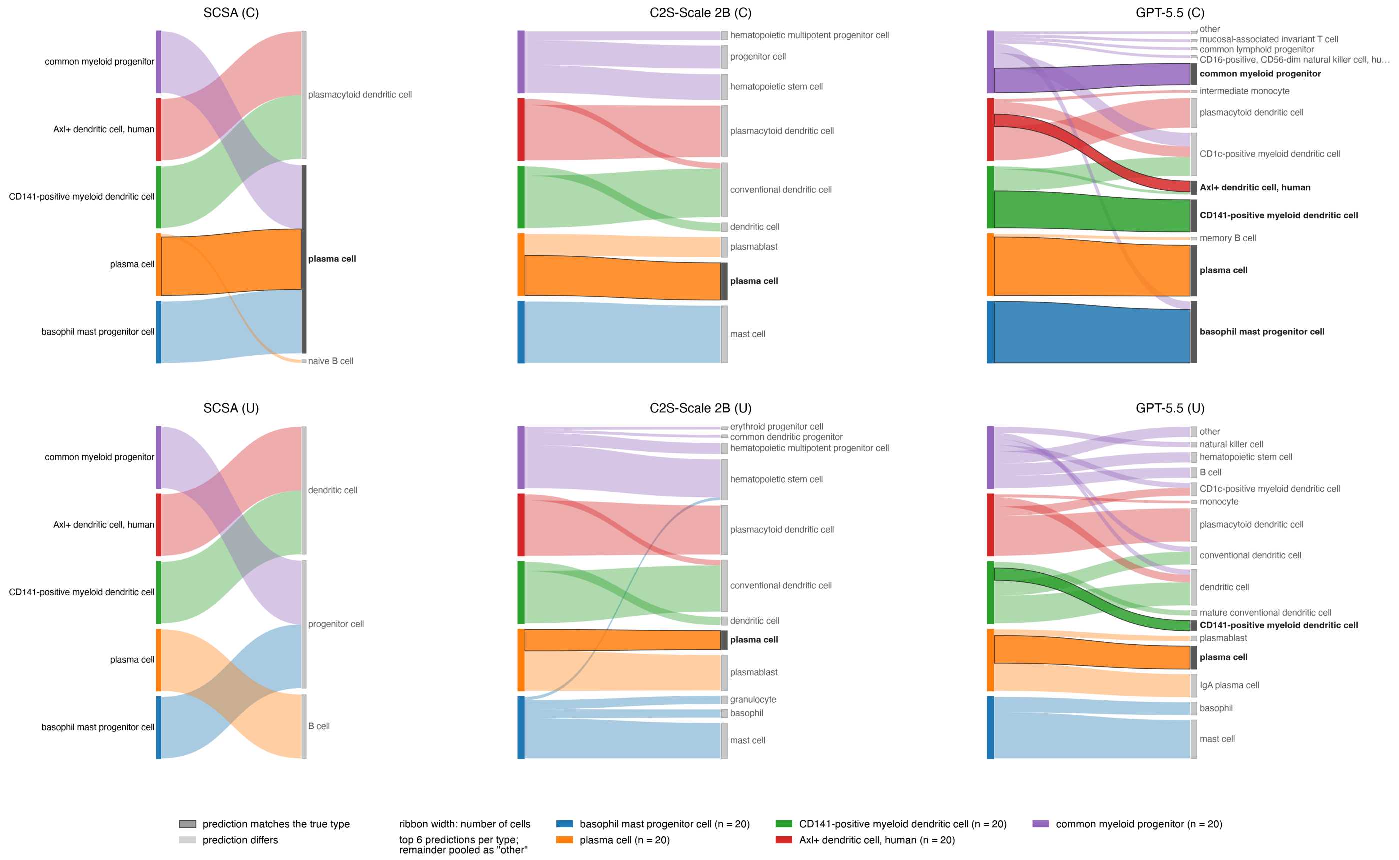


**Figure S12 Predicted labels for the cells of the five rarest cell types in HIHA.** Each alluvial diagram follows the cells of the five rarest cell types in HIHA, all below 1% of the full dataset, from their true type on the left to the labels one method assigned on the right. Ribbon width is proportional to the number of cells, and each true type has its own color. A right-hand label is dark gray and set in bold where it names one of the true types, and a ribbon that ends at its own true type is opaque with a dark edge. For each true type, the six most frequent labels are drawn separately and the rest pooled as "other". The top row shows runs with the list of expected cell types (C) and the bottom row runs without it (U). The columns are SCSA, C2S-Scale 2B and GPT-5.5, for each family the method whose constrained run had the highest balanced accuracy averaged over the four datasets. True and predicted labels are Cell Ontology terms after the mapping used for scoring, so a label on both sides names the same term. All methods ran on the 1,000-cell test set.

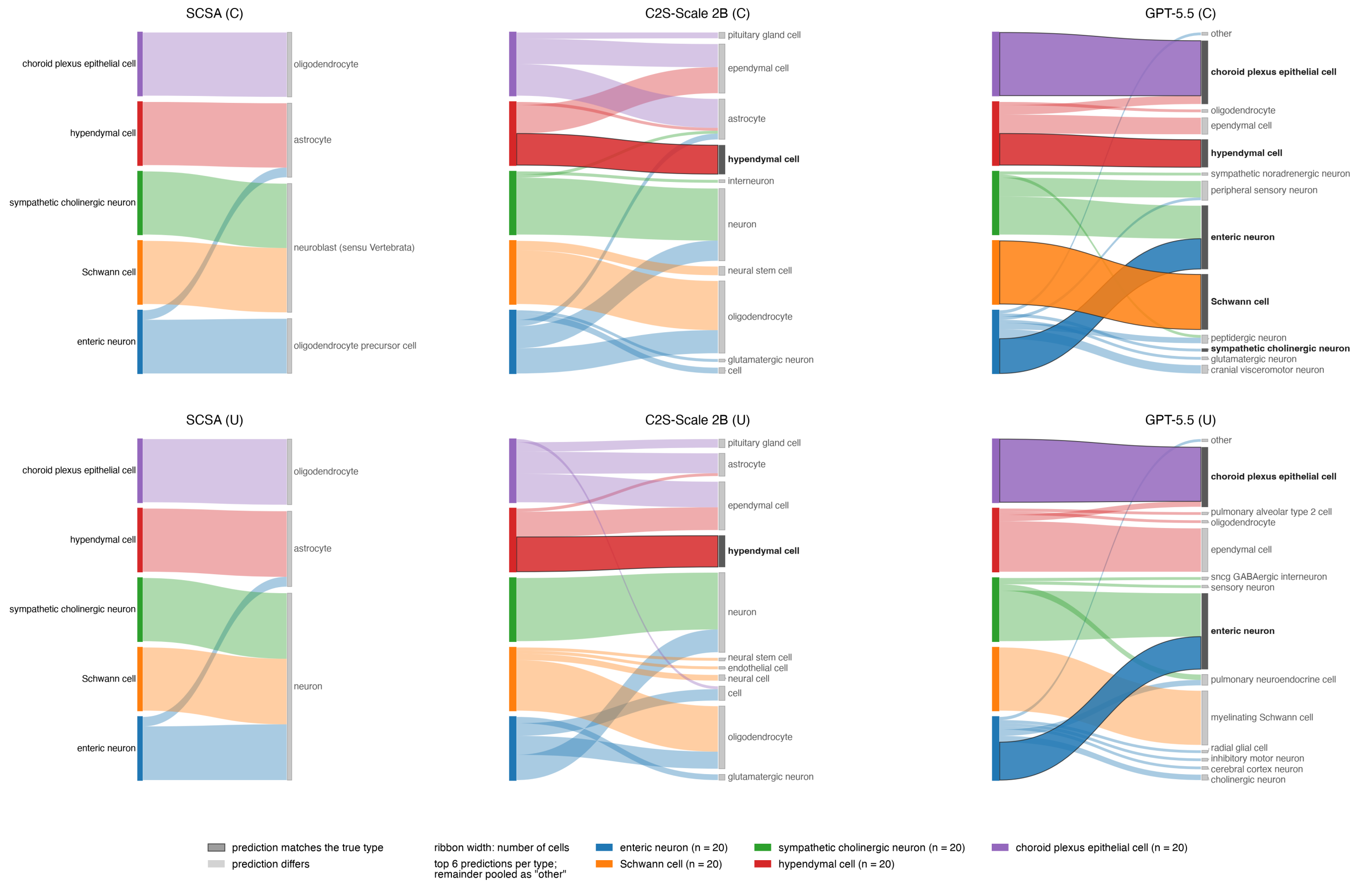


**Figure S13 Predicted labels for the cells of the five rarest cell types in Zeisel.** Each alluvial diagram follows the cells of the five rarest cell types in Zeisel, all below 1% of the full dataset, from their true type on the left to the labels one method assigned on the right. Ribbon width is proportional to the number of cells, and each true type has its own color. A right-hand label is dark gray and set in bold where it names one of the true types, and a ribbon that ends at its own true type is opaque with a dark edge. For each true type, the six most frequent labels are drawn separately and the rest pooled as "other". The top row shows runs with the list of expected cell types (C) and the bottom row runs without it (U). The columns are SCSA, C2S-Scale 2B and GPT-5.5, for each family the method whose constrained run had the highest balanced accuracy averaged over the four datasets. True and predicted labels are Cell Ontology terms after the mapping used for scoring, so a label on both sides names the same term. All methods ran on the 1,000-cell test set.

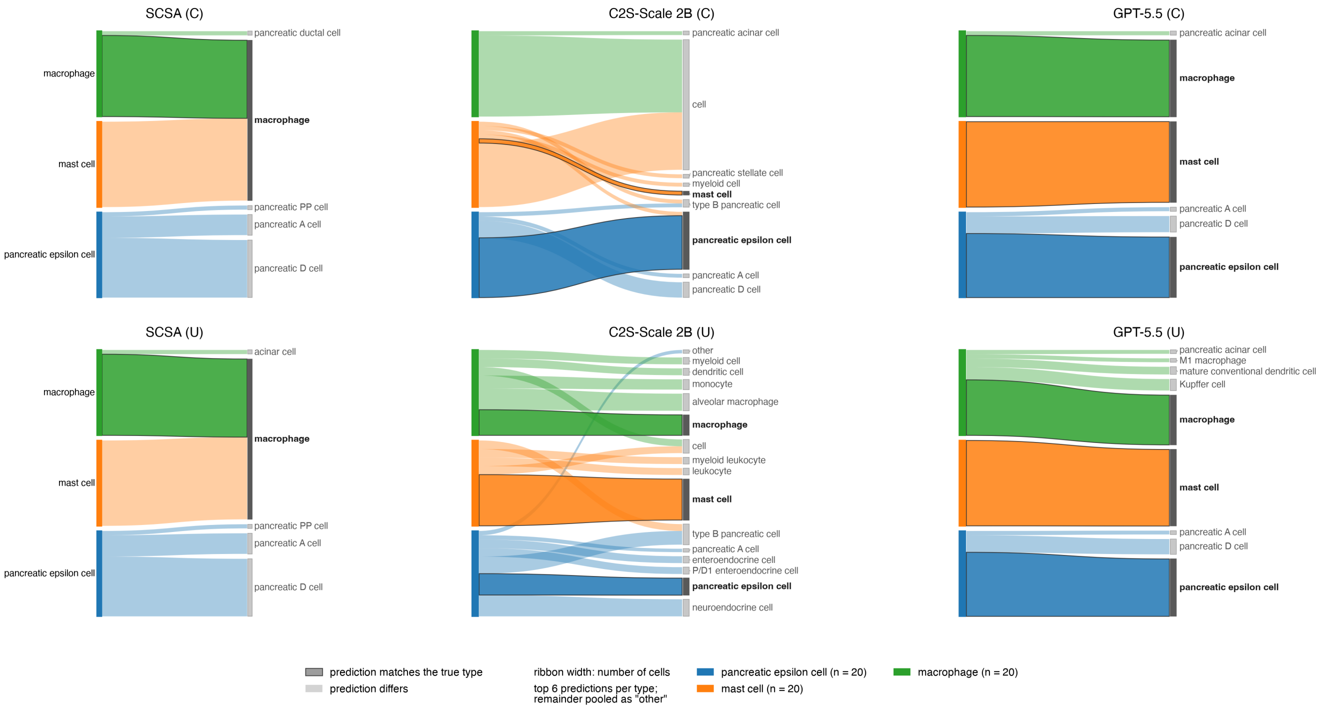


**Figure S14 Predicted labels for the cells of the three rare cell types in human pancreas.** Each alluvial diagram follows the cells of the three cell types in the human pancreas data below 1% of the full dataset, from their true type on the left to the labels one method assigned on the right. Ribbon width is proportional to the number of cells, and each true type has its own color. A right-hand label is dark gray and set in bold where it names one of the true types, and a ribbon that ends at its own true type is opaque with a dark edge. For each true type, the six most frequent labels are drawn separately and the rest pooled as "other". The top row shows runs with the list of expected cell types (C) and the bottom row runs without it (U). The columns are SCSA, C2S-Scale 2B and GPT-5.5, for each family the method whose constrained run had the highest balanced accuracy averaged over the four datasets. True and predicted labels are Cell Ontology terms after the mapping used for scoring, so a label on both sides names the same term. All methods ran on the 1,000-cell test set.

#### Supplementary Tables

| | HFRA | HIHA | Zeisel | Human pancreas |
|---|---|---|---|---|
| Source | Kriukov et al. (2025) | Gong et al. (2025) | Zeisel et al. (2018) | Luecken et al. (2022) |
| Organism | human | human | mouse | human |
| Tissue | fetal retina | peripheral blood | nervous system (19 regions) | pancreas |
| Donors | 22 (9 to 27 weeks post-fertilization) | 108 (11 to 65 years) | – | – |
| Technology | 10x 3′ v1 to v3 | 10x 3′ v3 | 10x Chromium | inDrop, CEL-seq, CEL-seq2, Smart-seq2 |
| Cells as downloaded | 108,838 | 1,821,725 | 160,796 | 16,382 |
| Cells after quality control | 102,997 | 1,742,354 | 145,837 | 11,704 |
| Genes as downloaded | 32,055 | 32,357 | 27,998 | 19,093 |
| Genes after quality control | 21,604 | 25,234 | 20,316 | 14,970 |
| Cell types | 9 | 38 | 39 | 14 |
| Cell types scored | 9 | 38 | 29 | 11 |
| Labels | Cell Ontology | Cell Ontology | authors' | curators' |
| License | CC BY 4.0 | CC BY 4.0 | none stated | CC BY 4.0 |
| Test sets (cells) | 1,000 and 10,000 | 999 and 9,999 | 999 and 9,997 | 999 and 10,000 |

**Table S1 Datasets of the benchmark.** Cells and genes of the files as downloaded and after quality control. Cell types gives the number of distinct labels of a file. Cell types scored counts those left after the labels were mapped to the Cell Ontology, which merged some of the labels (Zeisel 39 to 29, human pancreas 14 to 13), and that have at least 20 cells. A test set draws each scored cell type in proportion to its share of the dataset, with at least 20 cells. HFRA compiles Lu et al. (2020) and Sridhar et al. (2020), and human pancreas compiles six studies, Baron et al. (2016), Muraro et al. (2016), Grün et al. (2016), Segerstolpe et al. (2016), Xin et al. (2016) and Lawlor et al. (2017). HFRA is the Human Fetal Retina Atlas and HIHA the Human Immune Health Atlas.

| Method | Subcategory | Reasoning | License | Parameters | Available | Publication |
|---|---|---|---|---|---|---|
| *Baselines* | | | | | | |
| SCSA | marker-set scoring | – | GPL-3.0 | – | 2019-08 | Front. Genet. 2020-05 |
| scType | marker-set scoring | – | GPL-3.0 | – | 2021-07 | Nat. Commun. 2022-03 |
| SingleR | reference-based classifier | – | GPL-3.0 | – | 2018-03 | Nat. Immunol. 2019-01 |
| CellTypist | reference-based classifier | – | MIT | 23.3K–212K per dataset | 2021-05 | Science 2022-05 |
| *Single-cell language models* | | | | | | |
| InstructCell | generative (free text) | – | MIT | 496M (TE 110M, TD 113M, EE 168M, ED 105M) | 2025-01 | preprint 2025-01 |
| C2S-Scale 2B | generative (free text) | – | CC BY 4.0, Gemma Terms of Use | 2.61B (TD) | 2025-10 | preprint 2025-04 |
| C2S-Scale 27B | generative (free text) | – | CC BY 4.0, Gemma Terms of Use | 27.2B (TD) | 2025-10 | preprint 2025-04 |
| LangCell | embedding (closed set) | – | MIT | 173M (TE 133M, EE 39.3M, heads 330K) | 2024-05 | ICML 2024-07 |
| CellWhisperer | embedding (closed set) | – | GPL-3.0 (code) | 161M (TE 108M, EE 39.3M, heads 13.7M) | 2024-10 | Nat. Biotechnol. 2025-11 |
| mmContext | embedding (closed set) | – | MIT (code) | 125M (TE 109M, EE 12.3M, heads 2.89M) | 2025-11 | Bioinformatics 2026-05 |
| *General-purpose LLMs* | | | | | | |
| Claude Sonnet 4.6 | closed-weight reasoning | 10,000 tokens | proprietary (API) | not disclosed | 2026-02 | – |
| Gemini 3.1 Pro | closed-weight reasoning | 10,000 tokens | proprietary (API) | not disclosed | 2026-02 | – |
| Claude Opus 4.7 | closed-weight reasoning | 10,000 tokens | proprietary (API) | not disclosed | 2026-04 | – |
| GPT-5.5 | closed-weight reasoning | xhigh | proprietary (API) | not disclosed | 2026-04 | – |
| GPT-OSS-120B | open-weight reasoning | medium | Apache-2.0 | 117B (5.1B active) | 2025-08 | – |
| GPT-OSS-20B | open-weight reasoning | medium | Apache-2.0 | 21B (3.6B active) | 2025-08 | – |
| DeepSeek-V3.2 | open-weight reasoning | medium | MIT | 671B (37B active) | 2025-12 | – |
| Kimi-K2.6 | open-weight reasoning | medium | Modified MIT | 1T (32B active) | 2026-04 | – |
| Llama-3.1-70B | open-weight non-reasoning | none | Llama 3.1 Community License | 70.6B | 2024-07 | – |
| Llama-3.3-70B | open-weight non-reasoning | none | Llama 3.3 Community License | 70.6B | 2024-12 | – |
| Llama-4-Maverick | open-weight non-reasoning | none | Llama 4 Community License | 400B (17B active) | 2025-04 | – |

**Table S2 Annotation methods.** Method names the methods as in the text, and Gemini 3.1 Pro stands for the preview release Gemini 3.1 Pro Preview. Subcategory groups the methods as in Figure 1B. Reasoning gives the setting sent with each request to a general-purpose LLM through the OpenRouter API, an effort level or, for the Anthropic and Google models, a budget of reasoning tokens, which OpenRouter passes on to the provider (for Gemini 3.1 Pro, Google maps the budget to a thinking level). It is none for the models that do not reason. License refers to the published weights or, where the weights state none, to the code (marked code). The proprietary models are used only through their API, and C2S-Scale, fine-tuned from Gemma 2, is also subject to the Gemma Terms of Use. Parameters count the learned weights of the checkpoint used here, with tied weights counted once. TE is the text encoder and TD the text decoder. C2S-Scale is decoder-only, and InstructCell's encoder and decoder share their word embeddings, which are counted with the TE. EE is the expression encoder, for mmContext an adapter on the log-normalized expression of 10,000 preselected genes and for CellWhisperer a frozen Geneformer. ED is the expression decoder, which InstructCell uses only to generate expression profiles, and heads are the projection and matching heads. CellTypist uses one logistic-regression model per dataset. For mixture-of-experts LLMs, the developer's total is followed by the parameters active per token in parentheses. Available gives the month in which the weights, software or API became public. Publication gives the journal or conference and the month of the peer-reviewed paper, or the month of the preprint. Dashes mark fields that do not apply.

| | HFRA | HIHA | Zeisel | Human pancreas |
|---|---|---|---|---|
| CellTypist model | Fetal_Human_Retina | Immune_All_High | Mouse_Isocortex_Hippocampus | Adult_Human_PancreaticIslet |
| SingleR reference | CELLxGENE, Zuo et al. (2024) | celldex MonacoImmuneData, Monaco et al. (2019) | CELLxGENE, Yao et al. (2021) | CELLxGENE, Xu et al. (2023) |
| scType tissue | Eye | Immune system | Brain | Pancreas |
| SCSA tissue | Retina | Peripheral blood | Brain | Pancreas |

**Table S3 Inputs of the baselines.** CellTypist model names the pretrained model CellTypist used. SingleR reference gives the annotated data SingleR compared each cell with, a CELLxGENE dataset or the MonacoImmuneData reference of the celldex package. scType tissue and SCSA tissue name the tissue whose marker genes scType and SCSA scored. HFRA is the Human Fetal Retina Atlas and HIHA the Human Immune Health Atlas.

| Metric | Definition | Chance |
|---|---|---|
| *Exact match* | | |
| Balanced accuracy | Mean over the true cell types of the share of their cells labeled correctly | $1/K$ |
| Accuracy | Share of cells labeled correctly | $\sum_k \pi_k^2$ |
| Macro F1 | Mean over the true cell types of the F1 score, the harmonic mean of precision and recall | $1/K$ |
| Weighted F1 | Mean of the F1 scores of the true cell types, weighted by their numbers of cells | $\sum_k \pi_k^2$ |
| Per-class recall | Share of the cells of one cell type labeled correctly | $\pi_k$ |
| *Ontology-aware* | | |
| Agreement score | 1 for the true type, 0.5 for a coarser type that is not broad or for a finer type (see answer type), and 0 otherwise, averaged per cell type and then over cell types. Adapted from Hou and Ji (2024), who judge the partial match by hand-assigned broad cell types | – |
| Rada similarity | $1/(1+d)$, with $d$ the number of is_a edges on the shortest path between the true and the predicted term, the distance of Rada et al. (1989), averaged over cells | – |
| Lin similarity | $2\,\mathrm{IC}(m)/(\mathrm{IC}(a)+\mathrm{IC}(b))$ (Lin, 1998), with $m$ the most informative common ancestor of the terms $a$ and $b$, and the intrinsic information content $\mathrm{IC}(t) = 1 - \log n_t / \log N$ of Seco et al. (2004), with $n_t$ the number of terms at or below $t$ and $N$ the number of terms in the hierarchy, averaged over cells | – |
| *Text similarity* | | |
| Text, mapped | Similarity in meaning between the answer after the curated mapping and the Cell Ontology name of the true type | – |
| Text, raw | The same for the answer as written, with Cell Ontology identifiers read as names. Needs no mapping of the answers | – |
| *Diagnostics* | | |
| Coverage | Share of cells whose answer as written is a Cell Ontology term or has a reviewed mapping. A run needs 0.9 to be scored | – |
| Answers needing the curated mapping | Share of cells whose answer as written is neither a Cell Ontology term nor one of the dataset's own labels | – |
| Balanced accuracy without the curated mapping | Balanced accuracy when only the answers that need no curated mapping count as correct | – |
| Answer type | Where the answer falls relative to the true type. It is the exact type, a coarser type (an ancestor), broad if more than 250 terms lie at or below it and otherwise specific, a finer type (a descendant), another cell type, or no usable answer, one that resolves to no term below 'cell'. Shares per cell type, averaged over cell types | – |
| Rare-type outcome | Whether a method labeled a rare cell type correctly for at least half of its cells, for fewer, only on other cells, or never | – |
| Distinct labels | Number of different answers as written, and of different cell types after the curated mapping | – |
| Answer format | Shares of cells whose answer as written matches no Cell Ontology term (non-CL), is empty, or runs past eight words | – |

**Table S4 Evaluation metrics.** Metrics range from 0 to 1, higher being better. $K$ is the number of cell types in a test set and $\pi_k$ the share of cell type $k$. Chance is the expected score of the random model of Supplementary Tables S5–S8, which assigns cell types in proportion to their frequency in the test set (closely approximated for the F1 scores). Its rows there give its other scores. Exact-match metrics follow scikit-learn. Ontology-aware metrics use the is_a hierarchy of the Cell Ontology (release 2026-06-08) from its root 'cell' down. An answer that resolves to no term of it scores 0, and one that resolves to 'cell', as the curated mapping does for answers naming no cell type, scores 0 on Lin and the agreement score but $1/(1+d)$ on Rada. Text similarity is the cosine between the SapBERT embeddings of the two labels (Liu et al., 2021, using the [CLS] token), averaged per cell type and then over cell types. The mean $s$ is rescaled as in BERTScore (Zhang et al., 2020) to $(s-b)/(1-b)$ and clipped to $[0,1]$, with $b$ the mean cosine over all pairs of different cell type names in the hierarchy, one value for all runs. An answer that is empty or 'cell', or in the mapped variant maps to 'cell', gets the cosine $b$, which rescales to 0.

| | | | Exact match | | | | Ontology-aware | | | Text similarity | | Curated mapping | | Distinct labels | |
|---|---|---|---|---|---|---|---|---|---|---|---|---|---|---|---|
| Family | Method | Setting | Bal. acc. | Acc. | Macro F1 | Wtd. F1 | Agree. | Rada | Lin | Mapped | Raw | Needed | Bal. acc. without | Mapped | Raw |
| **1,000 cells** | | | | | | | | | | | | | | | |
| LLM | Gemini 3.1 Pro | C | **0.594** | **0.654** | 0.542 | 0.608 | **0.594** | **0.728** | **0.715** | 0.771 | 0.771 | 0.000 | 0.594 | 9 | 9 |
| LLM | GPT-5.5 | C | 0.590 | 0.647 | **0.552** | **0.628** | 0.590 | 0.724 | 0.711 | **0.775** | **0.775** | 0.019 | 0.590 | 9 | 10 |
| LLM | Claude Sonnet 4.6 | C | 0.561 | 0.613 | 0.506 | 0.589 | 0.561 | 0.696 | 0.676 | 0.761 | 0.761 | 0.001 | 0.561 | 8 | 9 |
| LLM | Kimi-K2.6 | C | 0.548 | 0.470 | 0.518 | 0.509 | 0.548 | 0.590 | 0.530 | 0.673 | 0.673 | 0.198 | 0.548 | 9 | 11 |
| baseline | SCSA | C | 0.534 | 0.477 | 0.455 | 0.514 | 0.534 | 0.600 | 0.554 | 0.708 | 0.685 | 0.144 | 0.534 | 8 | 9 |
| LLM | GPT-OSS-120B | C | 0.529 | 0.517 | 0.469 | 0.489 | 0.529 | 0.618 | 0.584 | 0.743 | 0.742 | 0.004 | 0.528 | 9 | 12 |
| LLM | Claude Opus 4.7 | C | 0.526 | 0.617 | 0.459 | 0.564 | 0.526 | 0.695 | 0.683 | 0.740 | 0.740 | 0.002 | 0.526 | 9 | 11 |
| baseline | SingleR | C | 0.491 | 0.609 | 0.409 | 0.599 | 0.491 | 0.697 | 0.693 | 0.704 | 0.704 | 0.000 | 0.491 | 7 | 7 |
| LLM | DeepSeek-V3.2 | C | 0.483 | 0.593 | 0.453 | 0.546 | 0.483 | 0.676 | 0.643 | 0.717 | 0.717 | 0.001 | 0.483 | 8 | 9 |
| scLM | LangCell | C | 0.469 | 0.457 | 0.391 | 0.450 | 0.469 | 0.590 | 0.628 | 0.736 | 0.736 | 0.000 | 0.469 | 8 | 8 |
| LLM | Llama-4-Maverick | C | 0.469 | 0.430 | 0.373 | 0.414 | 0.469 | 0.549 | 0.546 | 0.721 | 0.721 | 0.000 | 0.469 | 8 | 8 |
| scLM | C2S-Scale 2B | C | 0.468 | 0.346 | 0.412 | 0.312 | 0.479 | 0.488 | 0.450 | 0.733 | 0.734 | 0.006 | 0.468 | 14 | 15 |
| LLM | GPT-OSS-20B | C | 0.451 | 0.475 | 0.416 | 0.464 | 0.451 | 0.587 | 0.562 | 0.710 | 0.713 | 0.017 | 0.450 | 9 | 15 |
| baseline | SingleR | U | 0.432 | 0.528 | 0.403 | 0.556 | 0.461 | 0.650 | 0.668 | 0.686 | 0.686 | 0.000 | 0.432 | 25 | 25 |
| scLM | CellWhisperer | C | 0.424 | 0.600 | 0.372 | 0.512 | 0.424 | 0.690 | 0.708 | 0.709 | 0.709 | 0.000 | 0.424 | 6 | 6 |
| baseline | scType | C | 0.422 | 0.485 | 0.369 | 0.506 | 0.422 | 0.615 | 0.602 | 0.640 | 0.627 | 0.067 | 0.422 | 7 | 8 |
| baseline | CellTypist | C | 0.418 | 0.429 | 0.339 | 0.398 | 0.418 | 0.559 | 0.566 | 0.681 | 0.681 | 0.000 | 0.418 | 6 | 6 |
| baseline | CellTypist | U | 0.399 | 0.400 | 0.373 | 0.429 | 0.501 | 0.543 | 0.600 | 0.724 | 0.189 | 0.998 | 0.000 | 10 | 23 |
| scLM | C2S-Scale 27B | C | 0.376 | 0.225 | 0.248 | 0.189 | 0.376 | 0.392 | 0.390 | 0.689 | 0.689 | 0.011 | 0.376 | 15 | 18 |
| baseline | SCSA | U | 0.362 | 0.315 | 0.357 | 0.395 | 0.383 | 0.476 | 0.440 | 0.583 | 0.613 | 0.759 | 0.261 | 12 | 13 |
| LLM | Gemini 3.1 Pro | U | 0.359 | 0.394 | 0.387 | 0.455 | 0.440 | 0.543 | 0.537 | 0.631 | 0.633 | 0.126 | 0.323 | 34 | 43 |
| LLM | Llama-3.1-70B | C | 0.358 | 0.218 | 0.219 | 0.201 | 0.358 | 0.386 | 0.381 | 0.687 | 0.688 | 0.003 | 0.358 | 10 | 11 |
| LLM | Llama-3.3-70B | C | 0.342 | 0.245 | 0.255 | 0.249 | 0.342 | 0.405 | 0.374 | 0.686 | 0.686 | 0.000 | 0.342 | 8 | 8 |
| LLM | GPT-5.5 | U | 0.330 | 0.339 | 0.361 | 0.410 | 0.414 | 0.502 | 0.500 | 0.602 | 0.604 | 0.204 | 0.320 | 62 | 70 |
| scLM | C2S-Scale 27B | U | 0.296 | 0.238 | 0.263 | 0.236 | 0.318 | 0.412 | 0.379 | 0.566 | 0.567 | 0.029 | 0.296 | 48 | 56 |
| scLM | C2S-Scale 2B | U | 0.295 | 0.224 | 0.321 | 0.238 | 0.314 | 0.406 | 0.428 | 0.522 | 0.525 | 0.030 | 0.295 | 38 | 39 |
| baseline | scType | U | 0.291 | 0.347 | 0.294 | 0.404 | 0.352 | 0.523 | 0.543 | 0.602 | 0.544 | 1.000 | 0.000 | 10 | 11 |
| scLM | mmContext | C | 0.286 | 0.360 | 0.214 | 0.305 | 0.286 | 0.492 | 0.463 | 0.640 | 0.640 | 0.000 | 0.286 | 9 | 9 |
| LLM | Kimi-K2.6 | U | 0.251 | 0.211 | 0.294 | 0.260 | 0.277 | 0.408 | 0.305 | 0.383 | 0.374 | 0.639 | 0.118 | 54 | 67 |
| LLM | Claude Sonnet 4.6 | U | 0.241 | 0.258 | 0.257 | 0.303 | 0.340 | 0.443 | 0.450 | 0.563 | 0.561 | 0.133 | 0.206 | 50 | 61 |
| LLM | Claude Opus 4.7 | U | 0.240 | 0.182 | 0.258 | 0.235 | 0.281 | 0.391 | 0.385 | 0.540 | 0.514 | 0.212 | 0.206 | 43 | 105 |
| LLM | DeepSeek-V3.2 | U | 0.227 | 0.187 | 0.279 | 0.227 | 0.300 | 0.387 | 0.384 | 0.499 | 0.492 | 0.215 | 0.099 | 58 | 78 |
| LLM | GPT-OSS-120B | U | 0.194 | 0.160 | 0.239 | 0.188 | 0.264 | 0.373 | 0.371 | 0.452 | 0.442 | 0.334 | 0.056 | 49 | 63 |
| LLM | GPT-OSS-20B | U | 0.127 | 0.114 | 0.173 | 0.150 | 0.171 | 0.328 | 0.315 | 0.359 | 0.357 | 0.291 | 0.038 | 74 | 124 |
| chance | Random Stratified | – | 0.112 | 0.237 | 0.112 | 0.235 | 0.112 | 0.392 | 0.356 | 0.515 | 0.515 | 0.000 | 0.112 | 9 | 9 |
| scLM | CellWhisperer | U | 0.083 | 0.135 | 0.092 | 0.180 | 0.111 | 0.349 | 0.406 | 0.411 | 0.411 | 0.000 | 0.083 | 43 | 43 |
| LLM | Llama-4-Maverick | U | 0.068 | 0.074 | 0.091 | 0.096 | 0.106 | 0.307 | 0.293 | 0.303 | 0.304 | 0.438 | 0.064 | 32 | 47 |
| scLM | mmContext | U | 0.060 | 0.102 | 0.089 | 0.141 | 0.117 | 0.281 | 0.295 | 0.400 | 0.400 | 0.000 | 0.060 | 237 | 238 |
| LLM | Llama-3.1-70B | U | 0.009 | 0.005 | 0.017 | 0.010 | 0.042 | 0.249 | 0.231 | 0.199 | 0.200 | 0.049 | 0.007 | 60 | 74 |
| LLM | Llama-3.3-70B | U | 0.001 | 0.001 | 0.002 | 0.002 | 0.048 | 0.251 | 0.231 | 0.211 | 0.210 | 0.018 | 0.001 | 18 | 28 |
| scLM | InstructCell | U | 0.000 | 0.000 | 0.000 | 0.000 | 0.044 | 0.214 | 0.042 | 0.070 | 0.000 | 0.788 | 0.000 | 12 | 27 |
| scLM | InstructCell | C | 0.000 | 0.000 | 0.000 | 0.000 | 0.046 | 0.207 | 0.038 | 0.063 | 0.000 | 0.731 | 0.000 | 14 | 26 |
| scLM | LangCell | U | 0.000 | 0.000 | 0.000 | 0.000 | 0.069 | 0.234 | 0.249 | 0.276 | 0.272 | 0.000 | 0.000 | 95 | 96 |

*continued on the next page*

*HFRA, continued*

| | | | Exact match | | | | Ontology-aware | | | Text similarity | | Curated mapping | | Distinct labels | |
|---|---|---|---|---|---|---|---|---|---|---|---|---|---|---|---|
| Family | Method | Setting | Bal. acc. | Acc. | Macro F1 | Wtd. F1 | Agree. | Rada | Lin | Mapped | Raw | Needed | Bal. acc. without | Mapped | Raw |
| **10,000 cells** | | | | | | | | | | | | | | | |
| baseline | SCSA | C | **0.552** | 0.476 | **0.443** | 0.509 | **0.552** | 0.597 | 0.545 | 0.721 | 0.701 | 0.139 | 0.552 | 8 | 9 |
| baseline | SingleR | C | 0.500 | **0.640** | 0.431 | **0.641** | 0.500 | **0.722** | **0.722** | 0.717 | 0.717 | 0.000 | 0.500 | 7 | 7 |
| scLM | C2S-Scale 2B | C | 0.475 | 0.345 | 0.415 | 0.305 | 0.485 | 0.485 | 0.445 | **0.733** | **0.735** | 0.005 | 0.475 | 13 | 14 |
| scLM | LangCell | C | 0.448 | 0.438 | 0.386 | 0.437 | 0.448 | 0.572 | 0.605 | 0.724 | 0.724 | 0.000 | 0.448 | 8 | 8 |
| baseline | SingleR | U | 0.435 | 0.547 | 0.415 | 0.584 | 0.469 | 0.668 | 0.688 | 0.692 | 0.692 | 0.000 | 0.435 | 27 | 27 |
| baseline | scType | C | 0.422 | 0.502 | 0.376 | 0.534 | 0.422 | 0.628 | 0.614 | 0.640 | 0.624 | 0.077 | 0.422 | 7 | 8 |
| baseline | CellTypist | C | 0.421 | 0.413 | 0.338 | 0.384 | 0.421 | 0.545 | 0.551 | 0.683 | 0.683 | 0.000 | 0.421 | 6 | 6 |
| scLM | CellWhisperer | C | 0.409 | 0.594 | 0.357 | 0.506 | 0.409 | 0.684 | 0.701 | 0.700 | 0.700 | 0.000 | 0.409 | 6 | 6 |
| baseline | CellTypist | U | 0.400 | 0.383 | 0.368 | 0.412 | 0.500 | 0.529 | 0.585 | 0.726 | 0.196 | 1.000 | 0.000 | 10 | 23 |
| baseline | SCSA | U | 0.372 | 0.303 | 0.341 | 0.385 | 0.389 | 0.464 | 0.420 | 0.579 | 0.608 | 0.785 | 0.268 | 12 | 13 |
| scLM | C2S-Scale 27B | C | 0.365 | 0.216 | 0.201 | 0.188 | 0.366 | 0.386 | 0.385 | 0.686 | 0.686 | 0.006 | 0.364 | 22 | 27 |
| scLM | C2S-Scale 2B | U | 0.315 | 0.234 | 0.340 | 0.247 | 0.329 | 0.414 | 0.435 | 0.537 | 0.540 | 0.029 | 0.315 | 43 | 44 |
| scLM | C2S-Scale 27B | U | 0.307 | 0.251 | 0.245 | 0.251 | 0.329 | 0.421 | 0.385 | 0.568 | 0.568 | 0.020 | 0.307 | 89 | 130 |
| baseline | scType | U | 0.300 | 0.368 | 0.307 | 0.433 | 0.352 | 0.536 | 0.556 | 0.604 | 0.545 | 1.000 | 0.000 | 10 | 11 |
| scLM | mmContext | C | 0.285 | 0.345 | 0.142 | 0.292 | 0.285 | 0.477 | 0.439 | 0.639 | 0.639 | 0.000 | 0.285 | 9 | 9 |
| chance | Random Stratified | – | 0.110 | 0.237 | 0.110 | 0.237 | 0.110 | 0.391 | 0.355 | 0.515 | 0.515 | 0.000 | 0.110 | 9 | 9 |
| scLM | CellWhisperer | U | 0.076 | 0.142 | 0.083 | 0.185 | 0.100 | 0.354 | 0.402 | 0.390 | 0.390 | 0.000 | 0.076 | 71 | 71 |
| scLM | mmContext | U | 0.062 | 0.103 | 0.084 | 0.140 | 0.107 | 0.281 | 0.278 | 0.393 | 0.393 | 0.000 | 0.062 | 667 | 676 |
| scLM | InstructCell | U | 0.000 | 0.000 | 0.000 | 0.000 | 0.049 | 0.213 | 0.030 | 0.068 | 0.000 | 0.783 | 0.000 | 15 | 70 |
| scLM | InstructCell | C | 0.000 | 0.000 | 0.000 | 0.000 | 0.054 | 0.207 | 0.025 | 0.063 | 0.000 | 0.735 | 0.000 | 16 | 59 |



Table S5. **Benchmark results for HFRA.** One row per method and setting, 43 at 1,000 cells and 20 at 10,000 cells, sorted by balanced accuracy within each size. The general-purpose LLMs and LangCell (U) ran at 1,000 cells only. *Family* and *Method* name the method of a run, and *Setting* is C for a run constrained by the list of expected cell types and U for a run without it. The random model, which assigns cell types in proportion to their frequency, runs in one setting. The best value of each metric at each size is highlighted in bold. *Bal. acc.* is balanced accuracy, *Acc.* accuracy, *Wtd. F1* weighted F1 and *Agree.* the agreement score (Supplementary Table S4). Text similarity uses the answer after the curated mapping (*Mapped*) or as written (*Raw*). *Needed* is the share of cells whose answer needs the curated mapping, and *Bal. acc. without* is the balanced accuracy when only the answers that need no curated mapping count as correct. *Distinct labels* counts the cell types named after the curated mapping (*Mapped*) and the different answers as written (*Raw*). Every answer was a Cell Ontology term or had a reviewed mapping.

| | | | Exact match | | | | Ontology-aware | | | Text similarity | | Curated mapping | | Distinct labels | |
|---|---|---|---|---|---|---|---|---|---|---|---|---|---|---|---|
| Family | Method | Setting | Bal. acc. | Acc. | Macro F1 | Wtd. F1 | Agree. | Rada | Lin | Mapped | Raw | Needed | Bal. acc. without | Mapped | Raw |
| **1,000 cells** | | | | | | | | | | | | | | | |
| LLM | GPT-5.5 | C | **0.405** | 0.461 | **0.366** | **0.416** | 0.454 | 0.588 | 0.783 | 0.653 | 0.652 | 0.001 | 0.405 | 30 | 31 |
| scLM | mmContext | C | **0.405** | 0.400 | 0.346 | 0.358 | 0.464 | 0.555 | 0.789 | **0.687** | **0.687** | 0.000 | 0.405 | 32 | 32 |
| LLM | Claude Opus 4.7 | C | **0.405** | 0.480 | 0.345 | 0.397 | 0.450 | **0.607** | 0.794 | 0.670 | 0.671 | 0.000 | 0.405 | 32 | 32 |
| LLM | Gemini 3.1 Pro | C | 0.392 | **0.482** | 0.343 | 0.410 | 0.434 | 0.606 | 0.795 | 0.650 | 0.651 | 0.000 | 0.392 | 31 | 31 |
| scLM | LangCell | C | 0.385 | 0.444 | 0.335 | 0.384 | 0.461 | 0.584 | 0.775 | 0.621 | 0.621 | 0.000 | 0.385 | 35 | 35 |
| LLM | Claude Sonnet 4.6 | C | 0.350 | 0.440 | 0.279 | 0.331 | 0.381 | 0.567 | 0.753 | 0.618 | 0.619 | 0.000 | 0.350 | 30 | 30 |
| LLM | GPT-OSS-120B | C | 0.337 | 0.431 | 0.283 | 0.346 | 0.392 | 0.561 | 0.750 | 0.592 | 0.592 | 0.005 | 0.337 | 29 | 31 |
| LLM | DeepSeek-V3.2 | C | 0.323 | 0.398 | 0.272 | 0.323 | 0.374 | 0.527 | 0.725 | 0.573 | 0.573 | 0.000 | 0.323 | 32 | 32 |
| LLM | Kimi-K2.6 | C | 0.306 | 0.348 | 0.273 | 0.313 | 0.345 | 0.476 | 0.601 | 0.512 | 0.512 | 0.211 | 0.306 | 31 | 32 |
| LLM | GPT-OSS-20B | C | 0.297 | 0.351 | 0.253 | 0.297 | 0.378 | 0.493 | 0.705 | 0.538 | 0.538 | 0.006 | 0.296 | 29 | 32 |
| scLM | C2S-Scale 2B | C | 0.294 | 0.365 | 0.262 | 0.319 | 0.414 | 0.566 | **0.801** | 0.668 | 0.669 | 0.000 | 0.294 | 40 | 41 |
| scLM | CellWhisperer | C | 0.277 | 0.300 | 0.210 | 0.241 | 0.321 | 0.467 | 0.736 | 0.598 | 0.598 | 0.000 | 0.277 | 31 | 31 |
| scLM | C2S-Scale 2B | U | 0.244 | 0.295 | 0.241 | 0.277 | 0.360 | 0.503 | 0.761 | 0.607 | 0.606 | 0.003 | 0.244 | 53 | 55 |
| baseline | SingleR | C | 0.241 | 0.279 | 0.143 | 0.205 | 0.427 | 0.446 | 0.694 | 0.452 | 0.451 | 0.000 | 0.241 | 12 | 12 |
| baseline | scType | C | 0.220 | 0.270 | 0.123 | 0.183 | 0.396 | 0.459 | 0.724 | 0.495 | 0.495 | 0.000 | 0.220 | 10 | 10 |
| LLM | Llama-4-Maverick | C | 0.200 | 0.241 | 0.155 | 0.190 | 0.255 | 0.415 | 0.683 | 0.537 | 0.536 | 0.002 | 0.200 | 20 | 22 |
| LLM | Claude Opus 4.7 | U | 0.185 | 0.219 | 0.165 | 0.204 | 0.375 | 0.446 | 0.721 | 0.541 | 0.534 | 0.030 | 0.185 | 53 | 69 |
| LLM | GPT-5.5 | U | 0.184 | 0.213 | 0.185 | 0.227 | 0.362 | 0.449 | 0.729 | 0.547 | 0.546 | 0.027 | 0.179 | 55 | 63 |
| baseline | SingleR | U | 0.177 | 0.228 | 0.136 | 0.198 | 0.336 | 0.451 | 0.725 | 0.535 | 0.535 | 0.000 | 0.177 | 27 | 27 |
| LLM | Claude Sonnet 4.6 | U | 0.169 | 0.161 | 0.149 | 0.151 | 0.303 | 0.416 | 0.693 | 0.531 | 0.535 | 0.149 | 0.161 | 48 | 59 |
| baseline | scType | U | 0.162 | 0.226 | 0.108 | 0.172 | 0.301 | 0.459 | 0.706 | 0.543 | 0.474 | 1.000 | 0.000 | 12 | 12 |
| baseline | SCSA | C | 0.157 | 0.124 | 0.062 | 0.050 | 0.294 | 0.347 | 0.648 | 0.431 | 0.431 | 0.000 | 0.157 | 10 | 10 |
| scLM | C2S-Scale 27B | U | 0.150 | 0.145 | 0.141 | 0.152 | 0.340 | 0.370 | 0.631 | 0.430 | 0.419 | 0.071 | 0.150 | 66 | 80 |
| LLM | Gemini 3.1 Pro | U | 0.148 | 0.177 | 0.128 | 0.169 | 0.365 | 0.430 | 0.732 | 0.556 | 0.556 | 0.021 | 0.142 | 40 | 46 |
| scLM | mmContext | U | 0.142 | 0.162 | 0.162 | 0.196 | 0.236 | 0.367 | 0.648 | 0.529 | 0.526 | 0.000 | 0.142 | 180 | 183 |
| LLM | GPT-OSS-120B | U | 0.139 | 0.147 | 0.116 | 0.132 | 0.336 | 0.400 | 0.694 | 0.512 | 0.509 | 0.185 | 0.110 | 39 | 80 |
| LLM | Kimi-K2.6 | U | 0.132 | 0.118 | 0.120 | 0.116 | 0.286 | 0.350 | 0.552 | 0.434 | 0.433 | 0.228 | 0.130 | 45 | 48 |
| LLM | Llama-3.3-70B | C | 0.123 | 0.179 | 0.092 | 0.134 | 0.175 | 0.344 | 0.589 | 0.366 | 0.366 | 0.000 | 0.123 | 29 | 29 |
| LLM | DeepSeek-V3.2 | U | 0.113 | 0.118 | 0.107 | 0.125 | 0.342 | 0.357 | 0.654 | 0.438 | 0.435 | 0.146 | 0.099 | 37 | 54 |
| scLM | LangCell | U | 0.113 | 0.150 | 0.128 | 0.171 | 0.210 | 0.364 | 0.616 | 0.470 | 0.470 | 0.000 | 0.113 | 121 | 122 |
| baseline | CellTypist | C | 0.101 | 0.077 | 0.026 | 0.020 | 0.350 | 0.260 | 0.545 | 0.184 | 0.184 | 0.000 | 0.101 | 4 | 4 |
| LLM | GPT-OSS-20B | U | 0.100 | 0.094 | 0.098 | 0.099 | 0.313 | 0.345 | 0.648 | 0.426 | 0.397 | 0.456 | 0.078 | 49 | 135 |
| baseline | CellTypist | U | 0.099 | 0.075 | 0.056 | 0.042 | **0.467** | 0.306 | 0.635 | 0.345 | 0.274 | 1.000 | 0.000 | 14 | 15 |
| scLM | CellWhisperer | U | 0.097 | 0.149 | 0.080 | 0.127 | 0.169 | 0.343 | 0.639 | 0.465 | 0.464 | 0.000 | 0.097 | 82 | 83 |
| LLM | Llama-3.1-70B | C | 0.093 | 0.121 | 0.080 | 0.115 | 0.163 | 0.293 | 0.551 | 0.313 | 0.312 | 0.009 | 0.093 | 40 | 48 |
| scLM | C2S-Scale 27B | C | 0.093 | 0.079 | 0.095 | 0.084 | 0.201 | 0.293 | 0.554 | 0.355 | 0.350 | 0.054 | 0.093 | 64 | 71 |
| LLM | Llama-4-Maverick | U | 0.088 | 0.067 | 0.069 | 0.052 | 0.238 | 0.313 | 0.630 | 0.473 | 0.469 | 0.028 | 0.083 | 19 | 39 |
| LLM | Llama-3.3-70B | U | 0.040 | 0.030 | 0.014 | 0.011 | 0.368 | 0.270 | 0.592 | 0.273 | 0.272 | 0.018 | 0.038 | 25 | 31 |
| LLM | Llama-3.1-70B | U | 0.037 | 0.028 | 0.030 | 0.023 | 0.269 | 0.251 | 0.542 | 0.269 | 0.259 | 0.222 | 0.032 | 41 | 54 |
| baseline | SCSA | U | 0.026 | 0.020 | 0.004 | 0.003 | 0.305 | 0.307 | 0.657 | 0.443 | 0.365 | 0.361 | 0.020 | 10 | 10 |
| chance | Random Stratified | – | 0.025 | 0.040 | 0.025 | 0.039 | 0.048 | 0.176 | 0.386 | 0.143 | 0.144 | 0.000 | 0.025 | 38 | 38 |
| scLM | InstructCell | C | 0.022 | 0.017 | 0.001 | 0.001 | 0.316 | 0.238 | 0.558 | 0.245 | 0.232 | 0.126 | 0.022 | 10 | 12 |
| scLM | InstructCell | U | 0.021 | 0.016 | 0.002 | 0.001 | 0.309 | 0.235 | 0.557 | 0.249 | 0.238 | 0.140 | 0.021 | 9 | 11 |

*continued on the next page*

*HIHA, continued*

| | | | Exact match | | | | Ontology-aware | | | Text similarity | | Curated mapping | | Distinct labels | |
|---|---|---|---|---|---|---|---|---|---|---|---|---|---|---|---|
| Family | Method | Setting | Bal. acc. | Acc. | Macro F1 | Wtd. F1 | Agree. | Rada | Lin | Mapped | Raw | Needed | Bal. acc. without | Mapped | Raw |
| **10,000 cells** | | | | | | | | | | | | | | | |
| scLM | mmContext | C | **0.383** | 0.382 | **0.310** | 0.402 | 0.449 | 0.553 | 0.810 | **0.685** | **0.686** | 0.000 | 0.383 | 35 | 35 |
| scLM | LangCell | C | 0.376 | **0.530** | 0.308 | 0.493 | 0.448 | 0.648 | 0.823 | 0.606 | 0.606 | 0.000 | 0.376 | 37 | 37 |
| scLM | C2S-Scale 2B | C | 0.295 | 0.524 | 0.288 | **0.512** | 0.409 | **0.692** | **0.884** | 0.665 | 0.665 | 0.000 | 0.295 | 49 | 52 |
| scLM | CellWhisperer | C | 0.292 | 0.364 | 0.242 | 0.314 | 0.335 | 0.528 | 0.787 | 0.611 | 0.611 | 0.000 | 0.292 | 33 | 33 |
| scLM | C2S-Scale 2B | U | 0.239 | 0.408 | 0.245 | 0.415 | 0.355 | 0.599 | 0.827 | 0.604 | 0.603 | 0.001 | 0.239 | 55 | 58 |
| baseline | SingleR | C | 0.236 | 0.366 | 0.145 | 0.355 | 0.423 | 0.514 | 0.759 | 0.444 | 0.443 | 0.000 | 0.236 | 12 | 12 |
| baseline | scType | C | 0.222 | 0.336 | 0.122 | 0.302 | 0.397 | 0.517 | 0.792 | 0.500 | 0.500 | 0.000 | 0.222 | 10 | 10 |
| baseline | SingleR | U | 0.174 | 0.332 | 0.135 | 0.334 | 0.330 | 0.534 | 0.781 | 0.522 | 0.522 | 0.000 | 0.174 | 27 | 27 |
| baseline | scType | U | 0.160 | 0.328 | 0.112 | 0.301 | 0.301 | 0.549 | 0.807 | 0.538 | 0.472 | 1.000 | 0.000 | 12 | 12 |
| baseline | SCSA | C | 0.158 | 0.067 | 0.064 | 0.040 | 0.291 | 0.317 | 0.657 | 0.428 | 0.428 | 0.000 | 0.158 | 10 | 10 |
| scLM | C2S-Scale 27B | U | 0.154 | 0.138 | 0.132 | 0.196 | 0.324 | 0.358 | 0.615 | 0.427 | 0.407 | 0.115 | 0.152 | 97 | 214 |
| scLM | mmContext | U | 0.136 | 0.198 | 0.151 | 0.270 | 0.232 | 0.396 | 0.688 | 0.523 | 0.520 | 0.000 | 0.136 | 296 | 299 |
| baseline | CellTypist | C | 0.104 | 0.009 | 0.006 | 0.001 | 0.352 | 0.205 | 0.518 | 0.188 | 0.188 | 0.000 | 0.104 | 4 | 4 |
| scLM | CellWhisperer | U | 0.101 | 0.253 | 0.093 | 0.237 | 0.170 | 0.429 | 0.719 | 0.474 | 0.472 | 0.000 | 0.101 | 125 | 126 |
| baseline | CellTypist | U | 0.100 | 0.009 | 0.048 | 0.005 | **0.464** | 0.246 | 0.611 | 0.345 | 0.276 | 1.000 | 0.000 | 14 | 16 |
| scLM | C2S-Scale 27B | C | 0.094 | 0.053 | 0.076 | 0.074 | 0.193 | 0.278 | 0.560 | 0.365 | 0.360 | 0.049 | 0.094 | 89 | 109 |
| chance | Random Stratified | – | 0.030 | 0.103 | 0.030 | 0.103 | 0.046 | 0.233 | 0.456 | 0.184 | 0.185 | 0.000 | 0.030 | 38 | 38 |
| baseline | SCSA | U | 0.028 | 0.004 | 0.001 | 0.000 | 0.306 | 0.289 | 0.666 | 0.443 | 0.365 | 0.479 | 0.021 | 10 | 10 |
| scLM | InstructCell | C | 0.024 | 0.002 | 0.000 | 0.000 | 0.315 | 0.231 | 0.578 | 0.250 | 0.238 | 0.119 | 0.024 | 9 | 40 |
| scLM | InstructCell | U | 0.020 | 0.002 | 0.000 | 0.000 | 0.308 | 0.230 | 0.578 | 0.251 | 0.241 | 0.137 | 0.020 | 9 | 46 |

Table S6. **Benchmark results for HIHA.** One row per method and setting, 43 at 1,000 cells and 20 at 10,000 cells, sorted by balanced accuracy within each size. The general-purpose LLMs and LangCell (U) ran at 1,000 cells only. Columns, abbreviations and bold type as in Supplementary Table S5. Every answer was a Cell Ontology term or had a reviewed mapping.

| | | | Exact match | | | | Ontology-aware | | | Text similarity | | Curated mapping | | Distinct labels | |
|---|---|---|---|---|---|---|---|---|---|---|---|---|---|---|---|
| Family | Method | Setting | Bal. acc. | Acc. | Macro F1 | Wtd. F1 | Agree. | Rada | Lin | Mapped | Raw | Needed | Bal. acc. without | Mapped | Raw |
| **1,000 cells** | | | | | | | | | | | | | | | |
| LLM | GPT-5.5 | C | **0.597** | **0.714** | **0.574** | **0.682** | **0.597** | **0.789** | 0.813 | **0.767** | **0.756** | 0.000 | 0.597 | 29 | 29 |
| LLM | Claude Opus 4.7 | C | 0.585 | 0.684 | 0.524 | 0.646 | 0.586 | 0.768 | 0.801 | 0.753 | 0.744 | 0.001 | 0.585 | 27 | 28 |
| LLM | Gemini 3.1 Pro | C | 0.570 | 0.700 | 0.548 | 0.665 | 0.572 | 0.780 | **0.814** | 0.751 | 0.743 | 0.000 | 0.570 | 29 | 29 |
| LLM | Claude Sonnet 4.6 | C | 0.532 | 0.642 | 0.497 | 0.605 | 0.534 | 0.739 | 0.774 | 0.728 | 0.720 | 0.000 | 0.532 | 26 | 26 |
| LLM | Kimi-K2.6 | C | 0.462 | 0.543 | 0.457 | 0.534 | 0.462 | 0.663 | 0.668 | 0.632 | 0.626 | 0.122 | 0.462 | 24 | 25 |
| LLM | DeepSeek-V3.2 | C | 0.462 | 0.593 | 0.424 | 0.540 | 0.463 | 0.695 | 0.744 | 0.670 | 0.665 | 0.001 | 0.462 | 26 | 27 |
| LLM | GPT-OSS-120B | C | 0.445 | 0.605 | 0.425 | 0.542 | 0.445 | 0.704 | 0.751 | 0.652 | 0.646 | 0.003 | 0.445 | 24 | 25 |
| LLM | GPT-OSS-20B | C | 0.404 | 0.561 | 0.387 | 0.498 | 0.404 | 0.673 | 0.720 | 0.636 | 0.629 | 0.001 | 0.402 | 26 | 27 |
| LLM | GPT-5.5 | U | 0.348 | 0.470 | 0.367 | 0.485 | 0.435 | 0.633 | 0.741 | 0.665 | 0.660 | 0.115 | 0.328 | 94 | 109 |
| LLM | Llama-4-Maverick | C | 0.340 | 0.519 | 0.312 | 0.454 | 0.341 | 0.641 | 0.693 | 0.593 | 0.589 | 0.002 | 0.340 | 22 | 24 |
| LLM | Gemini 3.1 Pro | U | 0.326 | 0.509 | 0.313 | 0.495 | 0.413 | 0.671 | 0.757 | 0.657 | 0.654 | 0.032 | 0.310 | 39 | 44 |
| LLM | Claude Opus 4.7 | U | 0.274 | 0.452 | 0.290 | 0.472 | 0.375 | 0.628 | 0.734 | 0.638 | 0.636 | 0.096 | 0.263 | 70 | 86 |
| scLM | mmContext | C | 0.263 | 0.326 | 0.202 | 0.292 | 0.263 | 0.475 | 0.523 | 0.526 | 0.519 | 0.000 | 0.263 | 28 | 28 |
| LLM | Claude Sonnet 4.6 | U | 0.261 | 0.434 | 0.263 | 0.419 | 0.361 | 0.612 | 0.735 | 0.609 | 0.605 | 0.101 | 0.249 | 52 | 71 |
| baseline | scType | C | 0.258 | 0.517 | 0.186 | 0.439 | 0.276 | 0.635 | 0.669 | 0.548 | 0.547 | 0.000 | 0.258 | 10 | 10 |
| scLM | CellWhisperer | C | 0.257 | 0.363 | 0.177 | 0.280 | 0.257 | 0.520 | 0.585 | 0.520 | 0.517 | 0.000 | 0.257 | 17 | 17 |
| baseline | SCSA | C | 0.247 | 0.477 | 0.163 | 0.370 | 0.247 | 0.589 | 0.638 | 0.479 | 0.478 | 0.000 | 0.247 | 8 | 8 |
| baseline | SingleR | C | 0.226 | 0.493 | 0.157 | 0.369 | 0.226 | 0.624 | 0.694 | 0.500 | 0.493 | 0.000 | 0.226 | 7 | 7 |
| LLM | DeepSeek-V3.2 | U | 0.225 | 0.394 | 0.229 | 0.375 | 0.290 | 0.572 | 0.670 | 0.547 | 0.551 | 0.085 | 0.216 | 69 | 83 |
| LLM | Llama-3.3-70B | C | 0.220 | 0.390 | 0.178 | 0.277 | 0.221 | 0.525 | 0.609 | 0.487 | 0.483 | 0.004 | 0.215 | 20 | 22 |
| baseline | scType | U | 0.213 | 0.458 | 0.199 | 0.458 | 0.247 | 0.628 | 0.669 | 0.536 | 0.509 | 1.000 | 0.000 | 16 | 16 |
| LLM | Llama-3.1-70B | C | 0.200 | 0.336 | 0.152 | 0.239 | 0.200 | 0.476 | 0.547 | 0.455 | 0.450 | 0.008 | 0.198 | 23 | 31 |
| LLM | GPT-OSS-120B | U | 0.199 | 0.356 | 0.209 | 0.331 | 0.296 | 0.549 | 0.677 | 0.547 | 0.543 | 0.237 | 0.146 | 58 | 84 |
| LLM | GPT-OSS-20B | U | 0.194 | 0.363 | 0.207 | 0.357 | 0.271 | 0.551 | 0.652 | 0.519 | 0.513 | 0.218 | 0.147 | 80 | 142 |
| scLM | LangCell | C | 0.192 | 0.284 | 0.198 | 0.326 | 0.194 | 0.458 | 0.538 | 0.463 | 0.482 | 0.000 | 0.192 | 28 | 28 |
| scLM | C2S-Scale 2B | U | 0.191 | 0.348 | 0.174 | 0.321 | 0.233 | 0.579 | 0.661 | 0.508 | 0.509 | 0.017 | 0.191 | 17 | 18 |
| scLM | C2S-Scale 27B | U | 0.190 | 0.332 | 0.174 | 0.309 | 0.227 | 0.557 | 0.630 | 0.508 | 0.508 | 0.011 | 0.190 | 34 | 40 |
| LLM | Llama-4-Maverick | U | 0.189 | 0.297 | 0.178 | 0.267 | 0.237 | 0.489 | 0.589 | 0.481 | 0.483 | 0.161 | 0.176 | 38 | 47 |
| LLM | Kimi-K2.6 | U | 0.188 | 0.308 | 0.194 | 0.308 | 0.251 | 0.505 | 0.510 | 0.418 | 0.416 | 0.371 | 0.176 | 58 | 76 |
| scLM | C2S-Scale 2B | C | 0.188 | 0.340 | 0.170 | 0.309 | 0.227 | 0.573 | 0.661 | 0.508 | 0.509 | 0.009 | 0.188 | 17 | 18 |
| baseline | CellTypist | C | 0.178 | 0.415 | 0.119 | 0.294 | 0.178 | 0.547 | 0.631 | 0.450 | 0.439 | 0.000 | 0.178 | 7 | 7 |
| baseline | SCSA | U | 0.168 | 0.299 | 0.133 | 0.229 | 0.186 | 0.537 | 0.632 | 0.512 | 0.503 | 0.063 | 0.138 | 10 | 11 |
| scLM | C2S-Scale 27B | C | 0.167 | 0.305 | 0.129 | 0.255 | 0.192 | 0.504 | 0.583 | 0.484 | 0.484 | 0.002 | 0.167 | 22 | 24 |
| baseline | SingleR | U | 0.167 | 0.447 | 0.148 | 0.394 | 0.202 | 0.585 | 0.590 | 0.402 | 0.381 | 0.057 | 0.167 | 9 | 10 |
| baseline | CellTypist | U | 0.139 | 0.278 | 0.092 | 0.189 | 0.189 | 0.450 | 0.600 | 0.461 | 0.000 | 1.000 | 0.000 | 24 | 29 |
| LLM | Llama-3.1-70B | U | 0.097 | 0.201 | 0.090 | 0.170 | 0.131 | 0.471 | 0.555 | 0.424 | 0.423 | 0.024 | 0.097 | 42 | 46 |
| LLM | Llama-3.3-70B | U | 0.087 | 0.189 | 0.092 | 0.171 | 0.142 | 0.470 | 0.558 | 0.435 | 0.435 | 0.004 | 0.085 | 17 | 19 |
| chance | Random Stratified | – | 0.038 | 0.076 | 0.037 | 0.074 | 0.038 | 0.260 | 0.325 | 0.252 | 0.253 | 0.000 | 0.038 | 29 | 29 |
| scLM | mmContext | U | 0.021 | 0.020 | 0.036 | 0.036 | 0.079 | 0.271 | 0.400 | 0.307 | 0.304 | 0.001 | 0.021 | 324 | 332 |
| scLM | InstructCell | C | 0.019 | 0.011 | 0.001 | 0.001 | 0.070 | 0.210 | 0.286 | 0.310 | 0.166 | 0.398 | 0.019 | 17 | 29 |
| scLM | InstructCell | U | 0.014 | 0.008 | 0.001 | 0.001 | 0.064 | 0.204 | 0.247 | 0.271 | 0.088 | 0.537 | 0.014 | 17 | 28 |
| scLM | LangCell | U | 0.006 | 0.007 | 0.011 | 0.013 | 0.062 | 0.256 | 0.379 | 0.284 | 0.284 | 0.004 | 0.006 | 183 | 183 |
| scLM | CellWhisperer | U | 0.006 | 0.005 | 0.011 | 0.009 | 0.099 | 0.302 | 0.433 | 0.335 | 0.335 | 0.000 | 0.006 | 95 | 95 |

*continued on the next page*

| Family | Method | Setting | Exact match: Bal. acc. | Exact match: Acc. | Exact match: Macro F1 | Exact match: Wtd. F1 | Ontology-aware: Agree. | Ontology-aware: Rada | Ontology-aware: Lin | Text similarity: Mapped | Text similarity: Raw | Curated mapping: Needed | Curated mapping: Bal. acc. without | Distinct labels: Mapped | Distinct labels: Raw |
|---|---|---|---|---|---|---|---|---|---|---|---|---|---|---|---|
| **10,000 cells** | | | | | | | | | | | | | | | |
| scLM | mmContext | C | **0.274** | 0.358 | 0.181 | 0.353 | 0.277 | 0.494 | 0.526 | 0.542 | 0.533 | 0.000 | 0.274 | 29 | 29 |
| baseline | scType | C | 0.262 | **0.675** | 0.186 | 0.618 | **0.279** | **0.756** | 0.766 | **0.549** | **0.549** | 0.000 | 0.262 | 10 | 10 |
| scLM | CellWhisperer | C | 0.253 | 0.424 | 0.159 | 0.358 | 0.253 | 0.569 | 0.621 | 0.519 | 0.517 | 0.000 | 0.253 | 18 | 18 |
| baseline | SCSA | C | 0.249 | 0.642 | 0.182 | 0.570 | 0.249 | 0.723 | 0.752 | 0.480 | 0.479 | 0.000 | 0.249 | 8 | 8 |
| baseline | SingleR | C | 0.224 | 0.651 | 0.183 | 0.559 | 0.224 | 0.746 | **0.778** | 0.499 | 0.492 | 0.000 | 0.224 | 7 | 7 |
| baseline | scType | U | 0.214 | 0.611 | **0.206** | **0.624** | 0.248 | 0.744 | 0.766 | 0.536 | 0.510 | 1.000 | 0.000 | 16 | 16 |
| scLM | C2S-Scale 2B | U | 0.203 | 0.434 | 0.198 | 0.440 | 0.246 | 0.653 | 0.709 | 0.517 | 0.518 | 0.017 | 0.203 | 24 | 25 |
| scLM | C2S-Scale 2B | C | 0.200 | 0.416 | 0.195 | 0.413 | 0.239 | 0.643 | 0.705 | 0.511 | 0.512 | 0.009 | 0.200 | 20 | 21 |
| scLM | LangCell | C | 0.194 | 0.323 | 0.191 | 0.395 | 0.196 | 0.489 | 0.560 | 0.463 | 0.487 | 0.000 | 0.194 | 28 | 28 |
| scLM | C2S-Scale 27B | U | 0.193 | 0.412 | 0.175 | 0.416 | 0.227 | 0.626 | 0.679 | 0.509 | 0.510 | 0.006 | 0.193 | 62 | 80 |
| baseline | CellTypist | C | 0.183 | 0.561 | 0.146 | 0.468 | 0.183 | 0.662 | 0.705 | 0.459 | 0.447 | 0.000 | 0.183 | 7 | 7 |
| scLM | C2S-Scale 27B | C | 0.178 | 0.385 | 0.140 | 0.344 | 0.198 | 0.564 | 0.620 | 0.486 | 0.486 | 0.002 | 0.178 | 35 | 41 |
| baseline | SCSA | U | 0.168 | 0.387 | 0.149 | 0.341 | 0.185 | 0.626 | 0.700 | 0.514 | 0.506 | 0.070 | 0.138 | 10 | 11 |
| baseline | SingleR | U | 0.166 | 0.610 | 0.158 | 0.569 | 0.199 | 0.718 | 0.725 | 0.397 | 0.375 | 0.045 | 0.166 | 9 | 10 |
| baseline | CellTypist | U | 0.138 | 0.365 | 0.110 | 0.301 | 0.190 | 0.526 | 0.660 | 0.459 | 0.000 | 1.000 | 0.000 | 29 | 40 |
| chance | Random Stratified | – | 0.034 | 0.118 | 0.033 | 0.117 | 0.034 | 0.297 | 0.366 | 0.256 | 0.257 | 0.000 | 0.034 | 29 | 29 |
| scLM | InstructCell | C | 0.022 | 0.001 | 0.000 | 0.000 | 0.072 | 0.210 | 0.310 | 0.308 | 0.165 | 0.321 | 0.022 | 23 | 87 |
| scLM | mmContext | U | 0.021 | 0.012 | 0.034 | 0.022 | 0.087 | 0.277 | 0.401 | 0.310 | 0.305 | 0.001 | 0.021 | 958 | 979 |
| scLM | InstructCell | U | 0.012 | 0.001 | 0.000 | 0.000 | 0.060 | 0.205 | 0.273 | 0.269 | 0.088 | 0.441 | 0.012 | 24 | 76 |
| scLM | CellWhisperer | U | 0.008 | 0.006 | 0.014 | 0.010 | 0.110 | 0.330 | 0.476 | 0.333 | 0.333 | 0.001 | 0.008 | 200 | 203 |

Table S7. **Benchmark results for Zeisel.** One row per method and setting, 43 at 1,000 cells and 20 at 10,000 cells, sorted by balanced accuracy within each size. The general-purpose LLMs and LangCell (U) ran at 1,000 cells only. Columns, abbreviations and bold type as in Supplementary Table S5. Every answer was a Cell Ontology term or had a reviewed mapping.

| | | | Exact match | | | | Ontology-aware | | | Text similarity | | Curated mapping | | Distinct labels | |
|---|---|---|---|---|---|---|---|---|---|---|---|---|---|---|---|
| Family | Method | Setting | Bal. acc. | Acc. | Macro F1 | Wtd. F1 | Agree. | Rada | Lin | Mapped | Raw | Needed | Bal. acc. without | Mapped | Raw |
| **1,000 cells** | | | | | | | | | | | | | | | |
| LLM | GPT-5.5 | C | **0.937** | **0.938** | **0.939** | **0.938** | **0.937** | **0.954** | **0.980** | **0.975** | 0.777 | 0.000 | 0.937 | 11 | 11 |
| LLM | Gemini 3.1 Pro | C | 0.931 | 0.934 | 0.930 | 0.934 | 0.931 | 0.951 | 0.978 | 0.971 | 0.777 | 0.000 | 0.931 | 11 | 11 |
| LLM | Claude Opus 4.7 | C | 0.921 | 0.918 | 0.919 | 0.921 | 0.921 | 0.941 | 0.974 | 0.964 | 0.760 | 0.000 | 0.921 | 11 | 11 |
| LLM | Claude Sonnet 4.6 | C | 0.921 | 0.929 | 0.922 | 0.929 | 0.921 | 0.947 | 0.975 | 0.963 | 0.762 | 0.000 | 0.921 | 11 | 11 |
| LLM | GPT-OSS-120B | C | 0.904 | 0.920 | 0.911 | 0.920 | 0.904 | 0.940 | 0.972 | 0.952 | 0.759 | 0.000 | 0.904 | 11 | 11 |
| LLM | DeepSeek-V3.2 | C | 0.877 | 0.896 | 0.882 | 0.894 | 0.877 | 0.921 | 0.956 | 0.932 | 0.730 | 0.000 | 0.877 | 11 | 11 |
| LLM | GPT-OSS-20B | C | 0.863 | 0.892 | 0.881 | 0.892 | 0.863 | 0.920 | 0.959 | 0.931 | 0.755 | 0.002 | 0.863 | 11 | 12 |
| LLM | Kimi-K2.6 | C | 0.850 | 0.894 | 0.874 | 0.893 | 0.850 | 0.920 | 0.954 | 0.911 | 0.711 | 0.006 | 0.847 | 11 | 13 |
| LLM | Gemini 3.1 Pro | U | 0.824 | 0.851 | 0.846 | 0.869 | 0.853 | 0.897 | 0.938 | 0.892 | **0.890** | 0.029 | 0.815 | 27 | 33 |
| LLM | GPT-5.5 | U | 0.781 | 0.847 | 0.825 | 0.869 | 0.836 | 0.895 | 0.942 | 0.886 | 0.885 | 0.042 | 0.766 | 31 | 37 |
| LLM | Llama-4-Maverick | C | 0.757 | 0.753 | 0.777 | 0.755 | 0.757 | 0.827 | 0.931 | 0.902 | 0.710 | 0.000 | 0.757 | 11 | 11 |
| LLM | Claude Sonnet 4.6 | U | 0.755 | 0.813 | 0.791 | 0.824 | 0.795 | 0.866 | 0.919 | 0.848 | 0.835 | 0.074 | 0.643 | 39 | 51 |
| LLM | Claude Opus 4.7 | U | 0.723 | 0.795 | 0.753 | 0.817 | 0.790 | 0.861 | 0.926 | 0.838 | 0.823 | 0.170 | 0.653 | 28 | 41 |
| baseline | SCSA | C | 0.717 | 0.839 | 0.648 | 0.786 | 0.717 | 0.882 | 0.954 | 0.871 | 0.735 | 0.000 | 0.717 | 8 | 8 |
| scLM | InstructCell | C | 0.710 | 0.892 | 0.695 | 0.881 | 0.710 | 0.916 | 0.935 | 0.820 | 0.271 | 0.763 | 0.289 | 13 | 18 |
| scLM | InstructCell | U | 0.705 | 0.888 | 0.686 | 0.878 | 0.705 | 0.912 | 0.927 | 0.803 | 0.269 | 0.765 | 0.289 | 13 | 20 |
| LLM | DeepSeek-V3.2 | U | 0.673 | 0.768 | 0.703 | 0.764 | 0.721 | 0.831 | 0.891 | 0.790 | 0.752 | 0.430 | 0.506 | 43 | 70 |
| LLM | GPT-OSS-120B | U | 0.672 | 0.765 | 0.727 | 0.793 | 0.747 | 0.836 | 0.896 | 0.814 | 0.748 | 0.662 | 0.364 | 40 | 63 |
| LLM | Kimi-K2.6 | U | 0.671 | 0.756 | 0.732 | 0.784 | 0.717 | 0.818 | 0.853 | 0.780 | 0.753 | 0.461 | 0.536 | 43 | 53 |
| scLM | C2S-Scale 2B | C | 0.655 | 0.861 | 0.659 | 0.864 | 0.701 | 0.902 | 0.931 | 0.778 | 0.795 | 0.042 | 0.655 | 12 | 13 |
| baseline | SCSA | U | 0.625 | 0.792 | 0.584 | 0.780 | 0.715 | 0.885 | 0.944 | 0.822 | 0.622 | 0.778 | 0.177 | 9 | 9 |
| LLM | Llama-3.3-70B | C | 0.622 | 0.666 | 0.563 | 0.680 | 0.622 | 0.764 | 0.886 | 0.811 | 0.739 | 0.001 | 0.622 | 10 | 12 |
| baseline | scType | C | 0.617 | 0.800 | 0.559 | 0.770 | 0.617 | 0.845 | 0.853 | 0.732 | 0.605 | 0.088 | 0.617 | 7 | 8 |
| LLM | GPT-OSS-20B | U | 0.599 | 0.717 | 0.668 | 0.751 | 0.672 | 0.803 | 0.869 | 0.776 | 0.720 | 0.675 | 0.302 | 40 | 89 |
| scLM | C2S-Scale 2B | U | 0.598 | 0.790 | 0.658 | 0.820 | 0.675 | 0.850 | 0.891 | 0.784 | 0.790 | 0.033 | 0.598 | 36 | 37 |
| baseline | scType | U | 0.591 | 0.762 | 0.551 | 0.759 | 0.637 | 0.846 | 0.900 | 0.793 | 0.605 | 1.000 | 0.000 | 8 | 8 |
| scLM | C2S-Scale 27B | C | 0.573 | 0.772 | 0.612 | 0.801 | 0.609 | 0.833 | 0.864 | 0.725 | 0.743 | 0.089 | 0.573 | 15 | 18 |
| scLM | C2S-Scale 27B | U | 0.572 | 0.764 | 0.632 | 0.793 | 0.612 | 0.826 | 0.879 | 0.753 | 0.761 | 0.036 | 0.570 | 42 | 45 |
| scLM | CellWhisperer | C | 0.545 | 0.358 | 0.507 | 0.378 | 0.545 | 0.556 | 0.840 | 0.844 | 0.792 | 0.000 | 0.545 | 9 | 9 |
| scLM | mmContext | C | 0.533 | 0.470 | 0.513 | 0.493 | 0.533 | 0.623 | 0.848 | 0.819 | 0.804 | 0.000 | 0.533 | 9 | 9 |
| LLM | Llama-3.1-70B | C | 0.527 | 0.535 | 0.484 | 0.561 | 0.531 | 0.667 | 0.831 | 0.753 | 0.662 | 0.021 | 0.521 | 12 | 19 |
| scLM | LangCell | C | 0.482 | 0.415 | 0.449 | 0.418 | 0.482 | 0.593 | 0.840 | 0.770 | 0.745 | 0.000 | 0.482 | 9 | 9 |
| baseline | CellTypist | U | 0.336 | 0.578 | 0.247 | 0.461 | 0.336 | 0.663 | 0.803 | 0.599 | 0.000 | 0.826 | 0.088 | 4 | 12 |
| baseline | CellTypist | C | 0.336 | 0.578 | 0.247 | 0.461 | 0.336 | 0.663 | 0.803 | 0.599 | 0.390 | 0.000 | 0.336 | 4 | 4 |
| LLM | Llama-4-Maverick | U | 0.307 | 0.435 | 0.335 | 0.477 | 0.408 | 0.594 | 0.732 | 0.654 | 0.669 | 0.305 | 0.247 | 35 | 51 |
| baseline | SingleR | U | 0.295 | 0.490 | 0.208 | 0.420 | 0.295 | 0.603 | 0.783 | 0.590 | 0.590 | 0.000 | 0.295 | 4 | 4 |
| baseline | SingleR | C | 0.295 | 0.490 | 0.208 | 0.420 | 0.295 | 0.603 | 0.783 | 0.590 | 0.455 | 0.000 | 0.295 | 4 | 4 |
| LLM | Llama-3.3-70B | U | 0.294 | 0.281 | 0.287 | 0.280 | 0.341 | 0.474 | 0.643 | 0.580 | 0.548 | 0.490 | 0.241 | 33 | 47 |
| LLM | Llama-3.1-70B | U | 0.265 | 0.289 | 0.307 | 0.352 | 0.336 | 0.484 | 0.663 | 0.582 | 0.550 | 0.484 | 0.200 | 58 | 80 |
| scLM | LangCell | U | 0.114 | 0.131 | 0.107 | 0.173 | 0.232 | 0.388 | 0.690 | 0.528 | 0.528 | 0.000 | 0.114 | 94 | 95 |
| chance | Random Stratified | – | 0.100 | 0.185 | 0.100 | 0.185 | 0.100 | 0.376 | 0.651 | 0.487 | 0.375 | 0.000 | 0.100 | 11 | 11 |
| scLM | CellWhisperer | U | 0.066 | 0.084 | 0.044 | 0.106 | 0.181 | 0.346 | 0.673 | 0.520 | 0.520 | 0.000 | 0.066 | 84 | 84 |
| scLM | mmContext | U | 0.064 | 0.090 | 0.101 | 0.151 | 0.218 | 0.315 | 0.476 | 0.456 | 0.461 | 0.005 | 0.064 | 108 | 108 |

*continued on the next page*

*human pancreas, continued*

| | | | Exact match | | | | Ontology-aware | | | Text similarity | | Curated mapping | | Distinct labels | |
|---|---|---|---|---|---|---|---|---|---|---|---|---|---|---|---|
| Family | Method | Setting | Bal. acc. | Acc. | Macro F1 | Wtd. F1 | Agree. | Rada | Lin | Mapped | Raw | Needed | Bal. acc. without | Mapped | Raw |
| **10,000 cells** | | | | | | | | | | | | | | | |
| baseline | SCSA | C | **0.718** | 0.872 | 0.669 | 0.829 | **0.718** | 0.904 | **0.966** | **0.873** | 0.735 | 0.000 | 0.718 | 8 | 8 |
| scLM | InstructCell | C | 0.714 | **0.933** | **0.730** | **0.938** | 0.714 | **0.947** | 0.955 | 0.816 | 0.263 | 0.785 | 0.279 | 15 | 44 |
| scLM | InstructCell | U | 0.705 | 0.927 | 0.719 | 0.932 | 0.705 | 0.942 | 0.949 | 0.802 | 0.258 | 0.786 | 0.277 | 19 | 51 |
| scLM | C2S-Scale 2B | C | 0.650 | 0.891 | 0.670 | 0.898 | 0.696 | 0.926 | 0.955 | 0.774 | **0.794** | 0.023 | 0.650 | 16 | 17 |
| baseline | SCSA | U | 0.626 | 0.823 | 0.604 | 0.823 | 0.716 | 0.907 | 0.957 | 0.823 | 0.624 | 0.804 | 0.180 | 9 | 9 |
| baseline | scType | C | 0.610 | 0.839 | 0.582 | 0.822 | 0.610 | 0.877 | 0.890 | 0.732 | 0.608 | 0.082 | 0.610 | 7 | 8 |
| scLM | C2S-Scale 2B | U | 0.602 | 0.827 | 0.665 | 0.852 | 0.671 | 0.877 | 0.907 | 0.781 | 0.789 | 0.028 | 0.602 | 63 | 66 |
| baseline | scType | U | 0.583 | 0.797 | 0.576 | 0.811 | 0.629 | 0.875 | 0.931 | 0.787 | 0.600 | 1.000 | 0.000 | 8 | 8 |
| scLM | C2S-Scale 27B | C | 0.547 | 0.806 | 0.592 | 0.830 | 0.580 | 0.858 | 0.894 | 0.714 | 0.731 | 0.057 | 0.543 | 29 | 37 |
| scLM | C2S-Scale 27B | U | 0.528 | 0.775 | 0.591 | 0.801 | 0.569 | 0.834 | 0.883 | 0.729 | 0.735 | 0.036 | 0.527 | 104 | 124 |
| scLM | CellWhisperer | C | 0.525 | 0.327 | 0.490 | 0.348 | 0.525 | 0.534 | 0.834 | 0.833 | 0.782 | 0.000 | 0.525 | 10 | 10 |
| scLM | mmContext | C | 0.515 | 0.485 | 0.509 | 0.511 | 0.515 | 0.632 | 0.852 | 0.806 | 0.791 | 0.000 | 0.515 | 9 | 9 |
| scLM | LangCell | C | 0.443 | 0.387 | 0.430 | 0.396 | 0.443 | 0.575 | 0.840 | 0.750 | 0.727 | 0.000 | 0.443 | 11 | 11 |
| baseline | CellTypist | U | 0.336 | 0.615 | 0.262 | 0.502 | 0.336 | 0.691 | 0.828 | 0.598 | 0.000 | 0.839 | 0.088 | 4 | 12 |
| baseline | CellTypist | C | 0.336 | 0.615 | 0.262 | 0.502 | 0.336 | 0.691 | 0.828 | 0.598 | 0.380 | 0.000 | 0.336 | 4 | 4 |
| baseline | SingleR | U | 0.298 | 0.529 | 0.218 | 0.461 | 0.298 | 0.633 | 0.809 | 0.590 | 0.590 | 0.000 | 0.298 | 5 | 5 |
| baseline | SingleR | C | 0.298 | 0.529 | 0.218 | 0.461 | 0.298 | 0.633 | 0.809 | 0.590 | 0.461 | 0.000 | 0.298 | 5 | 5 |
| chance | Random Stratified | – | 0.090 | 0.194 | 0.090 | 0.195 | 0.090 | 0.385 | 0.671 | 0.494 | 0.396 | 0.000 | 0.090 | 11 | 11 |
| scLM | mmContext | U | 0.076 | 0.106 | 0.116 | 0.175 | 0.241 | 0.329 | 0.490 | 0.467 | 0.474 | 0.002 | 0.076 | 263 | 263 |
| scLM | CellWhisperer | U | 0.069 | 0.083 | 0.049 | 0.105 | 0.172 | 0.346 | 0.681 | 0.508 | 0.508 | 0.000 | 0.069 | 216 | 217 |



Table S8. **Benchmark results for human pancreas.** One row per method and setting, 43 at 1,000 cells and 20 at 10,000 cells, sorted by balanced accuracy within each size. The general-purpose LLMs and LangCell (U) ran at 1,000 cells only. Columns, abbreviations and bold type as in Supplementary Table S5. Every answer was a Cell Ontology term or had a reviewed mapping.

| Family | Method | HFRA | HIHA | Zeisel | human pancreas | Mean |
|---|---|---|---|---|---|---|
| scLM | InstructCell | 0.605 | 0.446 | 0.497 | 0.076 | 0.406 |
| scLM | C2S-Scale 2B | 0.108 | 0.384 | 0.510 | 0.098 | 0.275 |
| scLM | C2S-Scale 27B | 0.074 | 0.568 | 0.349 | 0.094 | 0.271 |
| LLM | Llama-3.1-70B | 0.001 | 0.059 | 0.017 | 0.009 | 0.021 |
| LLM | Llama-3.3-70B | 0.000 | 0.023 | 0.001 | 0.000 | 0.006 |
| LLM | GPT-OSS-20B | 0.000 | 0.001 | 0.000 | 0.000 | 0.000 |
| LLM | Claude Opus 4.7 | 0.000 | 0.000 | 0.000 | 0.000 | 0.000 |
| LLM | Claude Sonnet 4.6 | 0.000 | 0.000 | 0.000 | 0.000 | 0.000 |
| LLM | DeepSeek-V3.2 | 0.000 | 0.000 | 0.000 | 0.000 | 0.000 |
| LLM | GPT-5.5 | 0.000 | 0.000 | 0.000 | 0.000 | 0.000 |
| LLM | GPT-OSS-120B | 0.000 | 0.000 | 0.000 | 0.000 | 0.000 |
| LLM | Gemini 3.1 Pro | 0.000 | 0.000 | 0.000 | 0.000 | 0.000 |
| LLM | Kimi-K2.6 | 0.000 | 0.000 | 0.000 | 0.000 | 0.000 |
| LLM | Llama-4-Maverick | 0.000 | 0.000 | 0.000 | 0.000 | 0.000 |

**Table S9 Answers outside the list of expected cell types.** One row per generative model, the general-purpose LLMs and the generative scLMs, in its constrained run at 1,000 cells, sorted by the mean. *Family* and *Method* name the method. Each dataset column gives the share of cells whose answer named a cell type that is not on the list, and *Mean* averages the four datasets. Answers are compared with the listed types after the curated mapping, so another name for a listed type counts as listed, and an answer that names no cell type does not count as outside the list. Shares are averaged over cell types, as balanced accuracy is.

| Model | Identifier | Unit | Shuffled order | Variants |
|---|---|---|---|---|
| GPT-OSS-120B | `openai/gpt-oss-120b` | cell | yes | all genes<br>5,000 variable genes |
| Gemma-2-2B | `google/gemma-2-2b` | cell | – | all genes<br>5,000 variable genes |
| C2S-Scale 2B | `vandijklab/C2S-Scale-Gemma-2-2B` | cell | yes | all genes<br>5,000 variable genes |
| GPT-OSS-120B | `openai/gpt-oss-120b` | cluster | – | resolution 0.5<br>resolution 1.0 |
| Gemini 3.1 Pro | `google/gemini-3.1-pro-preview` | cluster | – | resolution 0.5<br>resolution 1.0 |

**Table S10 Arms of the prompt-context ablation.** Each row is one arm. *Identifier* is the model's identifier on OpenRouter for GPT-OSS-120B and Gemini 3.1 Pro and on Hugging Face for Gemma-2-2B and C2S-Scale 2B, and *Unit* is what one model call annotated, a single cell or a cluster. Every arm received genes only, genes + tissue, genes + list and genes + tissue + list, and *Shuffled order* marks the arms that also received the genes of genes only in shuffled order. *Variants* are the gene vocabularies of the per-cell arms and the clustering resolutions of the per-cluster arms, and the per-cell arms ran the variant with 5,000 variable genes in genes only and in genes + tissue + list.

| Model | Unit | Variant | Condition | Bal. acc. | Agree. | Lin | non-CL | >8 words |
|---|---|---|---|---|---|---|---|---|
| GPT-OSS-120B | cell | all genes | genes, shuffled order | 0.258 | 0.360 | 0.637 | 0.346 | 0.000 |
| | | | genes only | 0.306 | 0.409 | 0.653 | 0.365 | 0.000 |
| | | | genes + tissue | 0.409 | 0.484 | 0.709 | 0.441 | 0.000 |
| | | | genes + list | **0.540** | **0.554** | 0.759 | **0.078** | 0.000 |
| | | | genes + tissue + list | 0.535 | 0.549 | 0.751 | 0.079 | 0.000 |
| | | 5,000 variable genes | genes only | 0.316 | 0.412 | 0.671 | 0.406 | 0.000 |
| | | | genes + tissue + list | 0.534 | 0.545 | **0.777** | 0.080 | 0.000 |
| Gemma-2-2B | cell | all genes | genes only | 0.000 | 0.000 | 0.000 | 1.000 | 0.113 |
| | | | genes + tissue | 0.000 | 0.000 | 0.000 | 1.000 | 0.720 |
| | | | genes + list | **0.042** | **0.042** | 0.150 | **0.750** | 0.366 |
| | | | genes + tissue + list | 0.039 | 0.039 | **0.162** | 0.777 | 0.547 |
| | | 5,000 variable genes | genes only | 0.000 | 0.000 | 0.000 | 1.000 | 0.231 |
| | | | genes + tissue + list | 0.037 | 0.037 | 0.147 | 0.789 | 0.595 |
| C2S-Scale 2B | cell | all genes | genes, shuffled order | 0.255 | 0.323 | 0.609 | 0.065 | 0.000 |
| | | | genes only | 0.332 | 0.396 | 0.685 | 0.024 | 0.001 |
| | | | genes + tissue | 0.389 | 0.446 | 0.706 | 0.020 | 0.000 |
| | | | genes + list | 0.401 | 0.455 | **0.711** | 0.019 | 0.000 |
| | | | genes + tissue + list | **0.403** | **0.457** | 0.710 | 0.018 | 0.000 |
| | | 5,000 variable genes | genes only | 0.322 | 0.414 | 0.654 | **0.009** | 0.000 |
| | | | genes + tissue + list | 0.377 | 0.434 | 0.695 | 0.010 | 0.000 |
| GPT-OSS-120B | cluster | resolution 0.5 | genes only | 0.257 | 0.352 | 0.634 | 0.340 | 0.000 |
| | | | genes + tissue | 0.382 | 0.439 | 0.712 | 0.440 | 0.000 |
| | | | genes + list | 0.491 | 0.503 | 0.746 | 0.062 | 0.000 |
| | | | genes + tissue + list | 0.462 | 0.478 | 0.722 | 0.087 | 0.000 |
| | | resolution 1.0 | genes only | 0.256 | 0.363 | 0.637 | 0.315 | 0.000 |
| | | | genes + tissue | 0.402 | 0.483 | 0.722 | 0.393 | 0.000 |
| | | | genes + list | 0.507 | 0.521 | 0.727 | 0.088 | 0.007 |
| | | | genes + tissue + list | **0.524** | **0.539** | **0.748** | **0.055** | 0.000 |
| Gemini 3.1 Pro | cluster | resolution 0.5 | genes only | 0.314 | 0.431 | 0.696 | 0.071 | 0.000 |
| | | | genes + tissue | 0.409 | 0.492 | 0.774 | 0.082 | 0.000 |
| | | | genes + list | 0.517 | 0.531 | 0.797 | **0.054** | 0.000 |
| | | | genes + tissue + list | 0.500 | 0.520 | 0.793 | **0.054** | 0.000 |
| | | resolution 1.0 | genes only | 0.372 | 0.468 | 0.718 | 0.059 | 0.000 |
| | | | genes + tissue | 0.454 | 0.538 | 0.779 | 0.082 | 0.000 |
| | | | genes + list | **0.556** | **0.569** | 0.813 | 0.056 | 0.000 |
| | | | genes + tissue + list | 0.555 | 0.568 | **0.816** | 0.056 | 0.000 |

**Table S11 Results of the prompt-context ablation.** One row per arm, variant and condition. *Model* and *Unit* name the arm, and *Variant* is the gene vocabulary of a per-cell arm or the clustering resolution of a per-cluster arm. Each value is the mean over the four datasets, at 1,000 cells each, of the mean over three seeds per dataset. *Bal. acc.* is balanced accuracy and *Agree.* the agreement score, which gives a cell 1 for its true type, 0.5 for a finer type or for a coarser type with at most 250 terms at or below it in the Cell Ontology and 0 otherwise, both averaged over the cells of each type and then over the types. *Lin* is the Lin similarity between the answer and the true type in the Cell Ontology, averaged over cells. *non-CL* is the share of answers that are, as written, not the name or identifier of a Cell Ontology term, and *>8 words* the share that runs past eight words. Within each arm, the highest balanced accuracy, agreement score and Lin similarity and the lowest non-CL share are highlighted in bold.

| Method | Condition | Bal. acc. | Agree. | Lin | Listed | Coarser | Finer | Unc. missing | Unc. listed |
|---|---|---|---|---|---|---|---|---|---|
| SCSA | exact list | 0.414 | 0.448 | 0.698 | – | 0.062 | 0.007 | – | – |
| SCSA | A1 | 0.373 | 0.448 | 0.690 | – | 0.142 | 0.007 | – | – |
| SCSA | A2 (2 types) | 0.401 | 0.435 | 0.692 | – | 0.062 | 0.007 | – | – |
| SCSA | A2 | 0.396 | 0.431 | 0.692 | – | 0.062 | 0.007 | – | – |
| SCSA | A3 | 0.365 | 0.393 | 0.647 | 0.487 | 0.050 | 0.005 | – | – |
| SCSA | A1 + A2 + A3 | 0.281 | 0.395 | 0.638 | 0.377 | 0.221 | 0.007 | – | – |
| scType | exact list | 0.379 | 0.428 | 0.712 | – | 0.093 | 0.003 | – | – |
| scType | A1 | 0.346 | 0.417 | 0.701 | – | 0.124 | 0.018 | – | – |
| scType | A2 (2 types) | 0.374 | 0.422 | 0.708 | – | 0.093 | 0.003 | – | – |
| scType | A2 | 0.369 | 0.417 | 0.704 | – | 0.093 | 0.003 | – | – |
| scType | A3 | 0.342 | 0.379 | 0.652 | 0.457 | 0.068 | 0.006 | – | – |
| scType | A1 + A2 + A3 | 0.309 | 0.400 | 0.657 | 0.413 | 0.164 | 0.018 | – | – |
| CellTypist | exact list | 0.258 | 0.320 | 0.635 | – | 0.124 | 0.000 | – | – |
| CellTypist | A1 | 0.257 | 0.341 | 0.641 | – | 0.168 | 0.000 | – | – |
| CellTypist | A2 (2 types) | 0.257 | 0.319 | 0.634 | – | 0.124 | 0.000 | – | – |
| CellTypist | A2 | 0.256 | 0.318 | 0.635 | – | 0.124 | 0.000 | – | – |
| CellTypist | A3 | 0.205 | 0.247 | 0.583 | 0.272 | 0.083 | 0.001 | – | – |
| CellTypist | A1 + A2 + A3 | 0.200 | 0.277 | 0.583 | 0.265 | 0.154 | 0.002 | – | – |
| SingleR | exact list | 0.313 | 0.360 | 0.716 | – | 0.084 | 0.009 | – | – |
| SingleR | A1 | 0.297 | 0.353 | 0.717 | – | 0.093 | 0.020 | – | – |
| SingleR | A2 (2 types) | 0.312 | 0.359 | 0.715 | – | 0.084 | 0.009 | – | – |
| SingleR | A2 | 0.311 | 0.358 | 0.715 | – | 0.084 | 0.009 | – | – |
| SingleR | A3 | 0.280 | 0.319 | 0.660 | 0.371 | 0.070 | 0.008 | – | – |
| SingleR | A1 + A2 + A3 | 0.259 | 0.337 | 0.668 | 0.343 | 0.135 | 0.020 | – | – |
| C2S-Scale 2B | exact list | 0.402 | 0.456 | 0.711 | – | 0.097 | 0.011 | – | – |
| C2S-Scale 2B | exact list + uncertain | 0.399 | 0.453 | 0.710 | – | 0.096 | 0.011 | – | 0.000 |
| C2S-Scale 2B | A1 | 0.397 | 0.455 | 0.711 | – | 0.103 | 0.012 | – | – |
| C2S-Scale 2B | A2 (2 types) | 0.401 | 0.457 | 0.711 | – | 0.098 | 0.013 | – | – |
| C2S-Scale 2B | A2 | 0.408 | 0.466 | 0.714 | – | 0.101 | 0.014 | – | – |
| C2S-Scale 2B | A3 | 0.400 | 0.455 | 0.710 | 0.417 | 0.098 | 0.011 | 0.000 | 0.000 |
| C2S-Scale 2B | A1 + A2 + A3 | 0.408 | 0.470 | 0.717 | 0.423 | 0.113 | 0.013 | 0.000 | 0.000 |
| LangCell | exact list | 0.382 | 0.401 | 0.695 | – | 0.033 | 0.005 | – | – |
| LangCell | A1 | 0.262 | 0.354 | 0.664 | – | 0.119 | 0.066 | – | – |
| LangCell | A2 (2 types) | 0.370 | 0.389 | 0.681 | – | 0.033 | 0.005 | – | – |
| LangCell | A2 | 0.326 | 0.346 | 0.655 | – | 0.034 | 0.005 | – | – |
| LangCell | A3 | 0.323 | 0.345 | 0.655 | 0.431 | 0.035 | 0.009 | – | – |
| LangCell | A1 + A2 + A3 | 0.178 | 0.273 | 0.617 | 0.241 | 0.117 | 0.072 | – | – |
| mmContext | exact list | 0.372 | 0.387 | 0.656 | – | 0.010 | 0.019 | – | – |
| mmContext | A1 | 0.286 | 0.367 | 0.612 | – | 0.123 | 0.038 | – | – |
| mmContext | A2 (2 types) | 0.365 | 0.380 | 0.653 | – | 0.010 | 0.019 | – | – |
| mmContext | A2 | 0.352 | 0.367 | 0.647 | – | 0.010 | 0.019 | – | – |
| mmContext | A3 | 0.304 | 0.319 | 0.619 | 0.408 | 0.009 | 0.021 | – | – |
| mmContext | A1 + A2 + A3 | 0.208 | 0.298 | 0.577 | 0.279 | 0.140 | 0.041 | – | – |
| Gemini 3.1 Pro | exact list | 0.552 | 0.565 | 0.813 | – | 0.007 | 0.019 | – | – |
| Gemini 3.1 Pro | exact list + uncertain | 0.553 | 0.566 | 0.808 | – | 0.007 | 0.019 | – | 0.016 |
| Gemini 3.1 Pro | A1 | 0.497 | 0.555 | 0.805 | – | 0.090 | 0.026 | – | – |
| Gemini 3.1 Pro | A2 (2 types) | 0.554 | 0.567 | 0.812 | – | 0.007 | 0.019 | – | – |
| Gemini 3.1 Pro | A2 | 0.555 | 0.571 | 0.812 | – | 0.015 | 0.019 | – | – |
| Gemini 3.1 Pro | A3 | 0.499 | 0.512 | 0.720 | 0.665 | 0.005 | 0.021 | 0.211 | 0.027 |
| Gemini 3.1 Pro | A1 + A2 + A3 | 0.396 | 0.496 | 0.738 | 0.525 | 0.171 | 0.029 | 0.008 | 0.007 |
| GPT-OSS-120B | exact list | 0.551 | 0.564 | 0.763 | – | 0.015 | 0.012 | – | – |
| GPT-OSS-120B | exact list + uncertain | 0.542 | 0.557 | 0.759 | – | 0.017 | 0.012 | – | 0.003 |
| GPT-OSS-120B | A1 | 0.490 | 0.539 | 0.753 | – | 0.063 | 0.036 | – | – |
| GPT-OSS-120B | A2 (2 types) | 0.531 | 0.544 | 0.754 | – | 0.014 | 0.013 | – | – |
| GPT-OSS-120B | A2 | 0.528 | 0.542 | 0.758 | – | 0.016 | 0.013 | – | – |
| GPT-OSS-120B | A3 | 0.436 | 0.449 | 0.700 | 0.584 | 0.014 | 0.012 | 0.152 | 0.007 |

| | | | | | | | | | |
|---|---|---|---|---|---|---|---|---|---|
| GPT-OSS-120B | A1 + A2 + A3 | 0.359 | 0.449 | 0.705 | 0.481 | 0.141 | 0.039 | 0.043 | 0.002 |

Table S12. **Results of the imperfect-list analysis.** One row per method and condition. The conditions are the exact list, the exact list with the answer uncertain (exact list + uncertain), a parent or a child added for half of the types (A1), two or five types absent from the data added (A2 (2 types) or A2), a quarter of the types removed (A3) and all three errors (A1 + A2 + A3). Each value is the mean over the four datasets, at 1,000 cells each, of the mean over three seeds per dataset. *Bal. acc.* is balanced accuracy and *Agree.* the agreement score, which gives a cell 1 for its true type, 0.5 for a finer type or for a coarser type with at most 250 terms at or below it in the Cell Ontology and 0 otherwise, both averaged over the cells of each type and then over all true types, including those removed from the list. *Lin* is the Lin similarity between the answer and the true type in the Cell Ontology, averaged over cells. *Listed* is balanced accuracy over the types still on the list, given where types were removed. *Coarser* and *Finer* are the shares of cells answered with a coarser type that has at most 250 terms at or below it and with a finer type. *Unc. missing* and *Unc. listed* are the shares of the cells of the missing and of the listed types that were answered uncertain, given where that answer was offered. Shares are averaged over cell types. Gemini 3.1 Pro, scType and SCSA annotated clusters, the other methods single cells.
